# Beyond dual hubs: Task and aging shape taxonomic and thematic semantic relationships in the human brain

**DOI:** 10.64898/2026.02.19.706838

**Authors:** Philipp Kuhnke, Sandra Martin, Curtiss A. Chapman, Gesa Hartwigsen

**Author notes:** Leipzig University, Neumarkt 9, 04109 Leipzig, Germany. Shared first authors (Philipp Kuhnke, Sandra Martin).

## Abstract

Semantic knowledge about concepts and their relationships is central to human cognition. Taxonomic relationships link concepts belonging to the same category (e.g. *dog* and *bear*), while thematic relationships connect concepts co-occurring in the same events (e.g. *dog* and *leash*). The dual-hub theory proposes that taxonomic relations preferentially engage the anterior temporal lobe (ATL), whereas thematic relations preferentially involve the temporo-parietal cortex (TPC). However, it remains unclear whether taxonomic and thematic representations depend on the concurrent task and how they change with aging. The present fMRI study addressed these gaps by jointly investigating the effects of semantic relationship, task, and age on semantic processing. 41 young and 34 older adults performed taxonomic and thematic judgments on picture pairs that were taxonomically related, thematically related, or unrelated. Our results do not support the dual-hub theory: Both TPC and ATL preferentially responded to thematic over taxonomic relationships. Moreover, their activity was task-dependent. Semantic control regions flexibly responded to task-relevant semantic relations, while inhibiting task-irrelevant relations. Finally, aging was associated with a decline in domain-general and domain-specific semantic control, more bilateral ATL engagement, as well as behavioral and neural shifts from taxonomic towards thematic processing. Increased thematic activity was associated with higher accuracy but slower responses. These findings support accounts of age-related neural dedifferentiation and semanticization: Older adults require increased cognitive resources to maintain accuracy at a high level, but this comes at the cost of efficiency.

## Introduction

Semantic knowledge – our understanding of concepts and their relationships – supports fundamental cognitive functions such as categorization, prediction, and goal-directed behavior (Lambon Ralph, 2013; Reilly et al., 2024). Concepts can have two distinct types of relations (Davis and Yee, 2019): *Taxonomic relations* link concepts that belong to the same semantic category based on shared features (e.g. *dog* and *bear*), whereas *thematic relations* connect concepts that co-occur or interact within the same situations or events (e.g. *dog* and *leash*). Behavioral and neural evidence suggests that taxonomic and thematic relationships rely on partly dissociable cognitive processes (for a review, see Mirman et al., 2017). However, it remains unclear to what extent taxonomic and thematic representations in the human brain are modulated by task demands and aging.

The “dual-hub theory” (Davis and Yee, 2019; Mirman et al., 2017) proposes that taxonomic and thematic relations depend on separate semantic “hubs” (Kuhnke et al., 2023a; Lambon Ralph et al., 2016): Taxonomic relations are assumed to predominantly engage the anterior temporal lobe (ATL), whereas thematic relations preferentially engage the temporo-parietal cortex (TPC) including parts of angular gyrus (AG) and posterior temporal lobe. This account is supported by lesion studies linking ATL damage with taxonomic deficits (Patterson et al., 2007; Rogers et al., 2004) and TPC damage with thematic deficits (Schwartz et al., 2011; Tsagkaridis et al., 2014). Neuroimaging studies also frequently report preferential ATL activation for taxonomic relations (Geng and Schnur, 2016; Lewis et al., 2015) and TPC activation for thematic relations (de Zubicaray et al., 2013; Henseler et al., 2014; Kalénine et al., 2009).

Despite this converging evidence, the dual-hub theory remains controversial. Contrary to the dual-hub view, ATL damage can be associated with thematic impairments (Riccardi et al., 2024) and TPC lesions can be associated with taxonomic deficits (Semenza et al., 1992). fMRI studies also often show overlapping activations for taxonomic and thematic processing in the ATL and TPC (Jackson et al., 2015; Zhang et al., 2023). A meta-analysis of 16 fMRI studies (Zhang et al., 2023) found that thematic relations indeed preferentially engage the left TPC, but the ATL is not consistently engaged for taxonomic relations; left lateral ATL even shows a preference for thematic relations.

Moreover, it is currently unclear to what extent brain activity for taxonomic and thematic relationships depends on the concurrent task. This is a crucial issue since previous evidence suggests that the retrieval of semantic features is task-dependent, such that semantic feature representations are more strongly or even selectively engaged when they are task-relevant (Kemmerer, 2015; Kuhnke et al., 2020b, 2021; Yee and Thompson-Schill, 2016). Therefore, it seems likely that also taxonomic and thematic representations in ATL and TPC are modulated by the task. Moreover, depending on task difficulty, one may also expect task-dependent recruitment of other brain regions. Beyond representational hubs in the ATL and TPC, demanding semantic tasks often recruit “semantic control” areas, such as the left inferior frontal gyrus (IFG) and posterior middle/inferior temporal gyrus (pMTG/ITG), which guide the retrieval and/or selection of task-relevant semantic information (Shao et al., 2026; Zhang et al., 2021), while inhibiting task-irrelevant information (Jackson, 2021; Jefferies, 2013; Martin et al., 2023a, 2025; Nieberlein et al., 2024). Thus, semantic control regions may be engaged for the retrieval of task-relevant taxonomic or thematic relationships, as well as for the inhibition of task-irrelevant relations. However, a potential task dependency of taxonomic and thematic representations in the human brain has not been systematically tested yet.

A further unresolved issue concerns how taxonomic and thematic processing change with aging. Behaviorally, older adults show reduced use of taxonomic categories and an increased reliance on thematic associations (Smiley and Brown, 1979; Verheyen et al., 2019). Across the lifespan, the pattern follows a U-shaped trajectory—from thematic preference in childhood to taxonomic dominance in young adulthood and a return to thematic bias in older age (Mirman et al., 2017). At the neural level, healthy aging is commonly associated with “neural dedifferentiation”, a reduction in the functional specificity of cortical representations (Koen and Rugg, 2019). Older adults show less distinct activation patterns for object categories in the ventral visual stream (Koen et al., 2020; Srokova et al., 2024), broader activity and decreased segregation of domain-general functional networks (Chan et al., 2014; Martin et al., 2023b, 2022), and reduced separability of semantic categories (Margolles and Soto, 2024). These findings suggest that aging may particularly affect taxonomic representations. However, it remains unclear whether thematic relationships change with age, and age effects on taxonomic and thematic processing have not been examined simultaneously within the same participants.

The present fMRI study addressed these gaps by jointly investigating the effects of relation type, task, and age on semantic processing. We measured the brain activity of 41 young adults and 34 older adults using fMRI, while they judged whether pairs of object drawings were taxonomically related (taxonomic task), or thematically related (thematic task). In both tasks, the drawings could be taxonomically related (e.g., *monkey* and *swan*), thematically related (e.g., *monkey* and *banana*), or unrelated (e.g., *monkey* and *telephone*). A non-semantic visual task on scrambled images served as control.

We first identified commonalities and differences in general semantic processing (semantic > non-semantic tasks) between young and older adults at the whole-brain level. Using subject-specific functional region-of-interest (fROI) analyses, we then investigated age and task effects on the response profiles of key semantic regions—the ATL and TPC, as well as semantic control areas—for taxonomic and thematic relations. Finally, by linking behavioral performance to activity in semantic fROIs, we examined how the behavioral relevance of these regions during the processing of thematic and taxonomic relationships varies with age.

Following the dual-hub theory, we expected that the ATL would show higher activity for taxonomic than thematic relationships, whereas TPC would show higher activity for thematic relations. Semantic control regions (e.g. left IFG, pMTG/ITG) should show increased activation for task-relevant semantic relations, that is, taxonomic pairs in the taxonomic task and thematic pairs in the thematic task. They should also be engaged for the inhibition of task-irrelevant semantic relations, that is, thematic pairs in the taxonomic task and taxonomic pairs in the thematic task. Finally, older adults should show reduced activation for taxonomic processing, but enhanced (or preserved) activation for thematic processing, consistent with behavioral shifts toward thematic reasoning and neural dedifferentiation.

## Materials and Methods

### Participants

41 young adults (20 female; mean age = 27.8 years, SD = 4.32, range = 21–35) and 34 older adults (18 female; mean age = 65.9 years, SD = 3.17, range = 60–70) were included in the final analyses. We exclusively included participants without cognitive impairments according to the Mini Mental State Examination (MMSE; young adults: mean = 29.24, SD = 1.20, range = 26–30; older adults: mean = 29.24; SD = 0.82; range = 28–30). 44 young adults and 42 older adults were originally measured, but 3 young and 8 old adults were excluded due to very low response accuracy (<33%) in at least one experimental condition. All participants were right-handed, native German speakers recruited via the participant database of the Max Planck Institute for Human Cognitive and Brain Sciences, Leipzig, Germany. A neuropsychological test battery was used to assess executive functions, including the Symbol Search, Digit-Symbol Substitution, and Digit Span Forward and Backward tests (Wechsler, 1997). Independent-samples t-tests revealed significant group differences in all tests except Digit Span Forward, with young adults outperforming older adults on the remaining tests (Figure 1B). Written informed consent was obtained from each participant prior to the experiment. The study was performed according to the guidelines of the Declaration of Helsinki and approved by the local ethics committee of the Medical Faculty at Leipzig University.

**Figure 1.**
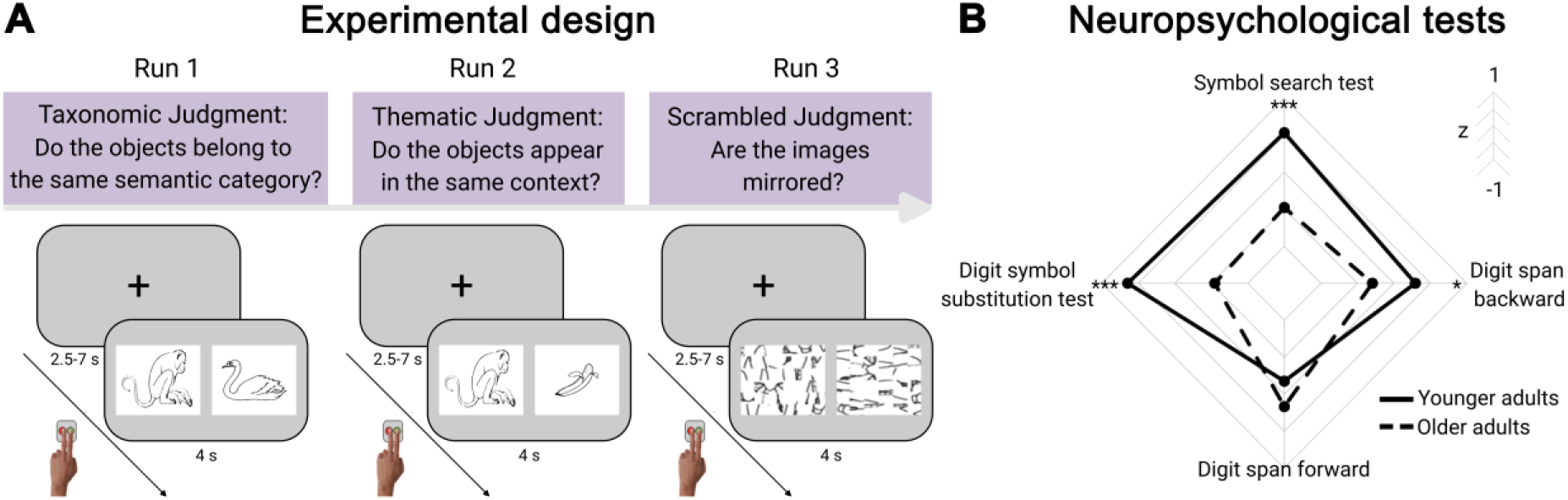
Experimental Design. (A) Participants completed three experimental runs, each with a different task. The task order was counterbalanced across participants. Below each task, an example trial is illustrated. (B) Neuropsychological profile of both age groups. Scores of all tests were z-transformed for comparisons using independent-samples t-tests. ***p < 0.001, *p < 0.05.

### Experimental procedures

Participants performed semantic and non-semantic tasks in separate fMRI runs (order counterbalanced; Figure 1A). The semantic tasks followed a 2 x 2 x 3 factorial design with the factors GROUP (young, older adults), TASK (taxonomic, thematic judgments), and RELATION (taxonomic, thematic, unrelated). Participants were presented with 144 pairs of object images (48 trials per condition), which could be taxonomically related (e.g. *monkey* – *swan*), thematically related (e.g. *monkey* – *banana*), or unrelated (e.g. *monkey* – *star*). During taxonomic judgments, participants had to decide whether the two items belonged to the same taxonomic category. During thematic judgments, they judged whether the two items occurred in the same thematic contexts.

In the non-semantic control task, participants were presented with (unrecognizable) scrambled versions of the same images. They had to decide whether the two images were mirrored versions of each other. Participants performed 96 trials: 48 trials of mirrored images (congruent) and 48 trials of 90° rotated images (incongruent).

On each trial of every task, a pair of images was presented for 4 s, followed by a jittered inter-trial interval (fixation cross) of 2.5 – 7 s (mean 4 s). Participants responded via button press with the index or middle finger of their left hand (counterbalanced across participants).

### Stimuli

Stimuli for the semantic tasks were 144 pairs of black-and-white object drawings (Table S83). Images were selected from several previous studies of taxonomic/thematic processing (Kalénine et al., 2009; Nishimoto et al., 2011; Snodgrass and Vanderwart, 1980), the International Picture Naming Project (Szekely et al., 2004), or hand-drawn by one of the authors (CAC). We balanced image pairings such that the 144 object images appeared three times: once in a taxonomic, once in a thematic, and once in an unrelated pairing with another image. We also aimed to balance the number of stimulus pairs for the object categories of natural vs. artifactual and manipulable vs. non-manipulable items as best as possible between experimental conditions (Table S84).

For the non-semantic control task, scrambled versions of the same pictures were created using the *Telegraphics Scramble* filter in *Adobe Photoshop* (http://telegraphics.com.au/sw/product/Scramble). For each scrambled image, a horizontally mirrored (congruent) and a 90° rotated (incongruent) image were generated.

To assess the individual degree of semantic relatedness for all stimulus pairs, participants completed a rating survey after the fMRI experiment. For each stimulus pair, they rated their semantic relatedness (taxonomic and thematic) on a Likert scale from 1 to 5 (Figure S10A). A linear mixed model (LMM) predicting association ratings with the factors RELATION (taxonomic, thematic, unrelated), JUDGMENT (taxonomic, thematic), and GROUP (young adults, older adults) revealed main effects of RELATION (X = 827.82, p < 0.001) and JUDGMENT (X = 663.55, p < 0.001), as well as an interaction of RELATION x JUDGMENT (X = 15341.77, p < 0.001) but no effect of GROUP (X = 2.61, p = 0.11), indicating similar ratings for young and older adults (Table S85). Across both age groups, taxonomic pairs received higher taxonomic ratings than thematic and unrelated ratings (Figure S10B; Table S86). Thematic pairs received higher thematic ratings than taxonomic and unrelated ratings. These results indicate that participants distinguished taxonomic and thematic relationships in our stimuli as intended.

### Behavioral analyses

We used mixed-effects regression models to analyze accuracy and reaction times in all tasks. To this end, a linear mixed model (LMM) was set up for log-transformed reaction time data and a generalized LMM (GLMM) with a binomial distribution and a logit link function was set up for accuracy data. We calculated separate models for the semantic tasks (taxonomic and thematic judgments) and the scrambled tasks to account for different numbers of conditions. For the semantic tasks, fixed effects included all main effects and two- and three-way interactions of TASK, RELATION, and GROUP, and regressors for task order and gender to account for potential influences. Models further included random intercepts for participants and trials as well as by-participant random slopes for TASK and RELATION. The models for the scrambled task were identical except for the TASK predictor. Models for reaction time and accuracy data were identical and are provided in the Supplementary Material (Tables S1 and S2). Factors were contrast coded using the simple coding scheme, a type of effect coding. Statistical models were run using the *glmmTMB* (Brooks et al., 2017) and *lme4* (Bates et al., 2015) packages in R (version 4.5.0). Post-hoc comparisons were run using the *ggeffects* package (Lüdecke, 2018) and plots were generated using *fmsb* and *ggplot2* (Wickham, 2011).

### fMRI acquisition and preprocessing

fMRI data were collected on a 3T Prisma scanner (Siemens, Erlangen, Germany) equipped with a 32-channel head coil. Functional blood oxygen level dependent (BOLD) images were acquired using a multiband (Feinberg et al., 2010) dual gradient-echo EPI sequence (TR = 2 s; TE = 12 & 33 ms; flip angle = 80°; FoV = 204 mm; voxel size = 2.5 x 2.5 x 2.5 mm; slice gap = 0.25 mm; phase encoding direction = A/P; multiband acceleration factor = 2). 60 slices covering the whole brain were recorded in axial orientation. We used a dual-echo sequence to maximize BOLD sensitivity throughout the entire brain, including in regions susceptible to signal loss in single-echo EPI, particularly the ATL (Halai et al., 2014; Poser et al., 2006). To further reduce susceptibility artifacts, slices were tilted 10° up from the AC-PC line (Weiskopf et al., 2006).

B0 field maps were acquired for susceptibility distortion correction using a gradient-echo sequence (TR = 0.62 s; TE = 4 & 6.46 ms; flip angle = 60°, other parameters identical to functional sequence). Structural T1-weighted images were acquired for normalization using an MPRAGE sequence (176 slices in sagittal orientation; TR = 2.3 s; TE = 2.98 ms; FoV = 256 mm; voxel size = 1 × 1 × 1 mm; no slice gap; flip angle = 9°; phase encoding direction = A/P).

fMRI analysis was performed using *SPM12* (Wellcome Trust Centre for Neuroimaging; http://www.fil.ion.ucl.ac.uk/spm/software/spm12/), implemented in *Matlab* (version 9.10; Mathworks, Natick, MA). The two images of the dual-echo sequence were combined using an average weighted by the temporal signal-to-noise ratio (tSNR) of each image at every voxel, which yields optimal BOLD sensitivity (Poser et al., 2006). Functional images were realigned, distortion-corrected (using the B0 field map), slice-timing corrected, normalized to MNI space (via unified segmentation of the co-registered structural image), and smoothed with a Gaussian kernel of 2x voxel size (FWHM = 5 x 5 x 5.5 mm).

### Whole-brain activation analysis

First, we aimed to assess commonalities and differences in general semantic processing between young and older adults at the whole-brain level. To this end, we performed a random-effects group analysis based on the general linear model (GLM) using the classical two-level approach in SPM.

At the first level, individual participant data were modelled separately. The GLM included regressors for the 8 experimental conditions, modeling trials as boxcar functions (4 s duration) convolved with the canonical HRF. Only correct trials were analyzed; errors and instructions were modelled in separate regressors-of-no-interest. To account for response time differences between trials and conditions, a duration-modulated parametric regressor (duration = RT) was included (Grinband et al., 2008). Nuisance regressors included the six motion parameters and individual regressors for time points with strong volume-to-volume movement (framewise displacement > 0.9; Siegel et al., 2014). The data were subjected to an AR(1) autocorrelation model and high-pass filtered (cutoff 128 s) to remove low-frequency noise.

To identify brain regions involved in general semantic processing, we calculated the contrast of [semantic > non-semantic tasks] within each participant. Only task-relevant trials were included in this contrast (taxonomic task: taxonomic, thematic task: thematic, scrambled task: congruent) to test general semantic retrieval free from additional cognitive demands by task-irrelevant pairings.

At the second (group) level, subject-specific contrast images were entered into non-parametric permutation tests using FSL’s *randomise* function (10,000 permutations; https://web.mit.edu/fsl_v5.0.10/fsl/doc/wiki/Randomise(2f)UserGuide.html). One-sample tests identified significant activation within each age group, and two-sample tests compared activity between groups. Conjunction analysis based on the minimum-statistic (Nichols et al., 2005) identified brain regions commonly engaged in young and older adults. All activation maps were thresholded at p < 0.05 family wise error (FWE) corrected using threshold-free cluster enhancement (TFCE) (Smith and Nichols, 2009).

Finally, we assessed inter-individual consistency in functional activation via threshold-weighted overlap maps (Seghier and Price, 2016), which quantify the percentage of participants activating a particular voxel (or its direct neighbors) weighted by their individual statistical thresholds.

### Functional region-of-interest (fROI) analysis

To investigate age and task effects on the response profiles of semantic brain regions – particularly the bilateral TPC and ATL – for taxonomic and thematic relationships, we performed a functional region-of-interest (fROI) analysis using the group-constrained subject-specific (GSS) approach (Fedorenko et al., 2010; Nieto-Castañón and Fedorenko, 2012). In contrast to standard group analyses that aggregate responses from the same location in standard space, fROI analyses aggregate responses from the same functional region across participants, resulting in higher sensitivity and functional resolution (i.e., the ability to separate adjacent but functionally distinct regions) (Fedorenko and Kanwisher, 2011, 2009).

For fROI definition, we used the general semantic contrast of [semantic > non-semantic tasks (task-relevant conditions only)]. Subject-specific activation maps were thresholded at p < 0.001 and overlaid on top of each other. The resulting overlap map was smoothed (5 mm^3^ FWHM), thresholded at two participants (Julian et al., 2012; Kuhnke et al., 2020b), and parcellated using a watershed algorithm (Meyer, 1991) implemented in the *spm_ss* toolbox (Nieto-Castañón and Fedorenko, 2012). For our primary analysis, we selected parcels in the left and right TPC and ATL, where the highest proportion of participants showed significant activation. For our secondary analysis, we retained 7 other parcels where >70% of participants showed activation. To maximize generalizability to the population, the final analysis included all subjects: fROIs were defined in each individual subject as the 10% most active voxels for the general semantic contrast within each parcel (Basilakos et al., 2018; Jiang et al., 2025).

Subsequently, percent signal change was estimated using the *MarsBaR* toolbox (v0.45; Brett et al., 2002). To avoid circularity, we exclusively compared activity between semantic conditions, independent from the ROI definition contrast (semantic > non-semantic tasks). Statistical inference on each fROI was performed with *JASP* (v0.95.4; https://jasp-stats.org/) using mixed ANOVAs with the factors GROUP (young, older adults), TASK (taxonomic, thematic judgments), and RELATION (taxonomic, thematic, unrelated). Significant interactions were resolved using post-hoc t-tests (Bonferroni-Holm corrected for multiple comparisons).

To relate functional activity to behavior, we examined percent signal change in fROIs that showed a GROUP x TASK or GROUP x RELATION interaction. Because 10 of the 11 fROIs met this criterion, we reduced the number of statistical tests by focusing on task-specific differences: For each fROI, we computed the difference in percent signal change between the two semantic tasks and correlated this value with the corresponding participant-wise differences in reaction time and accuracy. We applied FDR correction to account for two correlations (accuracy and reaction times) per fROI. In addition, we repeated the analysis considering only task-relevant items in the behavioral performance to control for a potential confound of semantic relationship.

## Results

### Behavioral analyses

The GLMM modeling accuracy in the semantic tasks showed significant main effects of GROUP (χ² = 11.44, p < .001), TASK (χ² = 38.21, p < .001), and RELATION (χ² = 62.30, p < .001), as well as significant two-way interactions of TASK x RELATION (χ² = 78.35, p < .001), GROUP and RELATION (χ² = 9.17, p < .010), and a three-way interaction of GROUP x TASK x RELATION (χ² = 49.16, p < .001). Results showed higher accuracy in younger compared to older adults (4% better performance on average), in the taxonomic relative to the thematic task across semantic relationships (4% better performance on average), and overall higher accuracy for thematic than taxonomic relationships (5% higher accuracy on average, see Figure S1). The interaction of TASK x RELATION showed that the difference in accuracy between taxonomic and thematic items was larger in the taxonomic task compared to the thematic task due to the overall better performance in the taxonomic task. The interaction of GROUP x RELATION revealed no difference in accuracy between taxonomic and thematic items in younger adults but better performance for thematic relative to taxonomic items in older adults (see Figure S1). Decomposing the three-way interaction, both groups showed unrelated > thematic > taxonomic accuracy in the taxonomic task, whereas in the thematic task, younger adults were lowest on thematic items and older adults were lowest on taxonomic items (Figure 2B). Table S1 and Figure S1 report all parameter estimates and model results.

**Figure 2.**
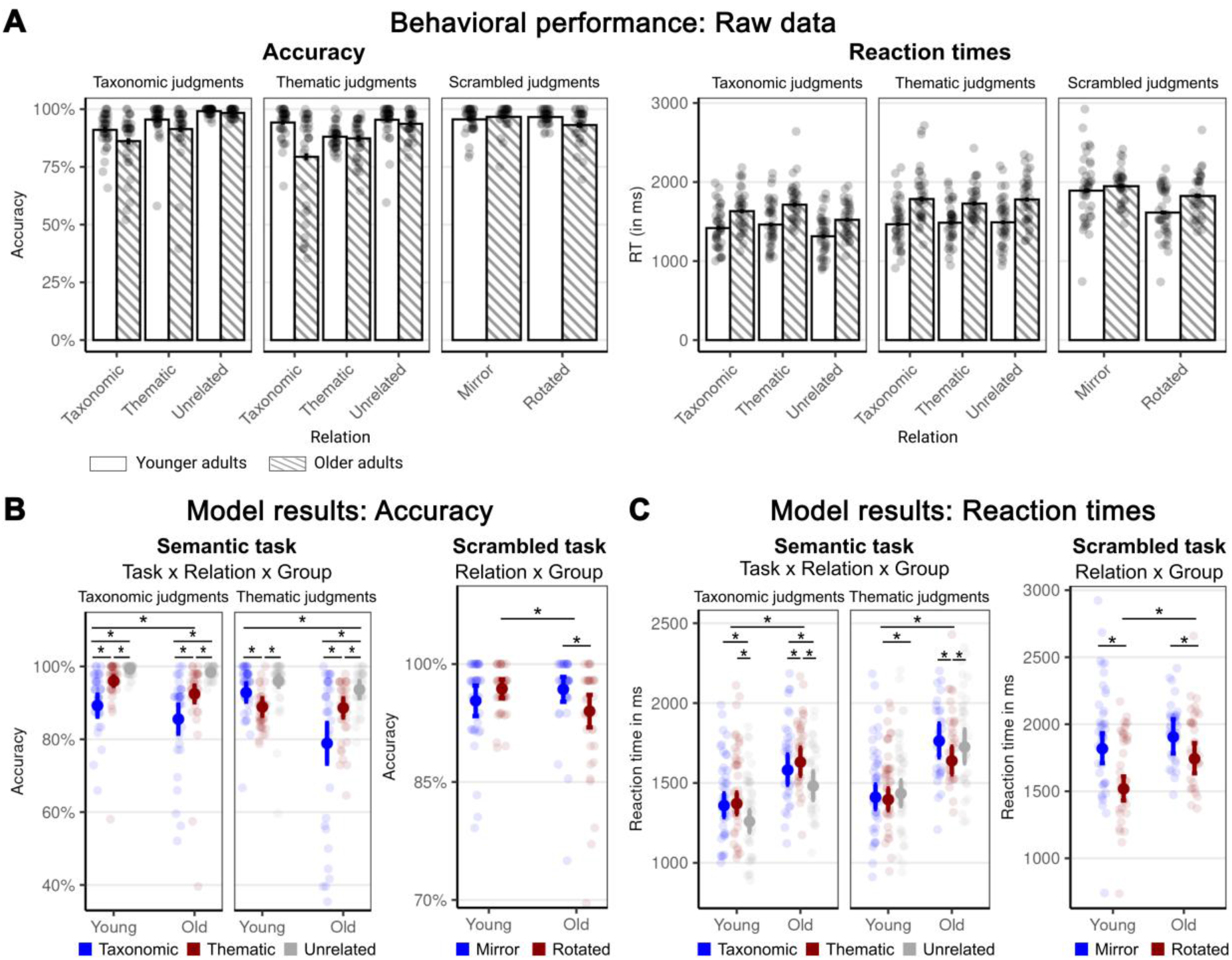
Behavioral results. (A) Raw data for accuracy and reaction times by task, relationship, and age group. Average marginal predictions for accuracy (B) and reaction times (C) for the highest significant interaction terms in the model for semantic tasks (taxonomic and thematic) and the scrambled task. Significant simple contrasts are marked with an asterisk. Full model results are presented in Figure S1 and Tables S1 and S2.

For reaction times in the semantic tasks, the LMM revealed main effects of GROUP (χ² = 25.94, p < .001), TASK (χ² = 39.20, p < .001), and RELATION (χ² = 9.97, p = .007), as well as a significant two-way interaction of TASK x RELATION (χ² = 253.77, p < .001), and a three-way interaction of GROUP x TASK x RELATION (χ² = 25.43, p < .001). Results showed that older adults generally responded slower across tasks and semantic relationships (on average 260 ms slower), reactions were slower for the thematic relative to the taxonomic task (on average 109 ms slower), and semantically unrelated items were generally answered fastest (on average 47 ms faster than other two relationships, see Fig. S1). The interaction of TASK x RELATION showed that semantically unrelated items were fastest in the taxonomic task whereas thematic items were fastest in the thematic task. Task-irrelevant items (taxonomic items in the thematic task and vice versa) were answered slowest in both tasks (all p < .05 after FDR correction, Figure S1). Finally, resolving the three-way interaction showed age-dependent effects of TASK and RELATION. While younger adults showed no difference in reaction times between taxonomic and thematic items in either task, older adults reacted faster to taxonomic items in the taxonomic task and thematic items in the thematic task (Figure 2C). Full model results can be found in Table S1 and Figure S1.

Results from the mixed models for accuracy and reaction times in the scrambled images control task revealed significant interactions of GROUP x RELATION. With respect to accuracy, older and younger adults showed equally high accuracy in the mirror condition, but older adults performed more poorly than younger adults in the rotated condition and more poorly in the rotated than the mirror condition (Figure 2B). For reaction times, both age groups responded more slowly in the rotated than in the mirrored condition, with young adults showing an even greater slowing in the rotated condition than older adults (Figure 2C). Full model results can be found in Table S2.

### Whole-brain activation analysis

To investigate commonalities and differences in general semantic processing between young and older adults, we performed a whole-brain activation analysis, comparing activity for semantic vs. non-semantic tasks (task-relevant conditions only).

In both young and older adults (Figure 3A-C; Tables S3-4), semantic processing engaged the bilateral temporo-parietal cortex (TPC), including parts of posterior middle and inferior temporal gyri (pMTG/ITG) and angular gyrus (AG), as well as bilateral posterior cingulate and precuneus (PCC / PreCun), fusiform gyri (FG), cerebellum, and the left inferior and middle frontal gyrus (IFG / MFG), dorsomedial prefrontal cortex (dmPFC) and anterior temporal lobe (ATL).

**Figure 3.**
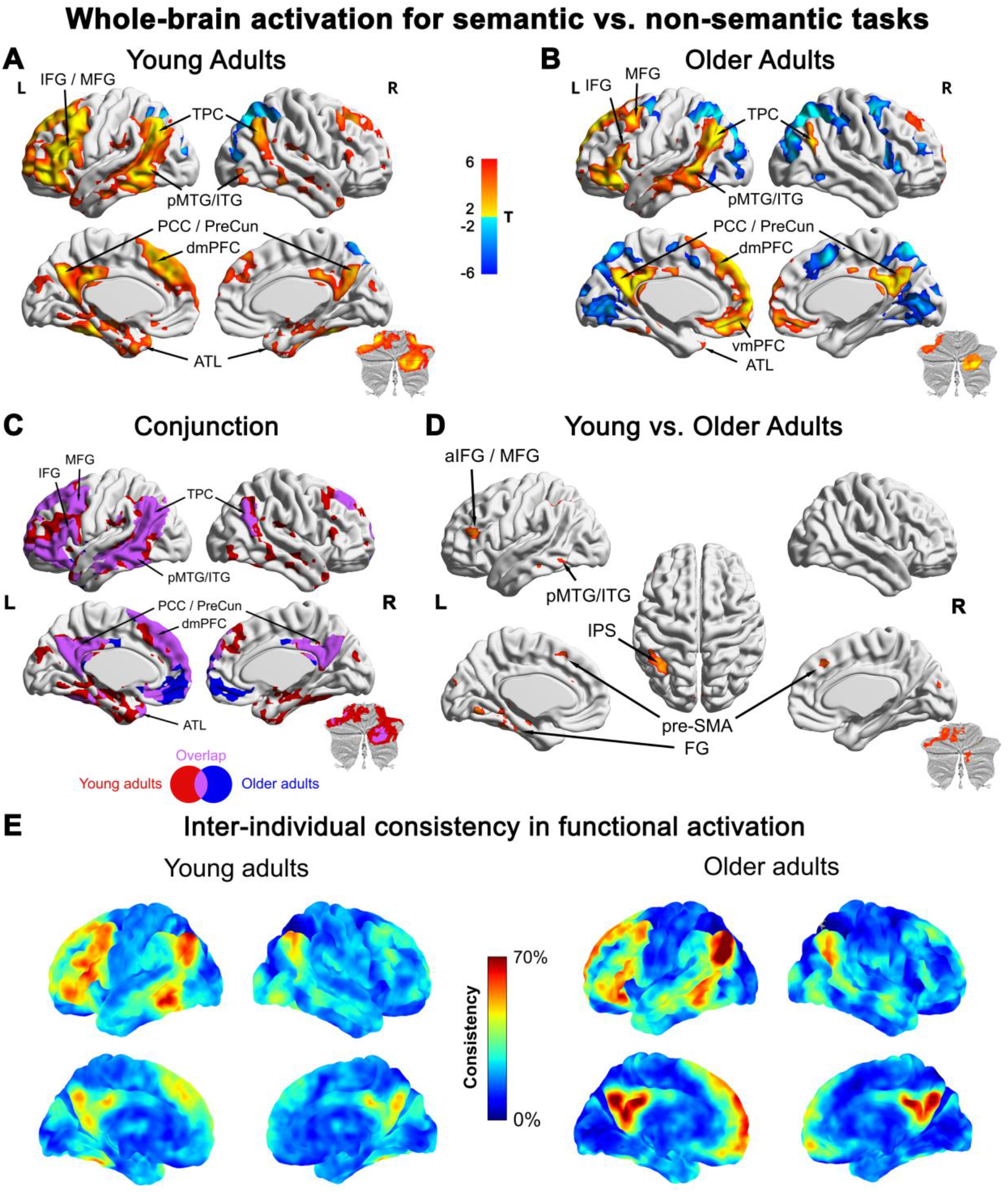
Whole-brain activation for general semantic processing in young (A) and older (B) adults, with their commonalities (C) and differences (D), as well as inter-individual consistency (E). All activation maps were thresholded at p < 0.05 FWE-corrected using threshold free cluster enhancement (TFCE). Inter-individual consistency in functional activation was assessed via threshold-weighted overlap maps.

Group comparison revealed that young adults showed higher semantic activity than older adults (Figure 3D; Table S5) in the left anterior IFG / MFG, pMTG/ITG, intraparietal sulcus (IPS), FG, as well as the bilateral pre-supplementary motor area (pre-SMA) and cerebellum (left lobules V/VI and right crus I/II). A supplementary analysis showed that left aIFG/MFG, pMTG/ITG and bilateral pre-SMA overlapped with the semantic control system (Figure S2), whereas left IPS exclusively overlapped with the domain-general multiple demand network (MDN) (Figure S3).

No brain region showed significantly higher semantic activity in older than young adults. Threshold-weighted overlap maps (Figure 3E) indicated that these group differences were not caused by larger inter-individual variability in older adults, suggesting that they indeed reflected higher activation magnitudes in young adults.

### Functional region-of-interest (fROI) analysis: TPC & ATL

To characterize task and age effects on the response profiles of semantic brain regions for taxonomic and thematic relationships, we performed a subject-specific functional region-of-interest (fROI) analysis. We identified fROIs using the general semantic contrast (semantic > non-semantic tasks) and then exclusively compared activity between different semantic conditions. As our primary analysis, we investigated fROIs in the left and right TPC and ATL where the highest proportion of participants showed significant activation. While our main ATL fROIs were located in the lateral ATL, we performed a supplementary analysis in the ventral ATL (Figure S7), as this area is widely considered the core location of the semantic “hub” (Binney et al., 2010; Lambon Ralph et al., 2016). The complete response profiles of all fROIs are displayed in Figure S5. For readability, we only report significant effects here (see the Supplementary Material for all ANOVAs and post-hoc comparisons).

According to the dual-hub theory, the TPC should be preferentially engaged for thematic relationships (as compared to taxonomic and unrelated pairs), whereas the ATL should prefer taxonomic relationships (versus thematic and unrelated pairs). We expected these activation patterns to be modulated by the task, with increased responses for task-relevant relationships. Finally, we expected an age-related shift from taxonomic towards thematic processing, in line with behavioral shifts towards thematic reasoning.

#### Left TPC

Left TPC (Figure 4A; Tables S6-S11) showed a GROUP x TASK interaction (F_1,73_ = 6.90, p = 0.01, η^2^_p_ = 0.086), driven by stronger activation for taxonomic judgments in young than older adults (t = 1.171, p = 0.044, Cohen’s d = 0.408). Young adults showed stronger engagement for taxonomic than thematic judgments (t = 1.774, p = 0.04, Cohen’s d = 0.244), whereas older adults showed stronger engagement for thematic than taxonomic judgments (t = 1.938, p = 0.029, Cohen’s d = 0.293).

**Figure 4.**
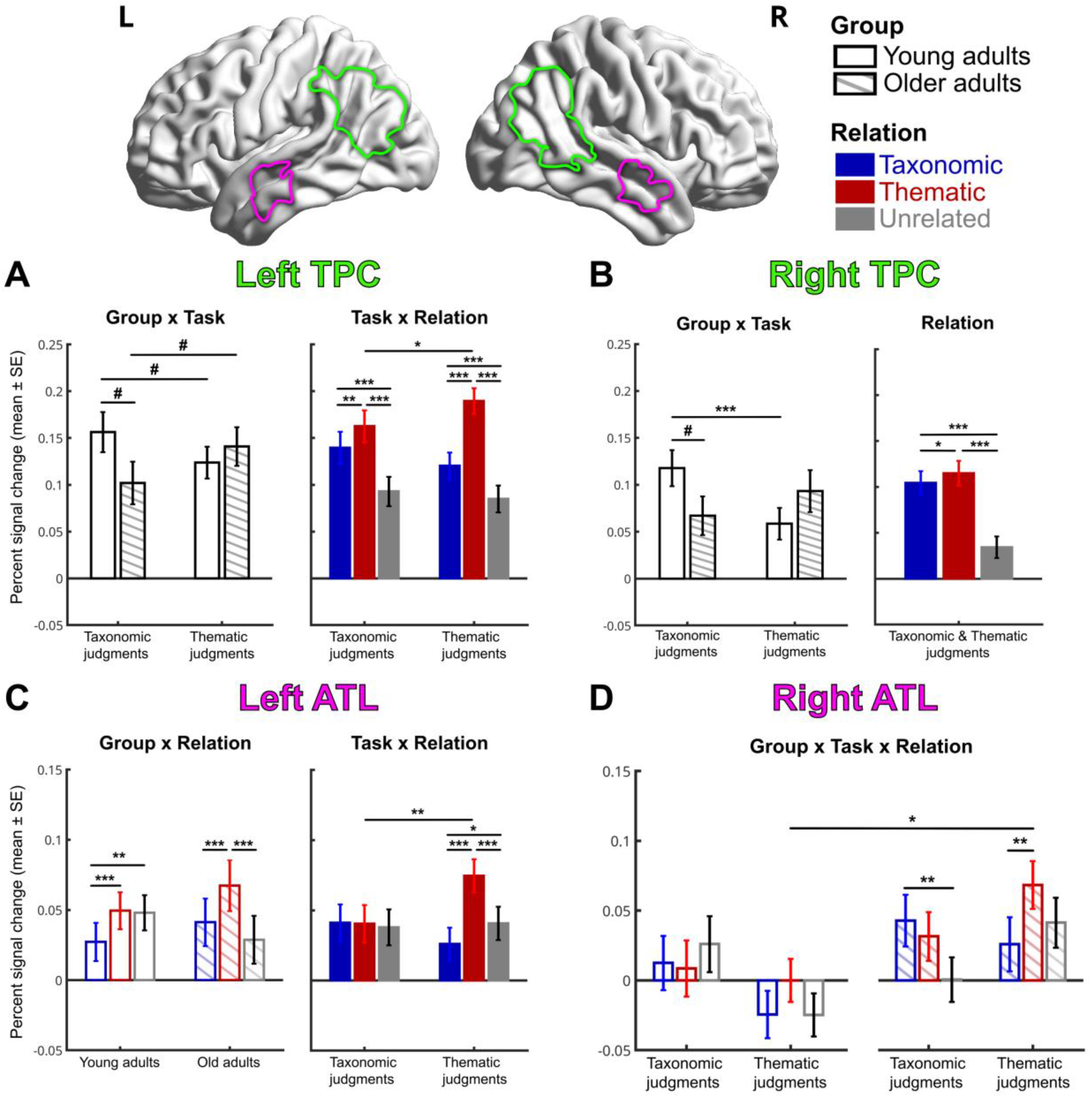
Response profiles for subject-specific fROIs in the left and right TPC and ATL. Mean percent signal change is shown for each experimental condition; error bars represent the standard error of the mean. ***p < 0.001 (corr.), **p < 0.01 (corr.), *p < 0.05 (corr.), ^#^p < 0.05 (uncorr.).

Moreover, a TASK x RELATION interaction (F_2,146_ = 7.35, p = 0.001, η^2^_p_ = 0.091) revealed stronger activity for thematic relationships during thematic than taxonomic judgments (t = 2.093, p = 0.013, Cohen’s d = 0.221) across groups. During both tasks, left TPC exhibited higher activity for thematic than taxonomic pairs (taxonomic task: t = 3.398, p = 0.001, Cohen’s d = 0.181; thematic task: t = 7.281, p < 0.001, Cohen’s d = 0.520), thematic than unrelated pairs (taxonomic task: t = 7.497, p < 0.001, Cohen’s d = 0.529; thematic task: t = 11.813, p < 0.001, Cohen’s d = 0.781), and taxonomic than unrelated pairs (taxonomic task: t = 4.449, p < 0.001, Cohen’s d = 0.349; thematic task: t = 4.214, p < 0.001, Cohen’s d = 0.262).

Overall, left TPC showed a preference for thematic (vs. taxonomic) relationships, in line with the dual-hub view. Notably, this thematic preference was pronounced when task-relevant. Moreover, left TPC exhibited an age-related decrease in taxonomic processing.

#### Right TPC

Right TPC (Figure 4B; Tables S12-S16) also showed a GROUP x TASK interaction (F_1,73_ = 10.943, p = 0.001, η^2^_p_ = 0.130), reflecting higher activity for taxonomic judgments in young than old adults (t = 1.792, p = 0.039, Cohen’s d = 0.395). Only in young adults, right TPC was more engaged during taxonomic than thematic judgments (t = 3.409, p < 0.001, Cohen’s d = 0.462).

In addition, a main effect of RELATION (F_2,146_ = 93.321, p < 0.001, η^2^_p_ = 0.561) showed that across age groups and tasks, right TPC was more engaged for thematic than taxonomic (t = -2.256, p = 0.027, Cohen’s d = 0.086), thematic than unrelated (t = 11.729, p < 0.001, Cohen’s d = 0.624), and taxonomic than unrelated (t = 9.701, p < 0.001, Cohen’s d = 0.538) pairs.

Overall, right TPC showed a similar response profile to its left-hemispheric homologue, with a preference for thematic (vs. taxonomic) relationships, in line with the dual-hub view. Right TPC also showed an age-related decrease in taxonomic processing. In contrast to left TPC, right TPC’s thematic response was not modulated by task relevance.

#### Left ATL

Left ATL (Figure 4C; Tables S17-S22) exhibited a GROUP x RELATION interaction (F_2,146_ = 9.687, p < 0.001, η^2^_p_ = 0.117), reflecting a clear preference in older adults for thematic vs. taxonomic (t = 4.096, p < 0.001, Cohen’s d = 0.206) and thematic vs. unrelated (t = 5.680, p < 0.001, Cohen’s d = 0.357) pairs, whereas young adults did not show a difference between thematic and unrelated pairs (t = 0.225, p = 0.823, Cohen’s d = 0.013).

Moreover, a TASK x RELATION interaction (F_2,146_ = 14.434, p < 0.001, η^2^_p_ = 0.165) revealed no differences between conditions during taxonomic judgments (all p > 0.9). However, during thematic judgments, left ATL showed stronger engagement for thematic than taxonomic (t = 6.787, p < 0.001, Cohen’s d = 0.456), thematic than unrelated (t = 5.592, p < 0.001, Cohen’s d = 0.325), and taxonomic than unrelated (t = 2.199, p = 0.031, Cohen’s d = 0.131) pairs. Activation for thematic relationships was significantly stronger during thematic than taxonomic judgments (t = 2.754, p = 0.002, Cohen’s d = 0.336).

Overall, left lateral ATL showed a selective and task-dependent response to thematic relationships, with no response to taxonomic relationships (even as compared to unrelated pairs). The selectivity for thematic relationships was particularly pronounced in older adults. These results contradict the dual-hub view, which predicts a preference for taxonomic relationships, as compared to thematic and unrelated pairs.

A supplementary analysis in left ventral ATL (Figure S7A; Tables S27-S31) yielded qualitatively similar results with a general preference for thematic over taxonomic relationships (main effect of RELATION: p < 0.001), and a trend towards an age-related increase in thematic processing (GROUP x TASK interaction: p = 0.07).

#### Right ATL

Right ATL (Figure 4D; Tables S23-S26) showed a GROUP x TASK x RELATION interaction (F_2,146_ = 4.978, p = 0.009, η^2^_p_ = 0.064). This occurred as young adults did not show any activation differences between conditions (all p > 0.3). In contrast, older adults showed higher activity for taxonomic than unrelated pairs during taxonomic judgments (t = 3.058, p = 0.003, Cohen’s d = 0.384), and for thematic than taxonomic relationships during thematic judgments (t = 4.284, p < 0.001, Cohen’s d = 0.385). Activity for thematic relationships during thematic judgments was significantly higher in older than young adults (t = 2.970, p = 0.004, Cohen’s d = 0.621).

Overall, right lateral ATL was only recruited in older adults, not in young adults. In older adults, right ATL showed a task-dependent response to both taxonomic and thematic relationships, with the strongest evidence for an age-related increase in thematic processing.

A supplementary analysis in right ventral ATL (Figure S7B; Tables S32-S35) showed qualitatively similar results with a trend towards stronger activation in older than young adults (main effect of GROUP: p = 0.07), and a preference for thematic relationships as compared to taxonomic and unrelated pairs (main effect of RELATION: p < 0.001).

### fROI Analysis: Other semantic regions

Beyond the bilateral TPC and ATL, we identified 7 fROIs where a high proportion of participants (>70%) showed significant activation for general semantic processing: the left precentral sulcus (PreCS), IFG/MFG, IPS, pMTG/ITG, pars orbitalis of the IFG (IFGorb), as well as the bilateral PreCun and mPFC. Their complete response profiles are displayed in Figure S6.

We expected that semantic control regions (especially left IFG/MFG and pMTG/ITG) should show a task-dependent response to taxonomic and thematic relationships: They should be engaged during the retrieval of task-relevant semantic information (taxonomic relations during taxonomic judgments, thematic relations during thematic judgments), as well as during the inhibition of task-irrelevant semantic information (thematic relations during taxonomic judgments, taxonomic relations during thematic judgments), as compared to unrelated pairs. Moreover, we hypothesized an age-related shift from taxonomic towards thematic processing throughout the semantic network.

#### Left PreCS

Left PreCS (Figure 5A; Tables S36-S40) showed a GROUP x TASK interaction (F_2,146_ = 5.228, p = 0.025, η^2^_p_ = 0.067), revealing that only older adults exhibited higher activity for thematic than taxonomic judgments (t = 1.916, p = 0.029, Cohen’s d = 0.280).

**Figure 5.**
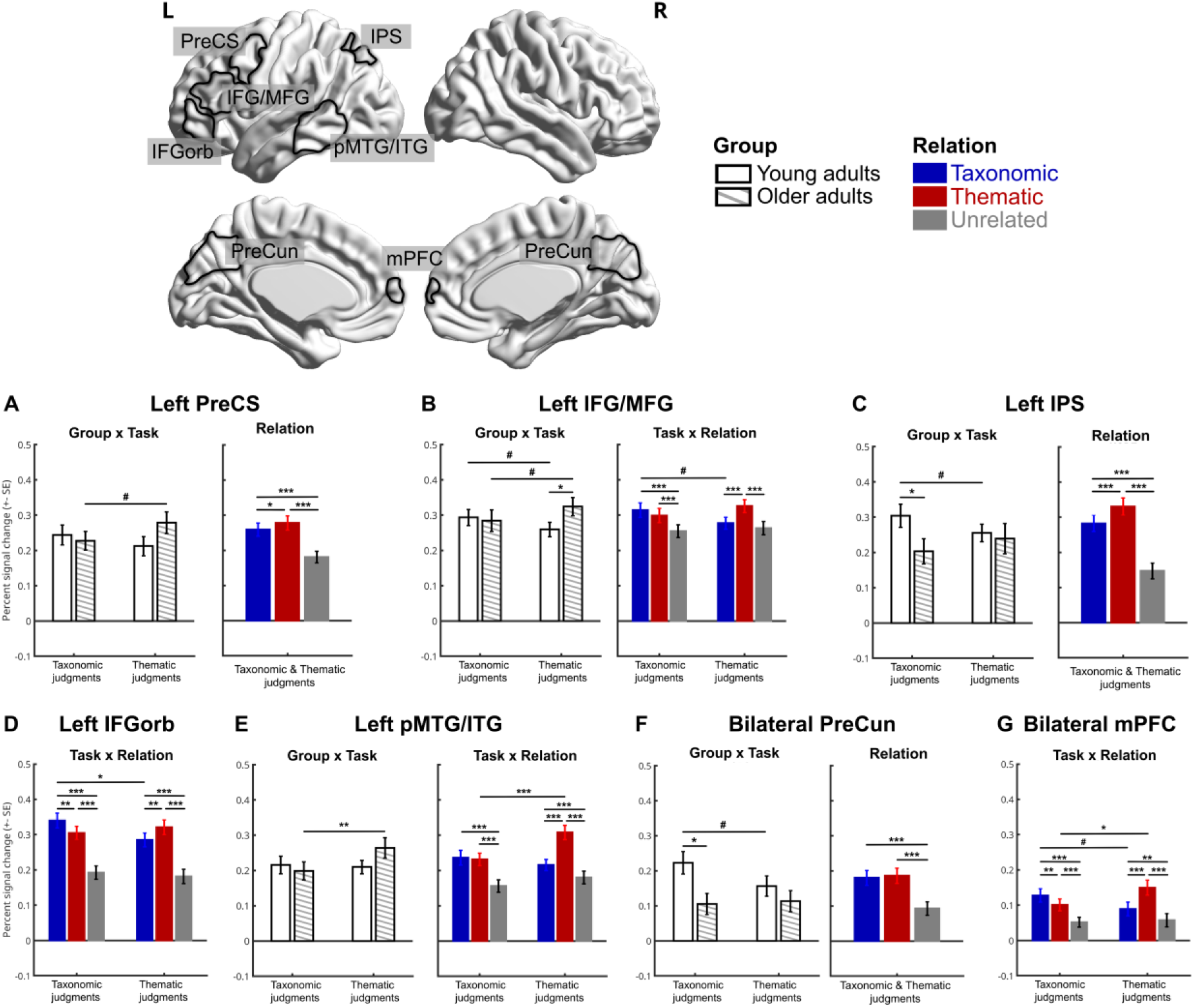
Response profiles for subject-specific fROIs in semantic regions beyond TPC and ATL. Mean percent signal change is shown for each experimental condition; error bars represent the standard error of the mean. ***p < 0.001 (corr.), **p < 0.01 (corr.), *p < 0.05 (corr.), ^#^p < 0.05 (uncorr.).

Moreover, a RELATION main effect (F_2,146_ = 60.551, p < 0.001, η^2^_p_ = 0.002) revealed higher activity across groups and tasks for thematic than taxonomic (t = 2.641, p = 0.01, Cohen’s d = 0.103), thematic than unrelated (t = 8.814, p < 0.001, Cohen’s d = 0.526), and taxonomic than unrelated (t = 8.278, p < 0.001, Cohen’s d = 0.423) pairs.

Overall, left PreCS showed a general, task-independent preference for thematic (vs. taxonomic) relationships, but was also sensitive to taxonomic (vs. unrelated) relations. Only in older adults, left PreCS showed an increased response to thematic (vs. taxonomic) judgments.

#### Left IFG/MFG

Left IFG/MFG (Figure 5B; Tables S41-S46) showed a GROUP x TASK interaction (F_1,73_ = 6.002, p = 0.017, η^2^_p_ = 0.076). Young adults exhibited higher activity for taxonomic than thematic judgments (t = 1.675, p = 0.049, Cohen’s d = 0.209), whereas old adults showed higher activity for thematic than taxonomic judgments (t = 1.789, p = 0.039, Cohen’s d = 0.245). Activity for thematic judgments was higher in young than old adults (t = 1.989, p = 0.025, Cohen’s d = 0.4).

In addition, left IFG/MFG exhibited a TASK x RELATION interaction (F_2,146_ = 7.921, p < 0.001, η^2^_p_ = 0.098). In the taxonomic task, activity was stronger for taxonomic than unrelated (t = 4.421, p < 0.001, Cohen’s d = 0.359) and thematic than unrelated (t = 3.735, p = 0.001, Cohen’s d = 0.275) pairs. In the thematic task, activity was higher for thematic than taxonomic (t = 3.585, p < 0.001, Cohen’s d = 0.292) and thematic than unrelated (t = 4.549, p < 0.001, Cohen’s d = 0.373) pairs.

Overall, left IFG/MFG exhibited a task-dependent response for thematic relationships: Thematic relations were preferred when task-relevant. At the same time, left IFG/MFG showed a strong response for the inhibition of task-irrelevant relations, especially during taxonomic judgments. This response profile is consistent with semantic control. Moreover, left IFG/MFG showed an age-related increase in thematic processing.

#### Left IPS

Left IPS (Figure 5C; Tables S47-S51) exhibited a GROUP x TASK interaction (F_1,73_ = 4.908, p = 0.030, η^2^_p_ = 0.063), driven by higher activity for taxonomic judgments in young than old adults (t = 2.078, p = 0.021, Cohen’s d = 0.454). Only young adults showed increased activity for taxonomic vs. thematic judgments (t = 1.893, p = 0.031, Cohen’s d = 0.220).

Left IPS also showed a RELATION main effect (F_1,73_ = 109.076, p < 0.001, η^2^_p_ = 0.599), reflecting higher activity for thematic than taxonomic (t = 4.835, p < 0.001, Cohen’s d = 0.219), thematic than unrelated (t = 12.082, p < 0.001, Cohen’s d = 0.826), and taxonomic than unrelated (t = 10.501, p < 0.001, Cohen’s d = 0.607) pairs.

Overall, left IPS showed a general, task-independent preference for thematic (vs. taxonomic) relationships, but was also sensitive to taxonomic (vs. unrelated) relations. Moreover, left IPS exhibited an age-related decrease in taxonomic processing.

#### Left IFGorb

Left IFGorb (Figure 5D; Tables S52-S55) showed no group differences, but a TASK x RELATION interaction (F_2,146_ = 8.364, p < 0.001, η^2^_p_ = 0.103). This occurred as activity was higher for taxonomic than thematic relationships during taxonomic judgments (t = 3.136, p = 0.002, Cohen’s d = 0.200), whereas activity was higher for thematic than taxonomic relationships during thematic judgments (t = 2.548, p = 0.010, Cohen’s d = 0.202). During both tasks, activity was increased for taxonomic vs. unrelated pairs (taxonomic task: t = 9.516, p < 0.001, Cohen’s d = 0.854; thematic task: t = 9.068, p < 0.001, Cohen’s d = 0.598) and for thematic vs. unrelated pairs (taxonomic task: t = 9.683, p < 0.001, Cohen’s d = 0.654; thematic task: t = 9.751, p < 0.001, Cohen’s d = 0.800).

Overall, left IFGorb showed a flexible, task-dependent response to task-relevant semantic relationships, responding maximally to taxonomic relations during taxonomic judgments and to thematic relations during thematic judgments. IFGorb activity was also elevated for task-irrelevant relationships, as compared to unrelated pairs. This response profile is in line with semantic control.

#### Left pMTG/ITG

Left pMTG/ITG (Figure 5E; Tables S56-S61) exhibited a GROUP x TASK interaction (F_1,73_ = 4.920, p = 0.030, η^2^_p_ = 0.063). Only in older adults, activity was higher during thematic than taxonomic judgments (t = 2.761, p = 0.004, Cohen’s d = 0.410).

Moreover, a TASK x RELATION interaction (F_2,146_ = 24.324, < 0.001, η^2^_p_ = 0.250) revealed that only during thematic judgments, activity was higher for thematic than taxonomic relationships (t = 8.483, p < 0.001, Cohen’s d = 0.579). During both tasks, activity was increased for taxonomic vs. unrelated pairs (taxonomic task: t = 6.570, p < 0.001, Cohen’s d = 0.500; thematic task: t = 3.711, p < 0.001, Cohen’s d = 0.219) and for thematic vs. unrelated pairs (taxonomic task: t = 7.273, p < 0.001, Cohen’s d = 0.472; thematic task: t = 10.690, p < 0.001, Cohen’s d = 0.798).

Overall, left pMTG/ITG showed a task-dependent preference for thematic (vs. taxonomic) relationships, but was also sensitive to taxonomic (vs. unrelated) relations. Only in older adults, left pMTG/ITG showed a preference for thematic vs. taxonomic tasks.

#### Bilateral PreCun

Bilateral PreCun (Figure 5F; Tables S62-S66) exhibited a GROUP x TASK interaction (F_1,73_ = 4.393, p = 0.040, η^2^_p_ = 0.057) as activity was higher for taxonomic judgments in young than older adults (t = 2.624, p = 0.006, Cohen’s d = 0.601). Only in young adults, activity was higher for taxonomic than thematic judgments (t = 2.784, p = 0.004, Cohen’s d = 0.341).

A RELATION main effect (F_2,146_ = 72.933, < 0.001, η^2^_p_ = 0.500) across groups and tasks showed that activity was increased for taxonomic vs. unrelated (t = 9.382, p < 0.001, Cohen’s d = 0.442) and thematic vs. unrelated (t = 9.720, p < 0.001, Cohen’s d = 0.476) pairs, but comparable for taxonomic and thematic pairs (t = 0.988, p = 0.326, Cohen’s d = 0.034).

Overall, bilateral PreCun showed a general, task-independent sensitivity to semantic relationships (vs. unrelated pairs), without a preference for taxonomic vs. thematic relations. Moreover, bilateral PreCun exhibited an age-related decrease in taxonomic processing.

#### Bilateral mPFC

Bilateral mPFC (Figure 5G; Tables S67-S70) showed no group differences, but a TASK x RELATION interaction (F_2,146_ = 16.755, < 0.001, η^2^_p_ = 0.187). Activity was higher for taxonomic than thematic relationships during taxonomic judgments (t = 2.786, p = 0.002, Cohen’s d = 0.170), whereas activity was higher for thematic than taxonomic relationships during thematic judgments (t = 5.833, p < 0.001, Cohen’s d = 0.385). During both tasks, activity was increased for taxonomic vs. unrelated pairs (taxonomic task: t = 5.684, p < 0.001, Cohen’s d = 0.487; thematic task: t = 2.868, p = 0.002, Cohen’s d = 0.191) and for thematic vs. unrelated pairs (taxonomic task: t = 4.170, p < 0.001, Cohen’s d = 0.317; thematic task: t = 8.086, p < 0.001, Cohen’s d = 0.576).

Overall, similar to left IFGorb, bilateral mPFC showed a flexible, task-dependent response to task-relevant semantic relationships (taxonomic during taxonomic judgments, thematic during thematic judgments). Bilateral mPFC also showed an increased response for task-irrelevant relationships, as compared to unrelated pairs. This response profile is clearly in line with semantic control.

### fROI Analysis: Relationship with behavioral performance

We correlated the difference in percent signal change between thematic and taxonomic tasks with the respective difference in response times and accuracy within each age group. We exclusively found brain–behavior relationships in older, not young, adults (Figure 6A). Specifically, relatively stronger activity in the bilateral PreCun and left IPS for the thematic vs. taxonomic task was associated with higher accuracy. In contrast, stronger activity for thematic than taxonomic judgments in the left ATL, IFG/MFG, IFGorb, pMTG/ITG, and PreCS correlated with slower response times in older adults. The strongest evidence for brain– behavior relationships was found in left IFG/MFG and PreCS, which were significant after FDR correction. Both showed increasing reaction times with increasing activity selectively in older adults (Figure 6B). We repeated the analysis considering only task-relevant items. Results were largely similar and showed the same significant correlation of increasing reaction times with increasing activity in the left IFG/MFG selectively in older adults (Figure S9).

**Figure 6.**
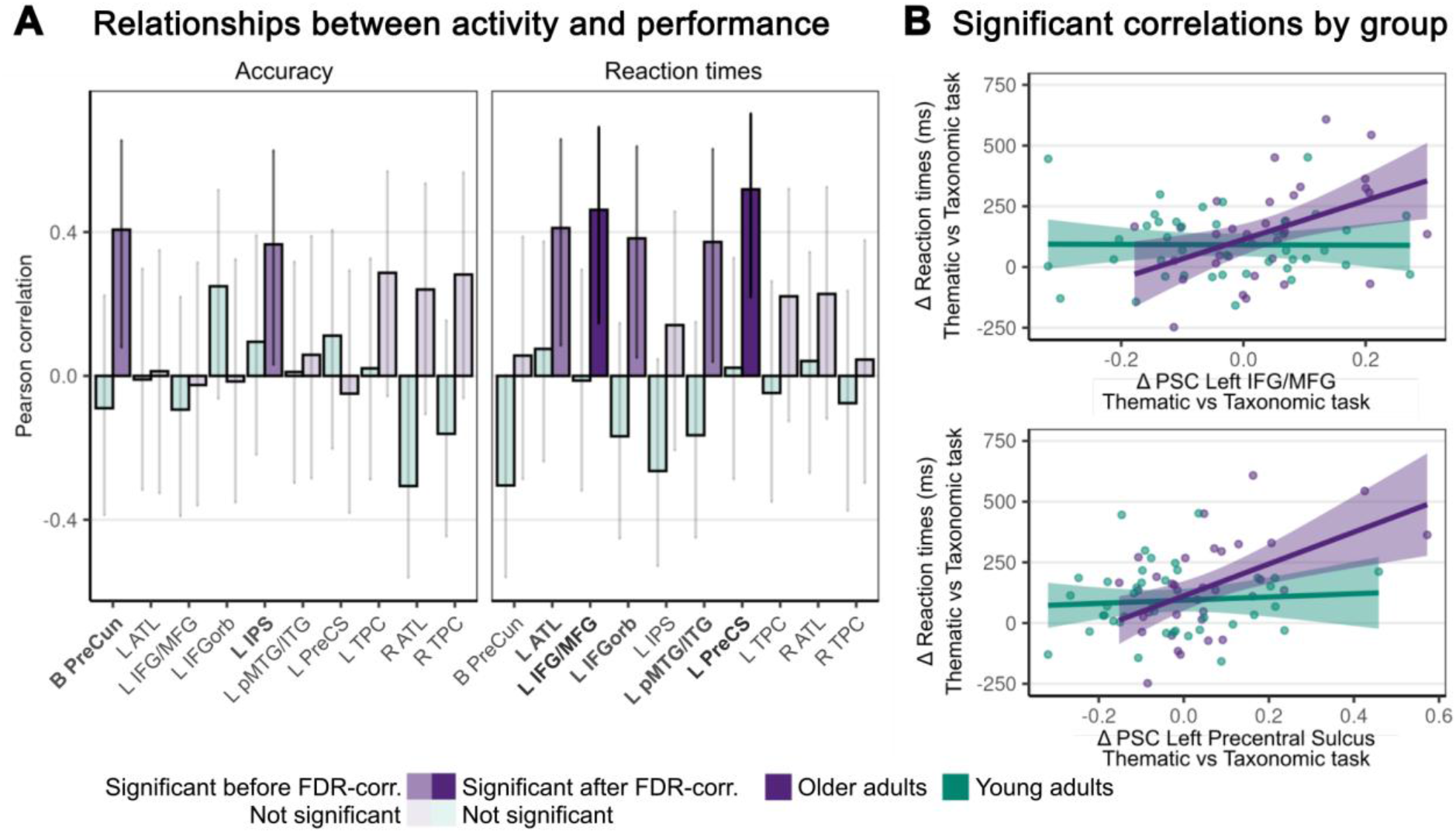
Brain–behavior relationships between task-related activity and behavioral performance. (A) Correlation strength of percent signal change in fROIs and behavioral performance by age group. (B) Significant correlations after FDR correction.

## Discussion

This fMRI study investigated how taxonomic and thematic relationships are represented in the human brain as a function of task and aging. Young and older adults performed taxonomic and thematic judgment tasks on picture pairs that were taxonomically related, thematically related, or unrelated.

Three key findings stand out. First, our results do not support the dual-hub theory, which proposes a preference for taxonomic relationships in the ATL and for thematic relationships in the TPC. Instead, both regions consistently preferred thematic relations.

Second, the response of several semantic areas for taxonomic and thematic relationships was task-dependent. Left TPC and ATL showed an enhanced response to task-relevant *thematic* relationships. Semantic control regions (particularly left IFG and mPFC) were flexibly engaged for task-relevant semantic relations.

Third, aging was associated with reduced activity for taxonomic processing and increased activity for thematic processing in different semantic regions. Selectively in older adults, relatively stronger activity for thematic than taxonomic judgments was associated with higher accuracy but slower responses. This suggests that older adults require increased cognitive resources to maintain accuracy at a high level, but this comes at the cost of processing efficiency.

### Challenging the dual-hub account

The dual-hub theory proposes that taxonomic relations are represented in the ATL, whereas thematic relations are represented in the TPC (including angular gyrus and posterior temporal cortex) (Davis and Yee, 2019; Mirman et al., 2017). Our data only partially align with this view.

Consistent with the dual-hub view, both left and right TPC indeed preferred thematic over taxonomic relationships. This finding converges with previous neuroimaging (de Zubicaray et al., 2013; Zhang et al., 2023) and neuropsychological lesion (Schwartz et al., 2011; Tsagkaridis et al., 2014) evidence for a thematic bias in the TPC. However, both left and right TPC were also sensitive to taxonomic relationships, as compared to unrelated images. This indicates that the TPC does not selectively respond to thematic relationships, but is sensitive to semantic relatedness more generally (Kuhnke et al., 2023b; Tong et al., 2022).

One mechanistic explanation for these results could be that the TPC functions as a multimodal convergence zone or “hub” that binds semantic features across perceptual-motor modalities (Fernandino et al., 2016; Kiefer et al., 2024; Kuhnke et al., 2025, 2022). Multimodal perceptual-motor information is particularly important for forming event-based thematic relations: The representation of events – also known as “situation models” – requires the binding of multimodal semantic content related to the participating entities, their action-based interactions, and the spatio-temporal context (Casto et al., 2025; Fernandino and Binder, 2024; Kemmerer, 2025; Kemmerer and Dove, 2026). However, the integration of multiple features of concepts is also relevant for forming taxonomic relationships, which rely on shared feature (Davis and Yee, 2019; Lambon Ralph et al., 2010b; Mirman et al., 2017). This may explain the contribution of the TPC to both taxonomic and thematic processing.

The ATL results pose a more direct challenge to the dual-hub view. Instead of a taxonomic preference, left ATL showed a robust task-dependent preference for *thematic* relations. Notably, we used dual-echo fMRI, slice tilting and susceptibility distortion correction to maximize BOLD sensitivity in the ATL (Halai et al., 2014; Poser et al., 2006; Weiskopf et al., 2006), rendering it highly unlikely that the absence of taxonomic effects was caused by measurement insensitivity. Moreover, our results are in line with a neuroimaging meta-analysis which found no taxonomic bias but a consistent preference for thematic relations in the lateral ATL (Zhang et al., 2023), as well as a large-scale neuropsychological lesion study which linked ATL damage to thematic naming deficits (Riccardi et al., 2024).

The right ATL was engaged for semantic vs. non-semantic tasks, but did not differentiate between semantic relation types in young adults. In older adults, however, right ATL showed a task-dependent response to both taxonomic and thematic relations. These findings support the view that ATL involvement in semantic cognition is bilateral (Jefferies, 2013; Lambon Ralph et al., 2016). In healthy young adults, left ATL seems to be dominant, with right ATL playing a subordinate role (Binney and Lambon Ralph, 2015; Jung and Lambon Ralph, 2016). However, the right ATL can compensate (to some extent) when the left ATL is perturbed or even damaged, as demonstrated via plasticity-inducing neurostimulation (Binney and Lambon Ralph, 2015; Jung and Lambon Ralph, 2016) and patients with unilateral ATL lesions (Lambon Ralph et al., 2012, 2010a). Our findings suggest a similar role during aging: The right ATL becomes more engaged in the processing of semantic relationships in older adults, potentially to compensate for age-related cognitive decline and to maintain a high level of semantic performance (Martin et al., 2023b, 2022). The finding is also in line with a recent meta-analysis on age-related changes to semantic processing in the brain, which found increased task-specific activity in the right hemisphere, including right ATL, especially when performance decreased (Hoffman and Morcom, 2018). Similarly, the “Hemispheric Asymmetry Reduction in Older Adults” (HAROLD) model assumes that the increased bilateral activity of the prefrontal cortex for cognitive tasks in older adults may be compensatory (Cabeza, 2002). Alternatively, more bilateral activity could reflect a loss of specialized, efficient neural organization rather than beneficial compensation.

Overall, in contrast to the dual-hub view that conceptualizes the ATL as a taxonomic node, our data fit better with models of the ATL as a semantic “hub” that integrates distributes semantic features associated with the same concept (Kuhnke et al., 2023a; Lambon Ralph et al., 2016; Patterson et al., 2007). According to these theories, the ATL represents a general abstract semantic similarity structure, regardless of whether that similarity is based on taxonomic or thematic relations (Jackson et al., 2015; Lambon Ralph et al., 2010b).

Notably, our ATL fROIs were located in the lateral ATL, as this was the anterior temporal region where the largest proportion of participants showed significant activation for the semantic > non-semantic contrast. However, the semantic “hub” is often localized to the ventral ATL, that is, anterior portions of the fusiform and/or inferior temporal gyri (Binney et al., 2010; Lambon Ralph et al., 2016). Therefore, we performed a supplementary analysis in the ventral ATL. This analysis yielded qualitatively converging results (Figure S7): Also the bilateral ventral ATL prefers thematic over taxonomic relations. Left ventral ATL shows an age-related increase in thematic processing, and right ventral ATL is generally more engaged in semantic processing in older than young adults.

One crucial open question is whether TPC and ATL play similar or distinct roles as semantic hubs. The present results show that both TPC and ATL respond to thematic and taxonomic relationships (as compared to unrelated pairs), with a preference for thematic relations. We propose that both regions integrate distributed semantic features from various perceptual-motor modalities. We can only speculate on functional differences between TPC and ATL based on other studies: Previous work suggests that the TPC acts as a “multimodal” hub that retains modality-specific information, whereas the ATL is an “amodal” hub that completely abstracts away from modality-specific content (Kuhnke et al., 2023a, 2022; Reilly et al., 2016). This is supported by previous evidence that the TPC is sensitive to semantic information related to several individual perceptual-motor modalities (Bonner et al., 2013; Fernandino et al., 2016; Kuhnke et al., 2020b; Tong et al., 2022), whereas the ATL is engaged in semantic processing in a modality-invariant fashion (Lambon Ralph et al., 2016; Patterson et al., 2007). Future studies should directly test the proposed multimodal vs. amodal distinction in TPC vs. ATL.

### Task-dependent activity for taxonomic and thematic relationships

Several semantic brain regions showed a task-dependent response to taxonomic and thematic relationships: Activity was enhanced when a given relation type was task-relevant.

Left TPC and ATL activity was increased for thematic relations during thematic as compared to taxonomic judgments, further supporting the view that the thematic bias in these “hubs” is not fixed but amplified when task-relevant (Binder and Desai, 2011; Kuhnke et al., 2020a).

Semantic control regions showed a flexible response to task-relevant semantic relations. The strongest evidence for this response profile was found in left IFGorb and mPFC, with maximal activity in congruent trials, that is, for taxonomic relations during taxonomic judgments, and for thematic relations during thematic judgments. At first glance, these results may be surprising. In classical paradigms, semantic control regions are engaged when a context-appropriate semantic relationship has to be retrieved in the presence of context-inappropriate competitors (Davey et al., 2015; Whitney et al., 2012). Thus, one may expect semantic control regions to be most strongly engaged in incongruent trials (i.e. taxonomic relations during thematic judgments, thematic relations during taxonomic judgments). However, our experimental design differs from classical semantic control paradigms. In our design, only a single relationship had to be evaluated on each trial, and congruent and incongruent trials were qualitatively different in their semantic control demands and response requirements: In congruent trials, a context-appropriate relationship had to be retrieved and approved (“yes” response). In incongruent trials, a context-inappropriate relationship had to be inhibited and rejected (“no” response). Hence, the asymmetry in our experimental design created a separation of different semantic control processes: Controlled retrieval (and approval) of task-relevant information vs. inhibition (and rejection) of task-irrelevant information.

Importantly, our results indicate that semantic control regions are involved in both processes. Behaviorally, congruent relations were associated with *worse* accuracy than incongruent relations in young adults, indicating that retrieval of task-relevant relations for “yes” responses was cognitively demanding and required control, even more than inhibition of task-irrelevant relations for “no” responses. These results are in line with previous evidence that semantic control regions support the retrieval of task-relevant information, even in the absence of task-irrelevant competitors (Shao et al., 2026; Zhang et al., 2021).

Crucially, however, semantic control regions also showed increased activity for the inhibition of task-irrelevant semantic relationships in incongruent trials, as compared to unrelated pairs. Left IFG/MFG and pMTG/ITG even showed an equal response to congruent and incongruent relationships during taxonomic judgments. This pattern was confirmed in a supplementary analysis that controlled for a potential bias in fROI definition towards congruent relations (Figure S8).

Overall, our results highlight that semantic control regions are engaged both in the retrieval of task-relevant semantic information and the inhibition of task-irrelevant information. Our findings align with previous neuroimaging and brain stimulation work that implicates left anterior IFG (Badre et al., 2005; Hodgson et al., 2021; Jackson, 2021; Martin et al., 2025; Wagner et al., 2001), pMTG/ITG (Hallam et al., 2016; Noonan et al., 2013; Whitney et al., 2011) and mPFC (Jackson, 2021; Liu et al., 2025) in the controlled retrieval of context-appropriate semantic information and the suppression of context-inappropriate information.

While the taxonomic–thematic distinction has been highly fruitful to study the representation of semantic relationships in the human brain, our results suggest that this dichotomy may be too coarse to fully capture the organization of semantic knowledge. The observed graded and partially overlapping neural responses suggest that both relation types comprise heterogeneous subcomponents that are flexibly recruited depending on task demands. Taxonomic relations can rely on perceptual feature overlap or abstract category structure, whereas thematic relations can draw on action, event, or contextual associations (Davis and Yee, 2019; Mirman et al., 2017). Consequently, broad taxonomic–thematic contrasts are unlikely to map cleanly onto distinct neural substrates. Our findings support a shift toward more fine-grained accounts that decompose semantic relations into their underlying feature components, an approach increasingly advocated in recent work (Zhang et al., 2023).

More generally, our results strongly support the view that semantic cognition relies on a dynamic, context-sensitive neural architecture (Hoenig et al., 2008; Kuhnke et al., 2021, 2020b; Popp et al., 2019). The neural representation of a semantic relationship is not a fixed, task-independent entity, but it is flexibly shaped to the requirements of the current context (Kemmerer, 2015; Lebois et al., 2015; Yee and Thompson-Schill, 2016).

### Aging and the shift from taxonomic to thematic processing

A major goal of this study was to examine how taxonomic and thematic processing change with healthy aging. Behaviorally, both groups performed well, but young adults were generally faster than older adults. Older adults showed worse accuracy on taxonomic than thematic relationships, especially during thematic judgments. These findings are broadly consistent with the previously reported U-shaped trajectory across the life span: from a thematic bias in childhood to taxonomic dominance in young adulthood and back to thematic preference in older age (Cicirelli, 1976; Kogan, 1974; Smiley and Brown, 1979).

At the neural level, young and older adults showed a similar general semantic network (for semantic > non-semantic tasks) comprising bilateral TPC and ATL, PCC/PreCun, left IFG/MFG, pMTG/ITG, and bilateral cerebellum (Binder et al., 2009; Kuhnke et al., 2023a; Turker et al., 2023). However, older adults exhibited reduced semantic activity in domain-general executive control or multiple demand network (MDN) nodes (left IPS) and domain-specific semantic control regions (left anterior IFG/MFG, pMTG/ITG and pre-SMA). The reduced activity in semantic control regions aligns with prior evidence for age-related decline in controlled semantic retrieval (Hoffman and Morcom, 2018). In contrast, numerous studies have reported increased activity in the domain-general MDN with aging (Hoffman and Morcom, 2018; Kang et al., 2022). This apparent discrepancy could reflect hemispheric and/or regional differences within the networks of semantic and domain-general cognitive control: Age-related activity increases are most consistently observed in right-hemispheric and prefrontal MDN areas (Hoffman and Morcom, 2018; Martin et al., 2022). These findings further support an increased recruitment of the right hemisphere in older age, in line with the HAROLD model and our result of increased right ATL involvement in the processing of semantic relationships. In contrast, left-hemispheric and parietal nodes like IPS may exhibit more heterogeneous responses, ranging from reduced to preserved recruitment depending on task-specific demands and individual performance differences (Hoffman and Morcom, 2018; Li et al., 2015).

Within the semantic tasks, older adults showed reduced activity for taxonomic judgments and/or increased activity for thematic judgments in several semantic areas, further supporting an age-related shift of the semantic network away from taxonomic processing towards thematic processing. This finding corroborates the notion of age-related “neural dedifferentiation”, conceptualized as a reduced specificity of neural representations (Koen et al., 2020; Koen and Rugg, 2019) and broader, less precise task-related functional activation (Chan et al., 2014; Martin et al., 2023b, 2022). For example, older adults show less distinct activation patterns for visual categories in occipito-temporal cortex (Koen et al., 2020; Park et al., 2004). Our results extend the previous literature by showing that the concept of neural dedifferentiation also applies to semantic representations, where thematic relationships can be seen as less specialized and broader than taxonomic relationships.

One may wonder whether our age effects could be explained by cohort differences. For instance, young adults may no longer be familiar with concepts like *payphone* or *cassette tape*. For these reasons, our stimulus set selectively included common objects and relationships that are well-known across generations (e.g. *cherry* - *pear*; *cow* - *grass*) (Table S71). Moreover, semantic relatedness ratings of our fMRI participants showed no effect of age (Table S73). Therefore, we are confident that our results cannot be explained by cohort effects and indeed reflect aging.

Brain–behavior correlations revealed that only in older adults, increased task-related activity for thematic vs. taxonomic judgments was associated with higher accuracy but slower response in different semantic regions. Activity was positively correlated with better accuracy in bilateral PreCun and left IPS. This is an intriguing result as the PreCun is associated with the default mode network (DMN) (Buckner, 2013; Raichle, 2015), while the IPS is part of the MDN (Assem et al., 2020; Duncan, 2010), which are usually anti-correlated. In line with the “default-executive coupling hypothesis of aging” framework (DECHA; Spreng and Turner, 2019; Turner and Spreng, 2015) and our previous results (Martin et al., 2023b, 2022), this finding corroborates the notion that increased functional coupling of DMN and MDN underlies age-related changes in cognition. Indeed, in semantic cognition, the joint contribution of DMN and MDN seems to be crucial for successful task performance, especially when controlled access to semantic memory is required (Krieger-Redwood et al., 2016; Kuhnke et al., 2023b; Martin et al., 2022). It is now widely recognized that the DMN plays a crucial role in semantic cognition, potentially by constructing multimodal situation models (Fernandino and Binder, 2024; Kemmerer and Dove, 2026).

In contrast, higher activity in semantic control regions (left IFG/MFG, IFGorb, PreCS, and pMTG/ITG) was associated with slower responses. As our data are correlational, it cannot be inferred that increased semantic control recruitment is maladaptive in older adults. Rather, these results suggest that stronger recruitment of semantic control areas is required under heightened processing demands (Jackson, 2021; Noonan et al., 2013). This is in line with the view that additional recruitment of control regions in older adulthood might not reflect compensation but rather reduced processing efficiency or specificity (Morcom and Henson, 2018).

Overall, our results converge with a wealth of evidence supporting a “semanticization of cognition” over the course of aging (Spreng and Turner, 2019): Semantic knowledge (or “crystallized intelligence”) remains stable or even increases until late in life (Baltes et al., 1995; Hedden and Gabrieli, 2004), whereas executive functions and processing speed (“fluid intelligence”) decline (Salthouse, 2019). Our results add to this previous evidence by showing that not only domain-general control, but also domain-specific semantic control functions decline with age. This is supported by worse performance on neurocognitive tests requiring executive control and slower performance on all semantic tasks requiring controlled access to semantic memory.

## Conclusions

In conclusion, taxonomic and thematic representations in the human brain are strongly modulated by task and age. Our results argue against a dual-hub organization of taxonomic and thematic knowledge. Instead, semantic representation hubs in TPC and ATL process both taxonomic and thematic relationships, with a graded thematic bias. Semantic control regions flexibly guide the retrieval of task-relevant taxonomic or thematic representations, while inhibiting task-irrelevant information. Finally, aging is associated with a shift of the semantic network from taxonomic towards thematic processing. Higher activity for thematic than taxonomic processing is linked to better accuracy but slower responses, which mirrors the “semanticization of cognition”. Older adults require increased cognitive resources to maintain semantic accuracy at a high level, but this comes at the cost of processing efficiency.

## Statements & Declarations

## Acknowledgements

We thank the medical technical assistants of MPI CBS for their help with data acquisition and Toralf Mildner for setting up the dual-echo fMRI sequence. We are also grateful to two anonymous reviewers, as well as the editors of the special issue Emiko Muraki and Richard Binney for their thoughtful comments which significantly improved the manuscript.

## Author Contributions (CRediT)

Philipp Kuhnke: Conceptualization, Data curation, Formal analysis, Visualization, Writing— original draft, Writing—review and editing;

Sandra Martin: Formal analysis, Visualization, Writing—original draft, Writing—review and editing;

Curtiss A. Chapman: Conceptualization, Investigation, Data curation, Formal analysis, Writing—review and editing;

Gesa Hartwigsen: Conceptualization, Funding acquisition, Supervision, Project administration, Writing—review and editing.

## Data Availability Statement

Behavioral and fMRI data, derivatives and analysis code are openly available via the Open Science Framework (OSF): https://osf.io/h985f

