## Supplementary Material for "Beyond dual hubs: Task and aging shape taxonomic and thematic semantic relationships in the human brain"

Kuhnke, Martin et al. (2026) *Cortex*

### Supplementary Results

#### Behavioral Analyses

**Model formulas:**

| **Accuracy** | **Reaction times** |
|---|---|
| m_Acc <- glmmTMB(Accuracy ~<br>Task +<br>Condition +<br>Group +<br>Task_order +<br>Gender +<br>Task : Condition +<br>Task : Group +<br>Condition : Group +<br>Task : Condition : Group +<br>(1 + Task + Condition \| Participant) +<br>(1 \| Trial),<br>data = df_Acc,<br>family = binomial(link = "logit")) | m_RT <- lmerTest::lmer(log(RT) ~<br>Task +<br>Condition +<br>Group +<br>Task_order +<br>Gender +<br>Task : Condition +<br>Task : Group +<br>Task : Condition : Group +<br>(1 + Task \| Participant) +<br>(1 \| Trial),<br>data = df_RT,<br>REML = TRUE) |

**Table S1. Results from mixed-effects models for accuracy and reaction times in the semantic tasks.**

| | **Accuracy** | | | | **Reaction times** | | | |
|---|---|---|---|---|---|---|---|---|
| *Predictors* | *Odds Ratios* | *std. Error* | *Statistic* | *p* | *Estimates* | *std. Error* | *Statistic* | *p* |
| Task [Thematic vs Taxonomic] | 0.40 | 0.04 | -8.56 | **5.18e-17** | 0.07 | 0.01 | 6.26 | **1.37e-09** |
| Relation [Incongruent vs Congruent] | 1.52 | 0.20 | 3.16 | **2.45e-03** | -0.01 | 0.01 | -1.25 | **2.94e-01** |
| Relation [Unrelated vs Congruent] | 7.40 | 1.64 | 9.02 | **1.32e-18** | -0.04 | 0.01 | -3.09 | **3.95e-03** |
| Group [Older vs Younger] | 0.51 | 0.07 | -4.65 | **6.49e-06** | 0.18 | 0.03 | 5.04 | **1.15e-06** |
| Task order [Them_Scram_Tax vs Tax_Scram_Them] | 1.00 | 0.13 | 0.03 | 9.73e-01 | 0.05 | 0.03 | 1.64 | 1.56e-01 |
| Gender [Male vs Female] | 0.86 | 0.11 | -1.20 | 2.91e-01 | 0.03 | 0.03 | 0.79 | 4.64e-01 |
| Task [Thematic] x Relation [Incongruent] | 0.38 | 0.05 | -7.26 | **1.11e-12** | -0.06 | 0.01 | -7.35 | **9.69e-13** |
| Task [Thematic] x Relation [Unrelated] | 0.15 | 0.03 | -8.41 | **1.42e-16** | 0.07 | 0.01 | 8.25 | **1.17e-15** |
| Task [Thematic] x Group [Older] | 0.99 | 0.20 | -0.07 | 9.73e-01 | 0.03 | 0.02 | 1.13 | 3.02e-01 |
| Relation [Incongruent] x Group [Older] | 1.75 | 0.41 | 2.38 | **2.44e-02** | -0.02 | 0.02 | -1.15 | 3.02e-01 |

| | | | | | | | | |
|---|---|---|---|---|---|---|---|---|
| Relation [Unrelated] x Group [Older] | 1.27 | 0.52 | 0.59 | 6.48e-01 | -0.01 | 0.02 | -0.57 | 5.66e-01 |
| Task [Thematic] x Relation [Incongruent] x Group [Older] | 5.98 | 1.56 | 6.88 | **1.44e-11** | -0.08 | 0.02 | -5.03 | **1.15e-06** |
| Task [Thematic] x Relation [Unrelated] x Group [Older] | 4.17 | 1.76 | 3.38 | **1.27e-03** | -0.05 | 0.02 | -2.93 | **5.94e-03** |
| Marginal $R^2$ / Conditional $R^2$ | 0.209 / 0.458 | | | | 0.166 / 0.478 | | | |

**Model formulas for scrambled items:**

| Accuracy | Reaction times |
|---|---|
| m_acc_scr <- glmmTMB(Accuracy ~<br>Condition +<br>Group +<br>Task_order +<br>Gender +<br>Condition : Group +<br>(1 + Condition \| Participant) +<br>(1 \| Trial),<br>data = df_acc_scr,<br>family = binomial(link = "logit")) | m_RT_scr <- lmer(log(RT) ~<br>Condition +<br>Group +<br>Task_order +<br>Gender +<br>Condition : Group +<br>(1 + Condition \| Participant) +<br>(1 \| Trial),<br>data = df_RT_scr,<br>REML = TRUE) |

**Table S2. Results from mixed-effects models for accuracy and reaction times in the scrambled images task.**

| | Accuracy | | | | Reaction times | | | |
|---|---|---|---|---|---|---|---|---|
| *Predictors* | *Odds Ratios* | *std. Error* | *Statistic* | *p* | *Estimates* | *std. Error* | *Statistic* | *p* |
| Condition [Rotated] | 1.22 | 0.31 | 0.79 | 4.28e-01 | -0.18 | 0.02 | -7.12 | **3.66e-12** |
| Group [Old] | 1.52 | 0.49 | 1.29 | 2.96e-01 | 0.05 | 0.05 | 1.04 | 4.49e-01 |
| Task order [Them_Scram_Tax] | 0.82 | 0.18 | -0.88 | 4.28e-01 | 0.01 | 0.04 | 0.17 | 8.68e-01 |
| Gender [M] | 1.42 | 0.31 | 1.57 | 2.34e-01 | -0.03 | 0.04 | -0.75 | 5.46e-01 |
| Condition [Rotated] × Group [Old] | 0.33 | 0.11 | -3.45 | **1.65e-03** | 0.09 | 0.04 | 2.50 | **2.48e-02** |
| Marginal $R^2$ / Conditional $R^2$ | 0.032 / 0.289 | | | | 0.078 / 0.488 | | | |

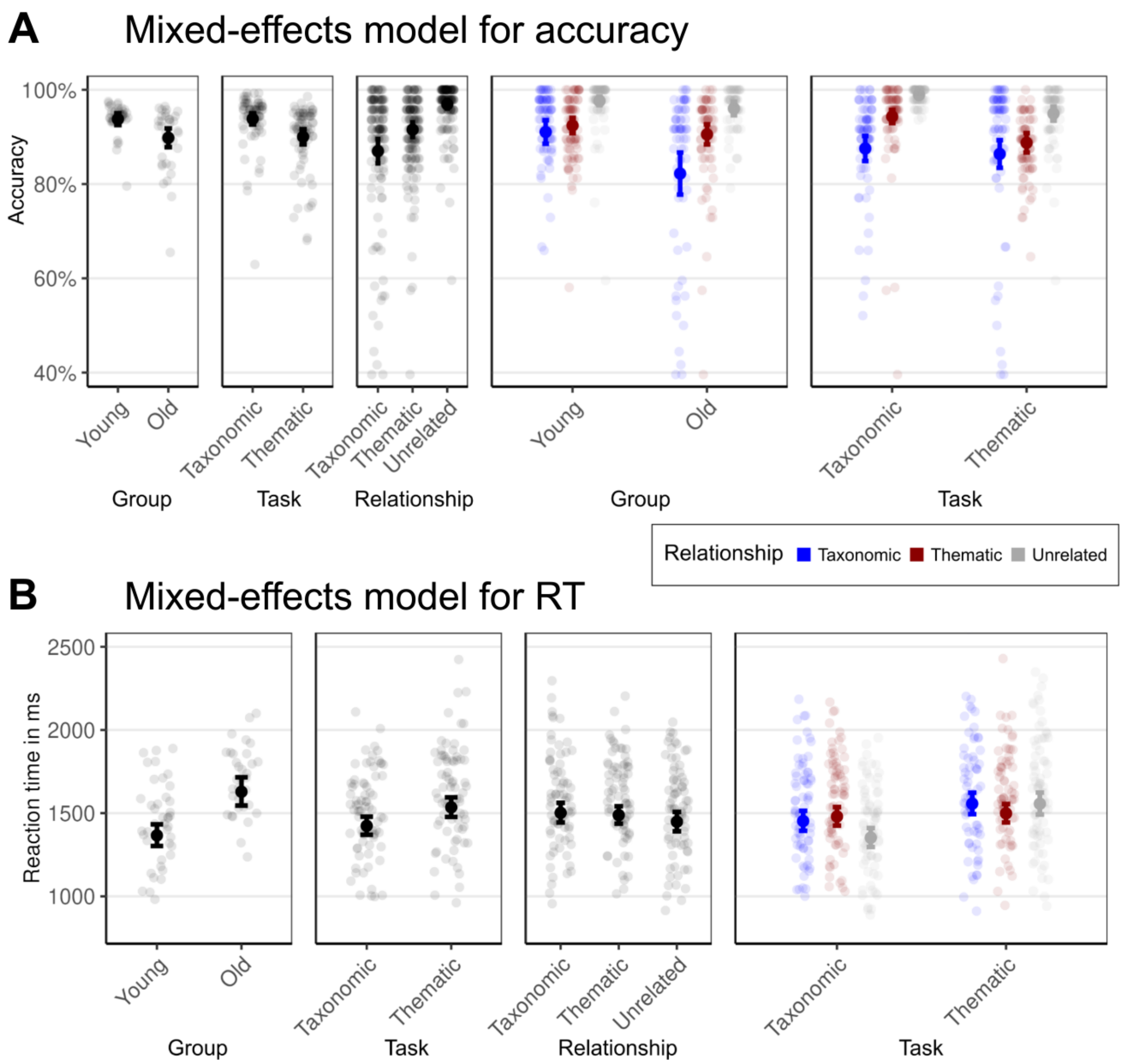

**Figure S1.** Significant effects in mixed-effects models for accuracy and reaction times.

#### Whole-Brain Activation Analyses

**Table S3. Whole-brain activation for semantic vs. non-semantic tasks (congruent only) in young adults.** Coordinates are in MNI space.

| Region | Cluster size ($mm^3$) | x | y | z | T |
|---|---|---|---|---|---|
| *Semantic > Non-Semantic Tasks* | | | | | |
| L IFG/MFG, pMTG, L/R pFG, mPFC, PreCun / PCC, Cerebellum | 342375 | | | | |
| L FOP (OP9) | | -46 | 25 | 12 | 9.59 |
| L IFGtri (45) | | -46 | 32 | 21 | 9.02 |
| L SFS (6d3) | | -31 | 15 | 51 | 8.82 |
| L pFG (FG3) | | -36 | -37 | -23 | 8.76 |
| L IFGtri (45) | | -51 | 30 | 15 | 8.69 |
| R pFG (FG4) | | 36 | -42 | -23 | 8.44 |
| L pFG (FG3) | | -31 | -42 | -23 | 8.42 |
| L IPS (hIP6) | | -41 | -62 | 43 | 8.18 |
| L MFG | | -29 | 17 | 54 | 8.16 |
| L pMTG | | -59 | -42 | -10 | 8.14 |
| R AG, pMTG, LOC (hOc4la) | 19922 | | | | |
| R AG | | 43 | -57 | 34 | 6.18 |
| R AG (PGa) | | 56 | -62 | 40 | 6.14 |
| R AG (PGp) | | 58 | -67 | 29 | 5.89 |
| R AG (PGa) | | 46 | -62 | 51 | 5.77 |
| R AG (PGp) | | 41 | -72 | 48 | 5.46 |
| R AG (PGp) | | 51 | -72 | 40 | 5.29 |
| R LOC (hOc4la) | | 51 | -77 | -4 | 4.94 |
| R IPS (hIP6) | | 38 | -65 | 43 | 4.61 |
| R pMTG | | 66 | -40 | -10 | 4.52 |
| R LOCsup | | 61 | -65 | 18 | 4.49 |
| R Parietal Operculum | | | | | |
| R ParOper (PFcm) | | 58 | -27 | 21 | 4.04 |
| R Insula (Ig2) | | 38 | -17 | 12 | 3.95 |
| R ParOper (PFcm) | | 48 | 32 | 21 | 3.72 |
| R S2 | | 56 | -22 | 15 | 3.64 |
| R ParOper (OP3) | | 46 | -5 | 10 | 3.23 |
| R ParOper (OP3) | | 48 | -3 | 12 | 3.20 |
| R Insula (Ig2) | | 41 | -15 | 1 | 3.08 |
| *Non-Semantic > Semantic Tasks* | | | | | |
| R IPS / SPL | 25688 | | | | |
| R SPL (7P) | | 16 | -67 | 54 | 8.37 |
| R SPL (7A) | | 21 | -62 | 51 | 8.06 |
| R IPS (hIP7) | | 31 | -77 | 23 | 8.02 |
| R SPL (7P) | | 23 | -75 | 51 | 7.81 |
| R IPS (hIP3) | | 23 | -57 | 54 | 7.60 |
| R LOC (hOc4lp) | | 36 | -85 | 18 | 7.07 |
| R IPS (hIP8) | | 26 | -70 | 37 | 7.01 |
| R IPS (hIP2) | | 41 | -40 | 45 | 6.54 |
| R IPS (hIP2) | | 43 | -35 | 43 | 6.36 |
| R SPL (7P) | | 13 | -67 | 67 | 5.56 |
| L IPS / SPL | 3359 | | | | |
| L SPL (7A) | | -17 | -67 | 56 | 6.34 |
| L SPL (7A) | | -17 | -62 | 54 | 5.94 |
| L IPS (hIP8) | | -14 | -72 | 48 | 5.63 |
| L IPS (hIP3) | | -22 | -60 | 54 | 5.53 |

| | | | | | |
|---|---|---|---|---|---|
| L SPL (7A) | | -12 | -72 | 54 | 5.51 |
| L IPS (hIP1) | | -29 | -45 | 43 | 5.46 |
| L IPS (hIP3) | | -26 | -55 | 54 | 5.06 |
| L LOC (hOc4lp), IPS | 3344 | | | | |
| L IPS (hIP4) | | -26 | -77 | 21 | 6.83 |
| L LOC (hOc4lp) | | -29 | -90 | 18 | 5.59 |
| R SFS (6d3) | 1219 | | | | |
| R SFS (6d3) | | 28 | -3 | 48 | 7.18 |
| R SFS (6d3) | | 23 | 5 | 48 | 7.04 |

**Table S4. Whole-brain activation for semantic vs. non-semantic tasks (congruent only) in older adults.** Coordinates are in MNI space.

| Region | Cluster size (mm³) | x | y | z | T |
|---|---|---|---|---|---|
| *Semantic > Non-Semantic Tasks* | | | | | |
| L IFG, OFC, L/R mPFC | 86063 | | | | |
| L OFC (Fo7) | | -46 | 35 | 15 | 10.00 |
| L mPFC (Fp2) | | -4 | 62 | 1 | 9.55 |
| L IFGorb | | -49 | 27 | -10 | 9.16 |
| L mPFC (Fp2) | | -7 | 67 | 7 | 8.75 |
| L dmPFC | | -7 | 45 | 48 | 8.30 |
| L vmPFC (s32) | | -4 | 35 | -18 | 8.01 |
| L vmPFC (s32) | | -7 | 40 | -15 | 7.94 |
| L dmPFC | | -9 | 52 | 40 | 7.94 |
| L IFGtri (45) | | -54 | 27 | -1 | 7.79 |
| L OFC (Fo7) | | -46 | 37 | -12 | 7.78 |
| L AG, STS / MTG / ITG | 49391 | | | | |
| L IPL (PFm) | | -44 | -62 | 29 | 10.10 |
| L AG (PGp) | | -51 | -67 | 29 | 9.53 |
| L AG (PGp) | | -41 | 72 | 40 | 7.71 |
| L pMTG | | -61 | -50 | 4 | 6.86 |
| L aMTG (TE 5) | | -61 | -15 | -15 | 6.29 |
| L aSTS (TE 5) | | -56 | -10 | -12 | 6.23 |
| L pMTG | | -56 | -47 | -1 | 5.87 |
| L pITG | | -56 | -45 | -15 | 5.79 |
| L pMTG | | -59 | -37 | -12 | 5.69 |
| L mSTS (TE 4) | | -51 | -22 | -12 | 5.35 |
| L/R PCC / PreCun | 16578 | | | | |
| L PCC | | -2 | -47 | 29 | 9.44 |
| L PCC | | -4 | -52 | 18 | 9.43 |
| L PCC | | -2 | -55 | 26 | 9.09 |
| R PCC | | 1 | -15 | 40 | 6.85 |
| L PreCun | | -2 | -62 | 34 | 6.78 |
| R PreCun | | 3 | -65 | 37 | 6.56 |
| L CC | | -7 | -3 | 40 | 3.42 |
| R Cerebellum (Crus I / II) | 8406 | | | | |
| R Cerebellum (Crus II) | | 38 | -75 | -40 | 9.56 |
| R Cerebellum (Crus II) | | 33 | -77 | -40 | 9.41 |
| R Cerebellum (Crus II) | | 21 | -85 | -40 | 6.46 |
| R Cerebellum (Crus II) | | 13 | -82 | -34 | 6.35 |
| R Cerebellum (Crus II) | | 11 | -87 | -32 | 5.13 |
| R Cerebellum (Crus I) | | 18 | -72 | -29 | 4.10 |
| R AG | 5016 | | | | |
| R AG (PGp) | | 51 | -62 | 26 | 6.43 |
| R AG (PGp) | | 58 | -65 | 18 | 6.15 |
| R AG (PGa) | | 53 | -62 | 43 | 5.77 |

| | | | | | |
|---|---|---|---|---|---|
| R AG (PGa) | | 63 | -57 | 15 | 5.60 |
| R AG (PGp) | | 46 | -72 | 45 | 4.85 |
| *Non-Semantic > Semantic Tasks* | | | | | |
| L/R IPS / SPL, LOC (hOc4lp), Cerebellum | 233281 | | | | |
| R IPS (hIP3) | | 26 | -60 | 51 | 11.30 |
| L IPS (hIP6) | | -24 | -65 | 54 | 10.90 |
| R SPL (7P) | | 18 | -72 | 54 | 10.70 |
| R IPS (hIP1) | | 38 | -47 | 43 | 10.50 |
| R IPS (hIP8) | | 23 | -70 | 45 | 10.50 |
| R IPS (hIP4) | | 36 | -77 | 29 | 10.20 |
| L hOc4d (V3A) | | -26 | -87 | 29 | 9.60 |
| R SPL (7A) | | 23 | -65 | 56 | 9.41 |
| R SPL (7P) | | 8 | -72 | 56 | 9.32 |
| R IPS (hIP3) | | 28 | -52 | 48 | 9.25 |
| R Frontal Pole | 4063 | | | | |
| R FP | | 36 | 47 | 29 | 6.31 |
| R FP | | 41 | 45 | 23 | 4.98 |
| R FP | | 36 | 40 | 26 | 4.85 |
| R FP | | 41 | 40 | 29 | 4.78 |
| R FP | | 38 | 40 | 40 | 4.27 |
| R FP | | 36 | 55 | 18 | 3.26 |
| L Thalamus / Caudate | 469 | | | | |
| L Thalamus | | -12 | -15 | 18 | 4.28 |
| L Caudate | | -17 | -22 | 21 | 3.68 |
| L Thalamus | | -12 | -20 | 10 | 3.62 |

**Table S5. Increased whole-brain activation for semantic vs. non-semantic tasks (congruent only) in young vs. older adults.** Coordinates are in MNI space.

| Region | Cluster size (mm$^3$) | x | y | z | T |
|---|---|---|---|---|---|
| *Young > older adults* | | | | | |
| L pFG / pITG, Hippocampus, Cerebellum | 9547 | | | | |
| L pFG (FG3) | | -34 | -42 | -21 | 4.86 |
| L pFG (FG3) | | -36 | -30 | -23 | 4.61 |
| L pFG (FG3) | | -34 | -52 | -7 | 4.48 |
| L Hippocampus (CA1) | | -39 | -17 | -21 | 4.09 |
| L Cerebellum (Lobule V) | | -9 | -50 | -7 | 4.01 |
| L Hippocampus (CA1) | | -17 | -35 | 1 | 3.85 |
| L Cerebellum (Lobule I-IV) | | -4 | -50 | -4 | 3.78 |
| L pITG | | -54 | -55 | -15 | 3.64 |
| L pFG (FG4) | | -41 | -57 | -7 | 3.54 |
| L Hippocampus (CA1) | | -34 | -37 | -12 | 3.43 |
| L IPS / SPL | 6531 | | | | |
| L IPS (hIP1) | | -41 | -47 | 45 | 5.41 |
| L IPS (hIP3) | | -31 | -55 | 43 | 4.92 |
| L IPS (hIP6) | | -39 | -60 | 43 | 4.85 |
| L IPS (hIP6) | | -26 | -67 | 45 | 4.23 |
| L IPL (PFm) | | -46 | -52 | 56 | 3.95 |
| L SPL (7A) | | -39 | -62 | 59 | 3.83 |
| L SPL (7A) | | -34 | -65 | 62 | 3.48 |

| | | | | | |
|---|---|---|---|---|---|
| L SPL (7A) | | -29 | -67 | 62 | 3.45 |
| L IPS (hIP1) | | -44 | -45 | 32 | 3.31 |
| L IPL (PF) | | -56 | -45 | 54 | 3.19 |
| L IFGtri / MFG | 2016 | | | | |
| L MFG | | -44 | 35 | 18 | 4.94 |
| L MFG | | -41 | 42 | 29 | 3.68 |
| L IFGtri (45) | | -51 | 37 | 12 | 3.36 |
| L Cerebellum, V2 / V3 | 1750 | | | | |
| L hOc2 (V2) | | -7 | -70 | -4 | 4.74 |
| L Cerebellum (Lobule VI) | | -19 | -60 | -18 | 3.52 |
| L Cerebellum (Lobule V) | | -12 | -55 | -21 | 2.92 |
| L Cerebellum (Lobule V) | | -14 | -50 | -18 | 2.50 |
| L/R pre-SMA / Paracingulate | 1359 | | | | |
| R Paracingulate | | 3 | 30 | 43 | 4.76 |
| L pre-SMA | | -4 | 22 | 45 | 4.21 |
| L medial SFG | | -9 | 25 | 54 | 4.00 |
| L/R V2 / V3 | 547 | | | | |
| R hOc3d (V3d) | | 11 | -87 | 23 | 3.76 |
| L hOc4d (V3A) | | -7 | -90 | 21 | 3.43 |
| R hOc2 (V2) | | 1 | -87 | 18 | 3.37 |
| R Cerebellum (Crus I / II) | 188 | | | | |
| R Cerebellum (Crus II) | | 11 | -75 | -32 | 3.03 |
| R Cerebellum (Crus II) | | 11 | -70 | -34 | 2.98 |
| R IPS (hIP6) | 188 | 36 | -67 | 45 | 4.70 |
| *Older > young adults* | | | | | |
| No significant activations | | | | | |

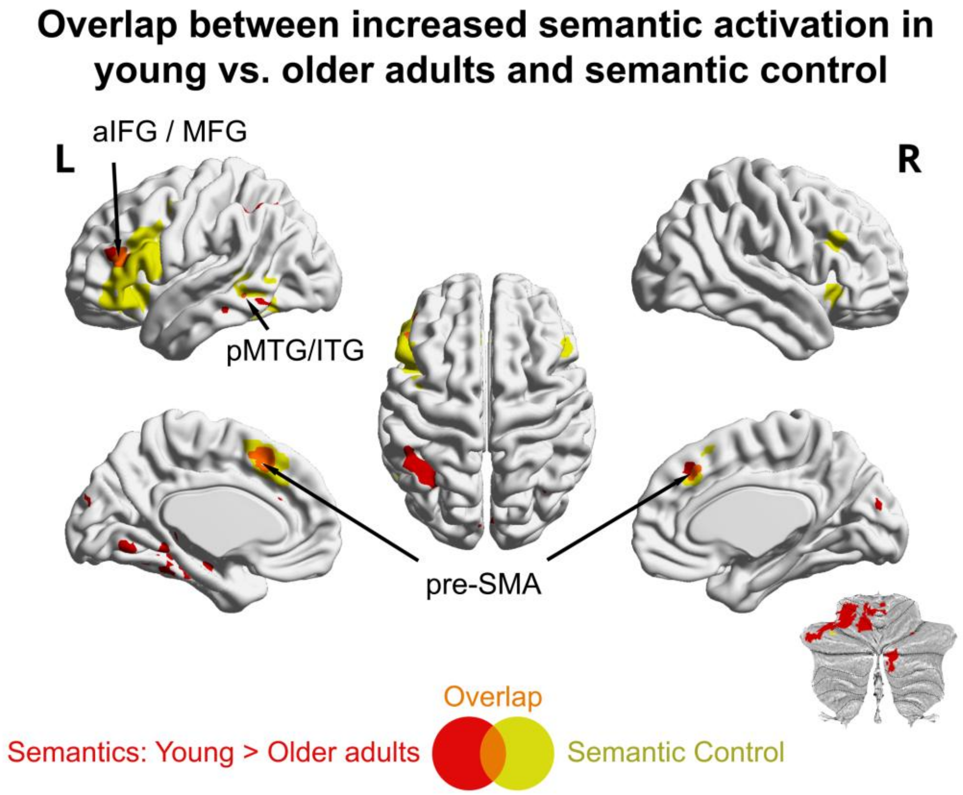

**Figure S2. Conjunction between stronger activation for semantic > non-semantic tasks in young than older adults and the semantic control system.** The map of semantic control is derived from the neuroimaging meta-analysis by Jackson (2021)**.**

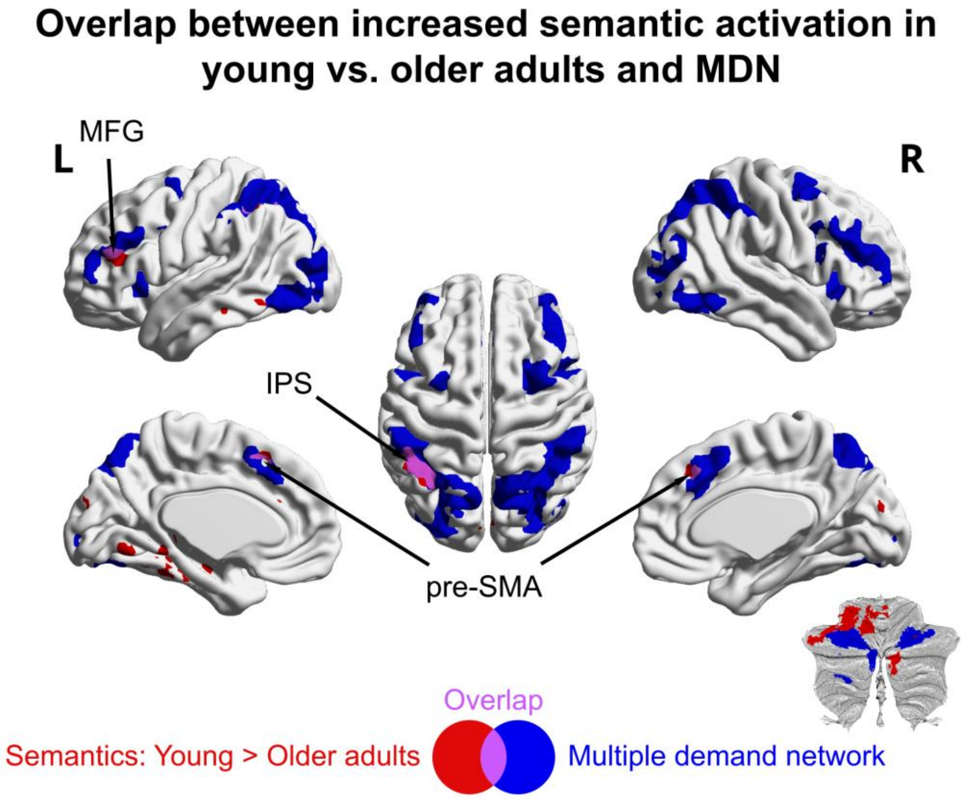

**Figure S3. Conjunction between stronger activation for semantic > non-semantic tasks in young than older adults and the multiple demand network (MDN).** The MDN map is derived from the probabilistic map of Lipkin et al. (2022), thresholded at 30% probability.

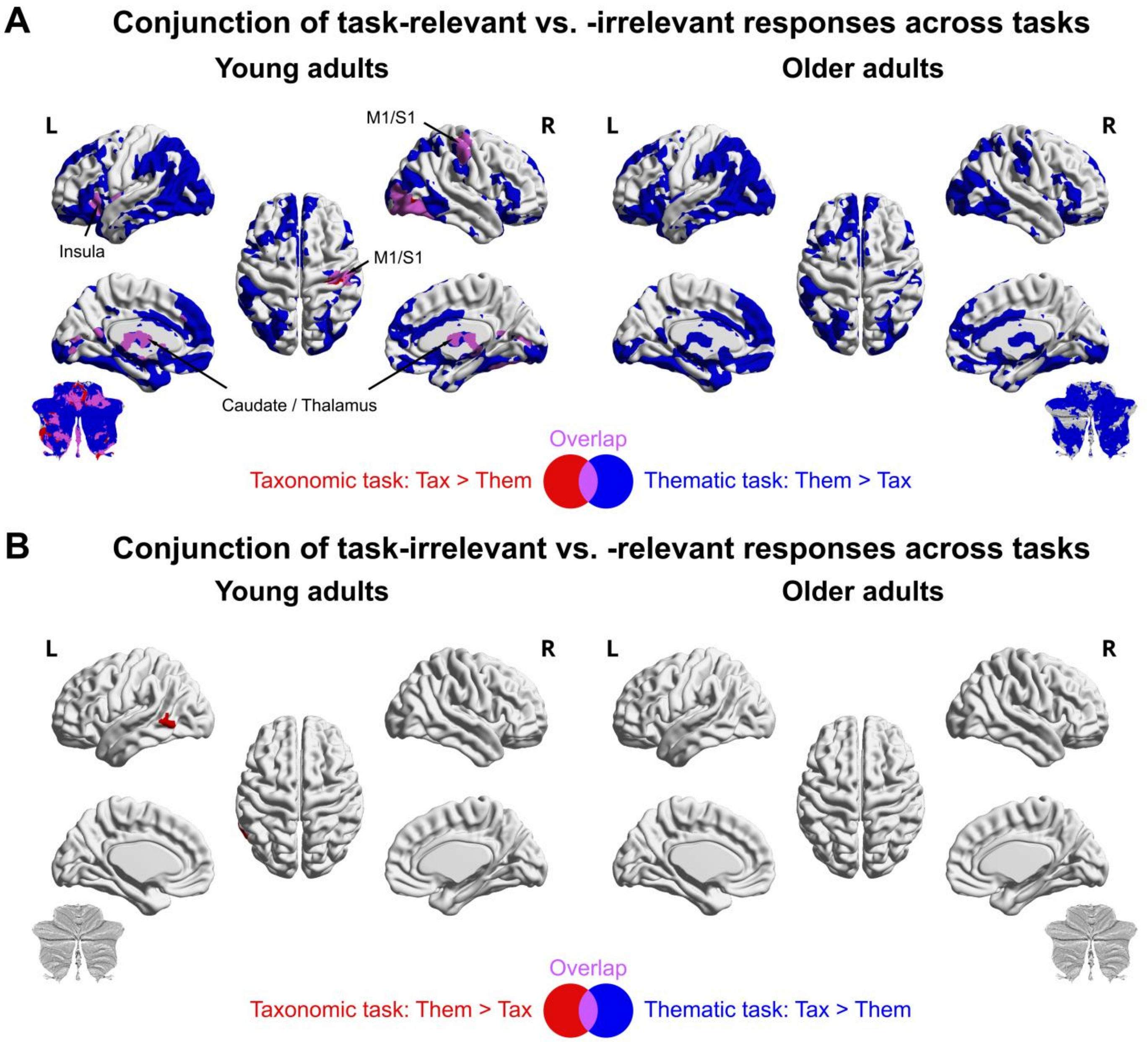

**Figure S4. Conjunction of task-relevant ("yes") vs. task-irrelevant ("no") responses across tasks.** All activation maps were thresholded at $p < 0.05$ FWE-corrected using threshold free cluster enhancement (TFCE).

#### ROI Analyses

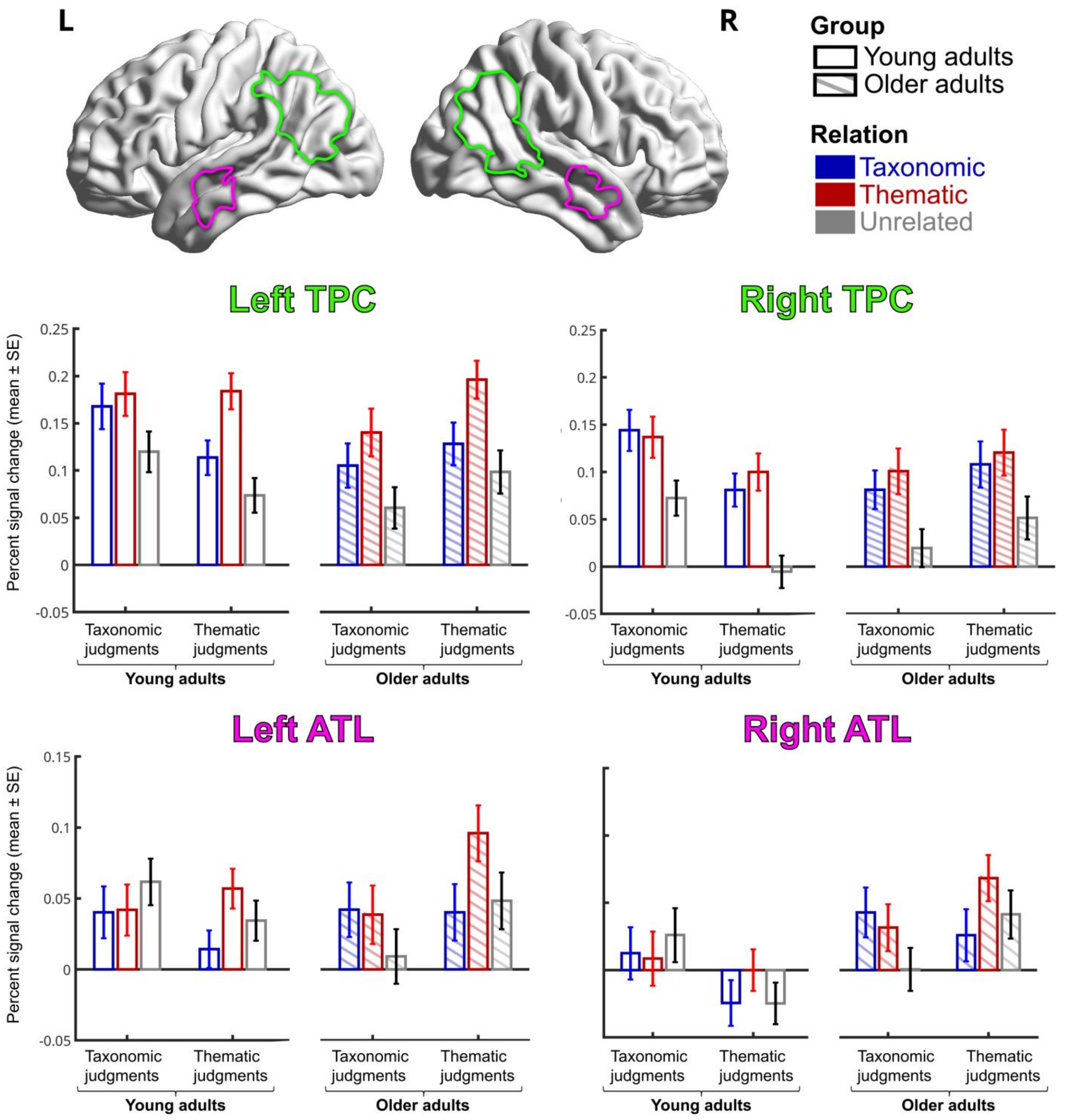

**Figure S5. Complete response profiles for subject-specific fROIs in the left and right TPC and ATL.** Mean percent signal change is shown for each experimental condition; error bars represent the standard error of the mean.

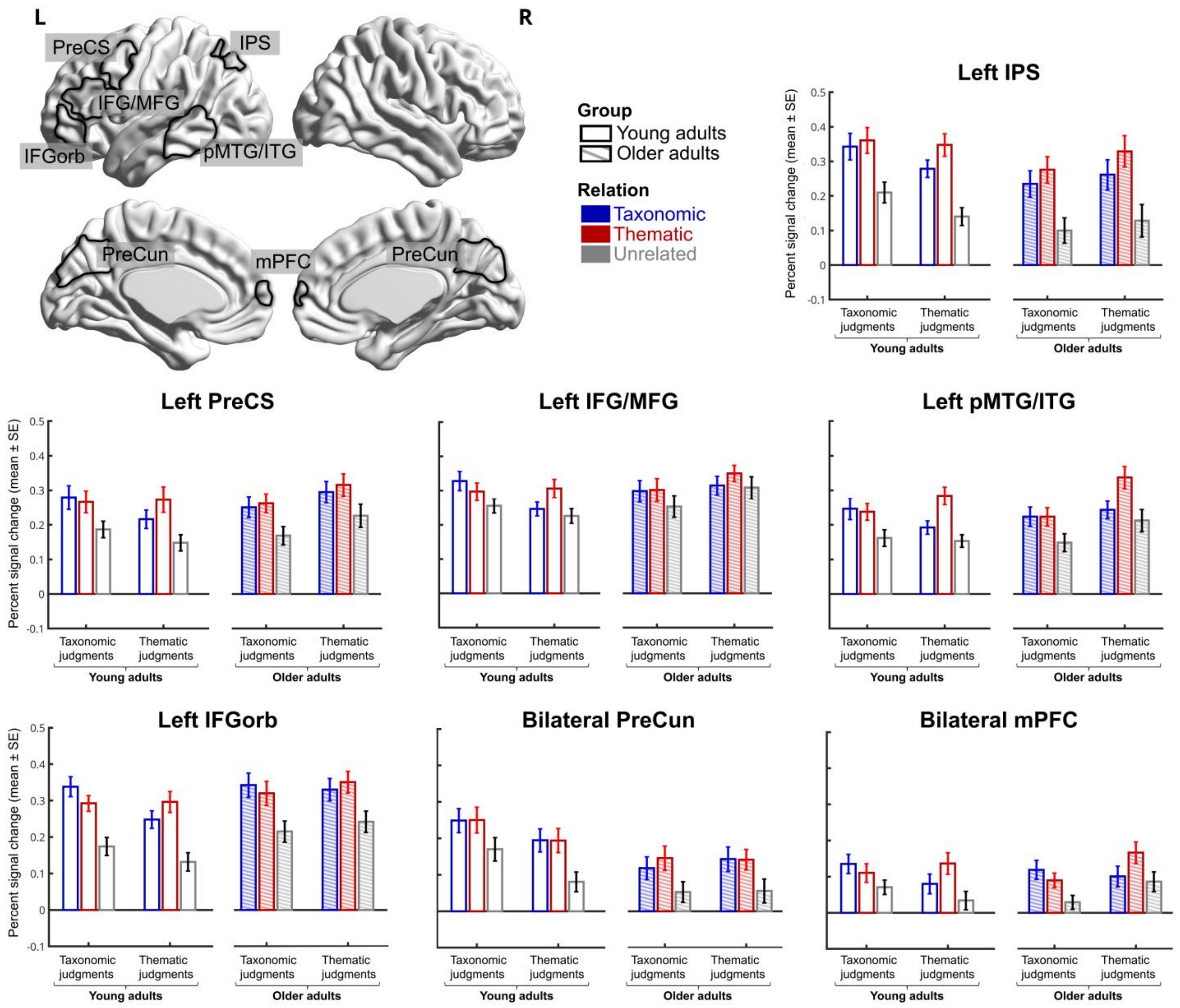

**Figure S6. Complete response profiles for subject-specific fROIs in semantic regions beyond TPC and ATL.** Mean percent signal change is shown for each experimental condition; error bars represent the standard error of the mean.

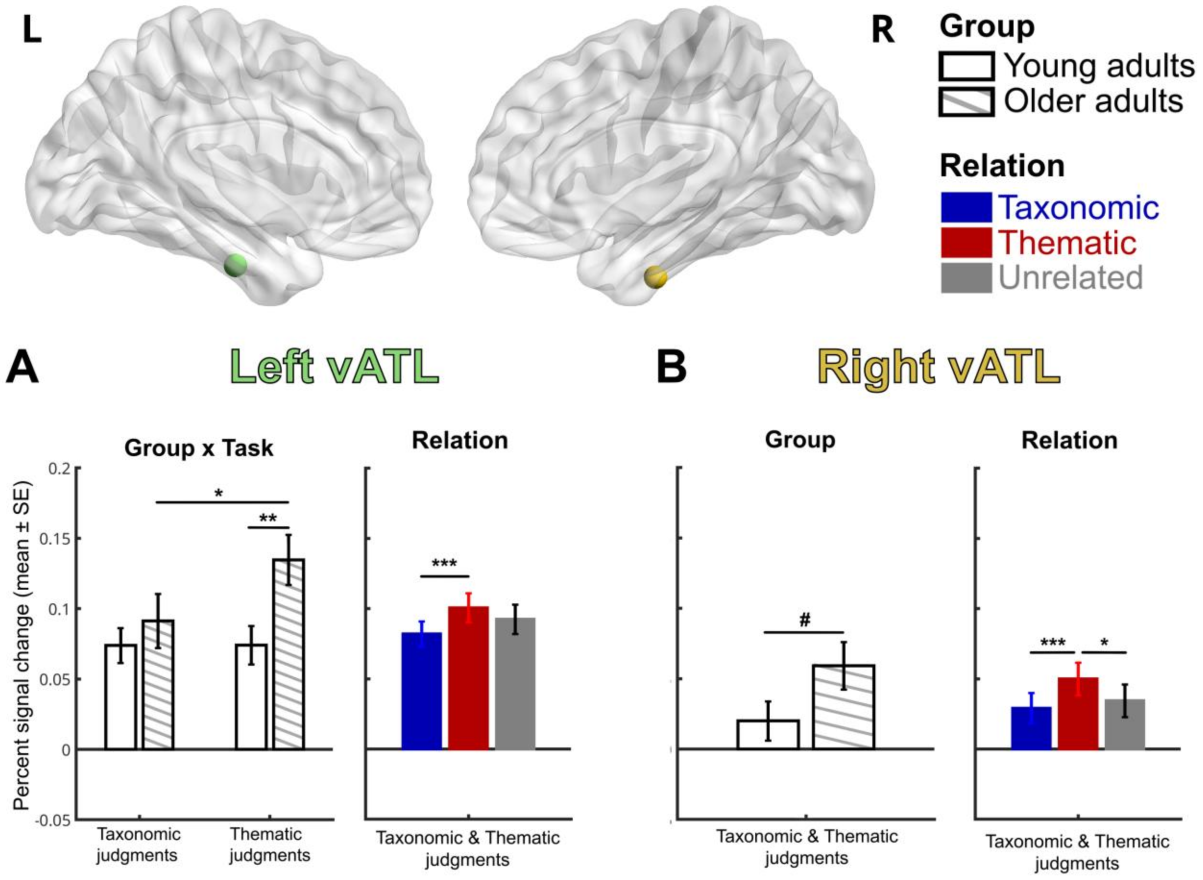

**Figure S7. Response profiles of the ventral ATLs.** ROIs were defined as 5-mm radius spheres around the peak coordinates for semantic vs. non-semantic tasks across eight distortion-corrected fMRI studies (Gonzalez Alam et al., 2021; Rice et al., 2018). ***p < 0.001 (corr.), **p < 0.01 (corr.), *p < 0.05 (corr.), $^{\#}p <$ 0.05 (uncorr.).

Coordinates:

- left vATL: MNI x = -41, y = -15, z = -31
- right vATL: MNI x = 44, y = -11, z = -36

##### Left ventral ATL (vATL)

Left vATL (Figure S7A; Tables S27-S31) exhibited a trend towards a GROUP x TASK interaction ($F_{1,73}$ = 3.385, p = 0.07, $\eta^2_p$ = 0.044), reflecting stronger activity for thematic than taxonomic judgments in older adults (t = 2.502, p = 0.008, Cohen's d = 0.425) and higher activity for thematic judgments in older than young adults (t = 2.742, p = 0.004, Cohen's d = 0.593). A main effect of RELATION ($F_{2,146}$ = 6.900, p < 0.001, $\eta^2_p$ = 0.086) further revealed stronger activity for thematic than taxonomic relationships across tasks and age groups (t = 4.147, p < 0.001, Cohen's d = 0.182).

##### Right ventral ATL (vATL)

Right vATL (Figure S7B; Tables S32-S35) showed a trend towards a GROUP main effect ($F_{1,73}$ = 3.291, p = 0.074, $\eta^2_p$ = 0.043), driven by generally stronger activation in older than young adults (t = 1.814, p = 0.074, Cohen's d = 0.325). Moreover, a RELATION main effect ($F_{2,146}$ = 6.900, p < 0.001, $\eta^2_p$ = 0.086) revealed stronger activity for thematic than taxonomic relationships (t = 4.028, p < 0.001, Cohen's d = 0.173) and for thematic than unrelated pairs (t = 2.408, p = 0.019, Cohen's d = 0.128).

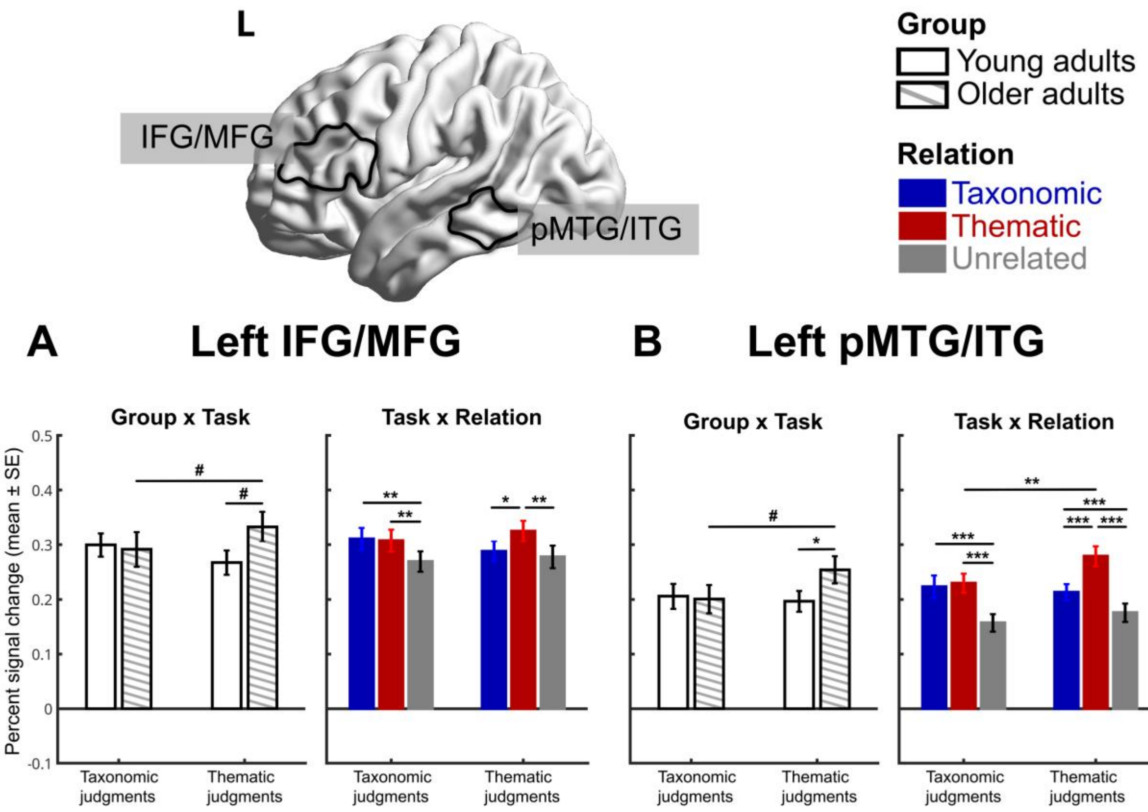

**Figure S8. Response profiles of two key semantic control fROIs when defined using the semantic > non-semantic task contrast including all conditions.** Mean percent signal change is shown for each experimental condition; error bars represent the standard error of the mean. ***p < 0.001 (corr.), **p < 0.01 (corr.), *p < 0.05 (corr.), #p < 0.05 (uncorr.).

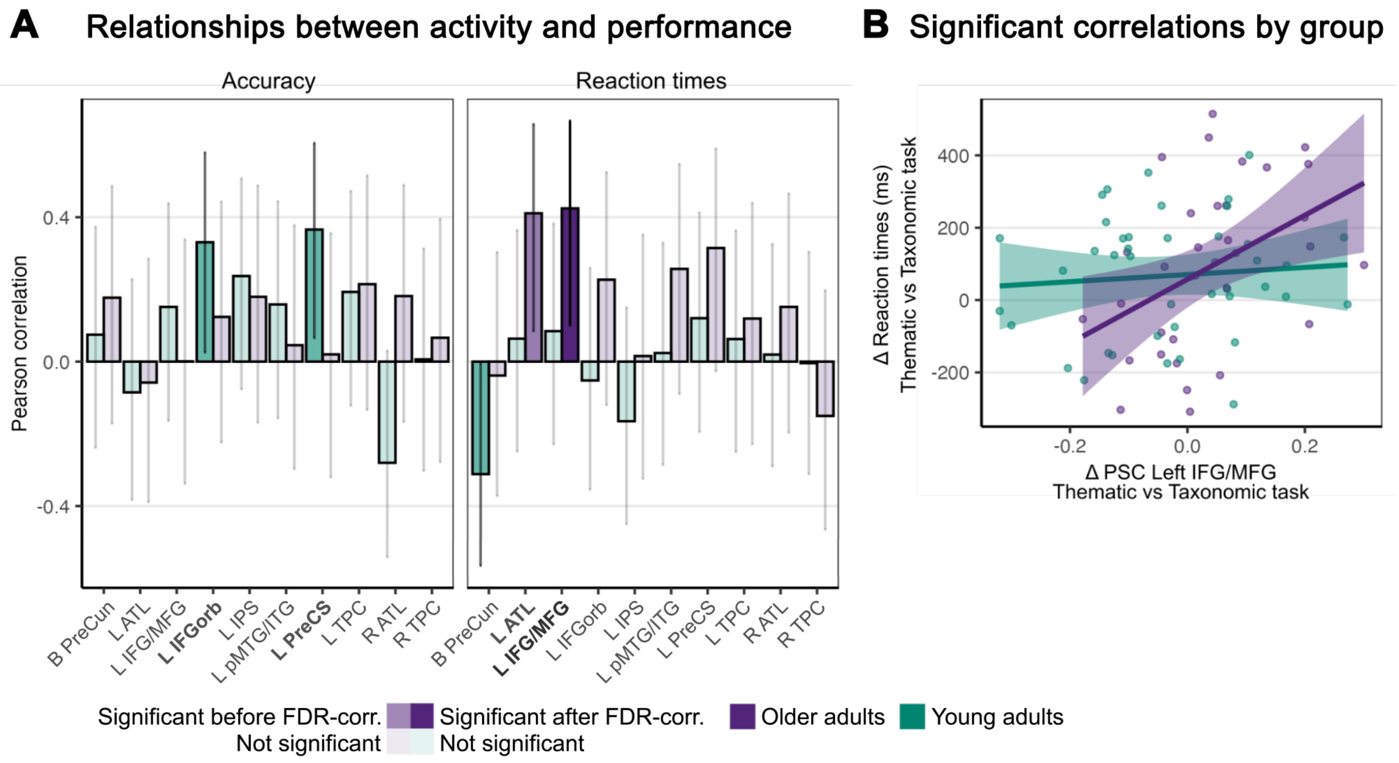

**Figure S9. Brain–behavior relationships between task-related activity and behavioral performance for task-relevant items only.** (A) Correlation strength of percent signal change in fROIs and behavioral performance by age group. (B) Significant correlations after FDR correction.

### ROI Analysis Statistics

#### ROI 1: Left TPC

##### Repeated Measures ANOVA

*Table S6. Within Subjects Effects*

| Cases | Sphericity Correction | Sum of Squares | df | Mean Square | F | p | $\eta^2_p$ |
|---|---|---|---|---|---|---|---|
| Task | None | 0.001 | 1.000 | 0.001 | 0.057 | .812 | $7.769 \times 10^{-4}$ |
| Task ✻ Group | None | 0.143 | 1.000 | 0.143 | 6.901 | .010 | 0.086 |
| Residuals | None | 1.508 | 73.000 | 0.021 | | | |
| Relation | None | 0.567[a] | 2.000[a] | 0.284[a] | 88.095[a] | < .001[a] | 0.547 |
| | Huynh-Feldt | 0.567 | 1.754 | 0.323 | 88.095 | < .001 | 0.547 |
| Relation ✻ Group | None | 0.002[a] | 2.000[a] | $9.207 \times 10^{-4}$[a] | 0.286[a] | .752[a] | 0.004 |
| | Huynh-Feldt | 0.002 | 1.754 | 0.001 | 0.286 | .723 | 0.004 |
| Residuals | None | 0.470 | 146.000 | 0.003 | | | |
| | Huynh-Feldt | 0.470 | 128.016 | 0.004 | | | |
| Task ✻ Relation | None | 0.041 | 2.000 | 0.020 | 7.350 | < .001 | 0.091 |
| | Huynh-Feldt | 0.041 | 1.929 | 0.021 | 7.350 | .001 | 0.091 |
| Task ✻ Relation ✻ Group | None | 0.005 | 2.000 | 0.002 | 0.892 | .412 | 0.012 |
| | Huynh-Feldt | 0.005 | 1.929 | 0.003 | 0.892 | .409 | 0.012 |
| Residuals | None | 0.405 | 146.000 | 0.003 | | | |
| | Huynh-Feldt | 0.405 | 140.786 | 0.003 | | | |

*Note.* Sphericity corrections not available for factors with 2 levels.
*Note.* Type III Sum of Squares
[a] Mauchly's test of sphericity indicates that the assumption of sphericity is violated (p < .05).

*Table S7. Between Subjects Effects*

| Cases | Sum of Squares | df | Mean Square | F | p | $\eta^2_p$ |
|---|---|---|---|---|---|---|
| Group | 0.038 | 1 | 0.038 | 0.518 | .474 | 0.007 |
| Residuals | 5.385 | 73 | 0.074 | | | |

*Note.* Type III Sum of Squares

##### Post Hoc Tests

*Table S8. Post Hoc Comparisons - Group ✻ Task - Conditional on Group*

| Group | | | Mean Difference | SE | df | t | Cohen's d | $p_{bonf}$ | $p_{holm}$ |
|---|---|---|---|---|---|---|---|---|---|
| 1 | TaxTask | ThemTask | 0.033 | 0.018 | 73 | 1.774 | 0.244 | .080 | .080 |
| 2 | | ThemTask | -0.039 | 0.020 | 73 | -1.938 | -0.293 | .057 | .057 |

*Note.* Results are averaged over the levels of: Relation

*Table S9. Post Hoc Comparisons - Group ✻ Task - Conditional on Task*

| Task | | | Mean Difference | SE | df | t | Cohen's d | $p_{bonf}$ | $p_{holm}$ |
|---|---|---|---|---|---|---|---|---|---|
| TaxTask | GroupY | GroupO | 0.054 | 0.031 | 73 | 1.727 | 0.408 | .088 | .088 |
| ThemTask | | GroupO | -0.017 | 0.027 | 73 | -0.648 | -0.129 | .519 | .519 |

*Note.* Results are averaged over the levels of: Relation

*Table S10. Post Hoc Comparisons - Task ✻ Relation - Conditional on Task*

| Task | | | Mean Difference | SE | df | t | Cohen's d | $p_{bonf}$ | $p_{holm}$ |
|---|---|---|---|---|---|---|---|---|---|
| TaxTask | Tax | Them | -0.024 | 0.007 | 73 | -3.398 | -0.181 | .003 | .001 |
| | | Unrel | 0.046 | 0.010 | 73 | 4.449 | 0.349 | < .001 | < .001 |
| | Them | Unrel | 0.071 | 0.009 | 73 | 7.497 | 0.529 | < .001 | < .001 |
| ThemTask | Tax | Them | -0.069 | 0.010 | 73 | -7.281 | -0.520 | < .001 | < .001 |
| | | Unrel | 0.035 | 0.008 | 73 | 4.214 | 0.262 | < .001 | < .001 |
| | Them | Unrel | 0.104 | 0.009 | 73 | 11.813 | 0.781 | < .001 | < .001 |

*Note.* P-value adjusted for comparing a family of 3 estimates.
*Note.* Results are averaged over the levels of: Group

*Table S11. Post Hoc Comparisons - Task ✻ Relation - Conditional on Relation*

| Relation | | | Mean Difference | SE | df | t | Cohen's d | $p_{bonf}$ | $p_{holm}$ |
|---|---|---|---|---|---|---|---|---|---|
| Tax | TaxTask | ThemTask | 0.016 | 0.017 | 73 | 0.928 | 0.118 | .356 | .356 |
| Them | | ThemTask | -0.029 | 0.014 | 73 | -2.093 | -0.221 | .040 | .040 |
| Unrel | | ThemTask | 0.004 | 0.015 | 73 | 0.273 | 0.031 | .785 | .785 |

*Note.* Results are averaged over the levels of: Group

### ROI 2: Right TPC

#### Repeated Measures ANOVA

*Table S12. Within Subjects Effects*

| Cases | Sphericity Correction | Sum of Squares | df | Mean Square | F | p | $\eta^2_p$ |
|---|---|---|---|---|---|---|---|
| Task | None | 0.031 | 1.000 | 0.031 | 1.645 | .204 | 0.022 |
| Task ✻ Group | None | 0.203 | 1.000 | 0.203 | 10.943 | .001 | 0.130 |
| Residuals | None | 1.357 | 73.000 | 0.019 | | | |
| Relation | None | 0.560[a] | 2.000[a] | 0.280[a] | 93.321[a] | < .001[a] | 0.561 |
| | Huynh-Feldt | 0.560 | 1.744 | 0.321 | 93.321 | < .001 | 0.561 |
| Relation ✻ Group | None | 0.007[a] | 2.000[a] | 0.004[a] | 1.216[a] | .299[a] | 0.016 |
| | Huynh-Feldt | 0.007 | 1.744 | 0.004 | 1.216 | .296 | 0.016 |
| Residuals | None | 0.438 | 146.000 | 0.003 | | | |
| | Huynh-Feldt | 0.438 | 127.342 | 0.003 | | | |
| Task ✻ Relation | None | 0.004 | 2.000 | 0.002 | 0.670 | .514 | 0.009 |
| | Huynh-Feldt | 0.004 | 1.899 | 0.002 | 0.670 | .506 | 0.009 |
| Task ✻ Relation ✻ Group | None | 0.013 | 2.000 | 0.007 | 2.242 | .110 | 0.030 |

*Table S12. Within Subjects Effects*

| Cases | Sphericity Correction | Sum of Squares | df | Mean Square | F | p | $\eta^2_p$ |
|---|---|---|---|---|---|---|---|
| | Huynh-Feldt | 0.013 | 1.899 | 0.007 | 2.242 | .113 | 0.030 |
| Residuals | None | 0.438 | 146.000 | 0.003 | | | |
| | Huynh-Feldt | 0.438 | 138.662 | 0.003 | | | |

*Note.* Sphericity corrections not available for factors with 2 levels.
*Note.* Type III Sum of Squares
[a] Mauchly's test of sphericity indicates that the assumption of sphericity is violated ($p < .05$).

*Table S13. Between Subjects Effects*

| Cases | Sum of Squares | df | Mean Square | F | p | $\eta^2_p$ |
|---|---|---|---|---|---|---|
| Group | 0.007 | 1 | 0.007 | 0.103 | .749 | 0.001 |
| Residuals | 4.982 | 73 | 0.068 | | | |

*Note.* Type III Sum of Squares

#### Post Hoc Tests

*Table S14. Post Hoc Comparisons - Group ✻ Task - Conditional on Group*

| Group | | | Mean Difference | SE | df | t | Cohen's d | $p_{bonf}$ | $p_{holm}$ |
|---|---|---|---|---|---|---|---|---|---|
| 1 | TaxTask | ThemTask | 0.059 | 0.017 | 73 | 3.409 | 0.462 | .001 | .001 |
| 2 | | ThemTask | -0.026 | 0.019 | 73 | -1.370 | -0.204 | .175 | .175 |

*Note.* Results are averaged over the levels of: Relation

*Table S15. Post Hoc Comparisons - Group ✻ Task - Conditional on Task*

| Task | | | Mean Difference | SE | df | t | Cohen's d | $p_{bonf}$ | $p_{holm}$ |
|---|---|---|---|---|---|---|---|---|---|
| TaxTask | Group1 | Group2 | 0.051 | 0.028 | 73 | 1.792 | 0.395 | .077 | .077 |
| ThemTask | | Group2 | -0.035 | 0.028 | 73 | -1.262 | -0.271 | .211 | .211 |

*Note.* Results are averaged over the levels of: Relation

*Table S16. Post Hoc Comparisons - Relation*

| | | Mean Difference | SE | df | t | Cohen's d | $p_{bonf}$ | $p_{holm}$ |
|---|---|---|---|---|---|---|---|---|
| Tax | Them | -0.011 | 0.005 | 73 | -2.256 | -0.086 | .081 | .027 |
| | Unrel | 0.069 | 0.007 | 73 | 9.701 | 0.538 | < .001 | < .001 |
| Them | Unrel | 0.080 | 0.007 | 73 | 11.729 | 0.624 | < .001 | < .001 |

*Note.* P-value adjusted for comparing a family of 3 estimates.
*Note.* Results are averaged over the levels of: Group, Task

#### ROI 3: Left ATL

##### Repeated Measures ANOVA

*Table S17. Within Subjects Effects*

| Cases | Sphericity Correction | Sum of Squares | df | Mean Square | F | p | $\eta^2_p$ |
|---|---|---|---|---|---|---|---|
| Task | None | 0.010 | 1.000 | 0.010 | 0.639 | .427 | 0.009 |
| Task ✻ Group | None | 0.055 | 1.000 | 0.055 | 3.546 | .064 | 0.046 |
| Residuals | None | 1.128 | 73.000 | 0.015 | | | |
| Relation | None | 0.049[a] | 2.000[a] | 0.025[a] | 15.460[a] | < .001[a] | 0.175 |
| | Huynh-Feldt | 0.049 | 1.880 | 0.026 | 15.460 | < .001 | 0.175 |
| Relation ✻ Group | None | 0.031[a] | 2.000[a] | 0.015[a] | 9.687[a] | < .001[a] | 0.117 |
| | Huynh-Feldt | 0.031 | 1.880 | 0.016 | 9.687 | < .001 | 0.117 |
| Residuals | None | 0.233 | 146.000 | 0.002 | | | |
| | Huynh-Feldt | 0.233 | 137.270 | 0.002 | | | |
| Task ✻ Relation | None | 0.047 | 2.000 | 0.024 | 14.434 | < .001 | 0.165 |
| | Huynh-Feldt | 0.047 | 1.980 | 0.024 | 14.434 | < .001 | 0.165 |
| Task ✻ Relation ✻ Group | None | 0.008 | 2.000 | 0.004 | 2.568 | .080 | 0.034 |
| | Huynh-Feldt | 0.008 | 1.980 | 0.004 | 2.568 | .081 | 0.034 |
| Residuals | None | 0.239 | 146.000 | 0.002 | | | |
| | Huynh-Feldt | 0.239 | 144.519 | 0.002 | | | |

*Note.* Sphericity corrections not available for factors with 2 levels.
*Note.* Type III Sum of Squares
[a] Mauchly's test of sphericity indicates that the assumption of sphericity is violated (p < .05).

*Table S18. Between Subjects Effects*

| Cases | Sum of Squares | df | Mean Square | F | p | $\eta^2_p$ |
|---|---|---|---|---|---|---|
| Group | 0.002 | 1 | 0.002 | 0.040 | .841 | $5.520\times10^{-4}$ |
| Residuals | 3.488 | 73 | 0.048 | | | |

*Note.* Type III Sum of Squares

##### Post Hoc Tests

*Table S19. Post Hoc Comparisons - Task ✻ Relation - Conditional on Task*

| Task | | | Mean Difference | SE | df | t | Cohen's d | $p_{bonf}$ | $p_{holm}$ |
|---|---|---|---|---|---|---|---|---|---|
| TaxTask | Tax | Them | $9.551\times10^{-4}$ | 0.005 | 73 | 0.176 | 0.009 | 1.000 | 1.000 |
| | | Unrel | 0.006 | 0.007 | 73 | 0.821 | 0.053 | 1.000 | 1.000 |
| | Them | Unrel | 0.005 | 0.007 | 73 | 0.683 | 0.045 | 1.000 | 1.000 |
| ThemTask | Tax | Them | -0.049 | 0.007 | 73 | -6.787 | -0.456 | < .001*** | < .001*** |
| | | Unrel | -0.014 | 0.006 | 73 | -2.199 | -0.131 | .093 | .031* |
| | Them | Unrel | 0.035 | 0.006 | 73 | 5.592 | 0.325 | < .001*** | < .001*** |

* p < .05, *** p < .001
*Note.* P-value adjusted for comparing a family of 3 estimates.
*Note.* Results are averaged over the levels of: Group

*Table S20. Post Hoc Comparisons - Task ✻ Relation - Conditional on Relation*

| Relation | | | Mean Difference | SE | df | t | Cohen's d | $p_{bonf}$ | $p_{holm}$ |
|---|---|---|---|---|---|---|---|---|---|
| Tax | TaxTask | ThemTask | 0.014 | 0.013 | 73 | 1.095 | 0.129 | .277 | .277 |
| Them | | ThemTask | -0.036 | 0.013 | 73 | -2.754 | -0.336 | .007** | .007** |
| Unrel | | ThemTask | -0.006 | 0.013 | 73 | -0.456 | -0.055 | .650 | .650 |

** p < .01

*Note.* Results are averaged over the levels of: Group

*Table S21. Post Hoc Comparisons - Group ✻ Relation - Conditional on Group*

| Group | | | Mean Difference | SE | df | t | Cohen's d | $p_{bonf}$ | $p_{holm}$ |
|---|---|---|---|---|---|---|---|---|---|
| 1 | Tax | Them | -0.022 | 0.005 | 73 | -4.096 | -0.206 | < .001*** | < .001*** |
| | | Unrel | -0.021 | 0.007 | 73 | -2.955 | -0.193 | .013* | .008** |
| | Them | Unrel | 0.001 | 0.006 | 73 | 0.225 | 0.013 | 1.000 | .823 |
| 2 | Tax | Them | -0.026 | 0.006 | 73 | -4.381 | -0.242 | < .001*** | < .001*** |
| | | Unrel | 0.012 | 0.008 | 73 | 1.613 | 0.116 | .333 | .111 |
| | Them | Unrel | 0.038 | 0.007 | 73 | 5.680 | 0.357 | < .001*** | < .001*** |

* p < .05, ** p < .01, *** p < .001

*Note.* P-value adjusted for comparing a family of 3 estimates.

*Note.* Results are averaged over the levels of: Task

*Table S22. Post Hoc Comparisons - Group ✻ Relation - Conditional on Relation*

| Relation | | | Mean Difference | SE | df | t | Cohen's d | $p_{bonf}$ | $p_{holm}$ |
|---|---|---|---|---|---|---|---|---|---|
| Tax | Group1 | Group2 | -0.014 | 0.022 | 73 | -0.648 | -0.129 | .519 | .519 |
| Them | | Group2 | -0.018 | 0.022 | 73 | -0.815 | -0.165 | .418 | .418 |
| Unrel | | Group2 | 0.019 | 0.021 | 73 | 0.929 | 0.179 | .356 | .356 |

*Note.* Results are averaged over the levels of: Task

#### ROI 4: Right ATL

##### Repeated Measures ANOVA

*Table S23. Within Subjects Effects*

| Cases | Sphericity Correction | Sum of Squares | df | Mean Square | F | p | $\eta^2_p$ |
|---|---|---|---|---|---|---|---|
| Task | None | 0.004 | 1.000 | 0.004 | 0.262 | .610 | 0.004 |
| Task ✻ Group | None | 0.076 | 1.000 | 0.076 | 5.085 | .027 | 0.065 |
| Residuals | None | 1.096 | 73.000 | 0.015 | | | |
| Relation | None | 0.022[a] | 2.000[a] | 0.011[a] | 4.652[a] | .011[a] | 0.060 |
| | Huynh-Feldt | 0.022 | 1.681 | 0.013 | 4.652 | .016 | 0.060 |
| Relation ✻ Group | None | 0.013[a] | 2.000[a] | 0.007[a] | 2.805[a] | .064[a] | 0.037 |
| | Huynh-Feldt | 0.013 | 1.681 | 0.008 | 2.805 | .074 | 0.037 |
| Residuals | None | 0.346 | 146.000 | 0.002 | | | |
| | Huynh-Feldt | 0.346 | 122.695 | 0.003 | | | |
| Task ✻ Relation | None | 0.031[a] | 2.000[a] | 0.016[a] | 6.380[a] | .002[a] | 0.080 |
| | Huynh-Feldt | 0.031 | 1.894 | 0.017 | 6.380 | .003 | 0.080 |

*Table S23. Within Subjects Effects*

| Cases | Sphericity Correction | Sum of Squares | df | Mean Square | F | p | $\eta^2_p$ |
|---|---|---|---|---|---|---|---|
| Task ✻ Relation ✻ Group | None | 0.024[a] | 2.000[a] | 0.012[a] | 4.978[a] | .008[a] | 0.064 |
| | Huynh-Feldt | 0.024 | 1.894 | 0.013 | 4.978 | .009 | 0.064 |
| Residuals | None | 0.359 | 146.000 | 0.002 | | | |
| | Huynh-Feldt | 0.359 | 138.276 | 0.003 | | | |

*Note.* Sphericity corrections not available for factors with 2 levels.
*Note.* Type III Sum of Squares
[a] Mauchly's test of sphericity indicates that the assumption of sphericity is violated (p < .05).

*Table S24. Between Subjects Effects*

| Cases | Sum of Squares | df | Mean Square | F | p | $\eta^2_p$ |
|---|---|---|---|---|---|---|
| Group | 0.140 | 1 | 0.140 | 2.908 | .092 | 0.038 |
| Residuals | 3.516 | 73 | 0.048 | | | |

*Note.* Type III Sum of Squares

#### Post Hoc Tests

*Table S25. Post Hoc Comparisons - Group ✻ Task ✻ Relation - Conditional on Group*

| Group | | | Mean Difference | SE | df | t | Cohen's d | $p_{bonf}$ | $p_{holm}$ |
|---|---|---|---|---|---|---|---|---|---|
| 1 | TaxTask, Tax | ThemTask, Tax | 0.037 | 0.019 | 73 | 1.968 | 0.335 | .794 | .529 |
| | | TaxTask, Them | 0.004 | 0.010 | 73 | 0.404 | 0.036 | 1.000 | 1.000 |
| | | TaxTask, Unrel | -0.014 | 0.013 | 73 | -1.072 | -0.123 | 1.000 | 1.000 |
| | ThemTask, Tax | ThemTask, Them | -0.024 | 0.009 | 73 | -2.705 | -0.221 | .127 | .127 |
| | | ThemTask, Unrel | $3.251\times10^{-4}$ | 0.012 | 73 | 0.028 | 0.003 | 1.000 | 1.000 |
| | TaxTask, Them | ThemTask, Them | 0.009 | 0.016 | 73 | 0.546 | 0.078 | 1.000 | 1.000 |
| | | TaxTask, Unrel | -0.017 | 0.012 | 73 | -1.491 | -0.158 | 1.000 | .982 |
| | ThemTask, Them | ThemTask, Unrel | 0.025 | 0.010 | 73 | 2.507 | 0.224 | .216 | .173 |
| | TaxTask, Unrel | ThemTask, Unrel | 0.051 | 0.019 | 73 | 2.637 | 0.461 | .153 | .143 |
| 2 | TaxTask, Tax | ThemTask, Tax | 0.017 | 0.021 | 73 | 0.822 | 0.154 | 1.000 | 1.000 |
| | | TaxTask, Them | 0.011 | 0.011 | 73 | 1.064 | 0.103 | 1.000 | 1.000 |
| | | TaxTask, Unrel | 0.042 | 0.014 | 73 | 3.058 | 0.384 | .047* | .041* |
| | ThemTask, Tax | ThemTask, Them | -0.042 | 0.010 | 73 | -4.284 | -0.385 | < .001*** | < .001 |
| | | ThemTask, Unrel | -0.015 | 0.013 | 73 | -1.195 | -0.140 | 1.000 | 1.000 |
| | TaxTask, Them | ThemTask, Them | -0.037 | 0.017 | 73 | -2.124 | -0.334 | .556 | .371 |
| | | TaxTask, Unrel | 0.031 | 0.013 | 73 | 2.414 | 0.281 | .274 | .201 |
| | ThemTask, Them | ThemTask, Unrel | 0.027 | 0.011 | 73 | 2.494 | 0.245 | .223 | .179 |

*Table S25. Post Hoc Comparisons - Group ✻ Task ✻ Relation - Conditional on Group*

| Group | | | Mean Difference | SE | df | t | Cohen's d | $p_{bonf}$ | $p_{holm}$ |
|---|---|---|---|---|---|---|---|---|---|
| | TaxTask, Unrel | ThemTask, Unrel | -0.041 | 0.021 | 73 | -1.930 | -0.370 | .863 | .518 |

* p < .05, *** p < .001
*Note.* P-value adjusted for comparing a family of 15 estimates.

*Table S26. Post Hoc Comparisons - Group ✻ Task ✻ Relation - Conditional on Task*

| Task | | | Mean Difference | SE | df | t | Cohen's d | $p_{bonf}$ | $p_{holm}$ |
|---|---|---|---|---|---|---|---|---|---|
| TaxTask | GroupY, Tax | GroupO, Tax | -0.030 | 0.027 | 73 | -1.117 | -0.275 | 1.000 | 1.000 |
| | GroupY, Them | GroupO, Them | -0.023 | 0.027 | 73 | -0.843 | -0.208 | 1.000 | 1.000 |
| | GroupY, Unrel | GroupO, Unrel | 0.025 | 0.026 | 73 | 0.964 | 0.231 | 1.000 | 1.000 |
| ThemTask | GroupY, Tax | GroupO, Tax | -0.050 | 0.026 | 73 | -1.963 | -0.457 | .802 | .267 |
| | GroupY, Them | GroupO, Them | -0.068 | 0.023 | 73 | -2.970 | -0.621 | .060 | .048* |
| | GroupY, Unrel | GroupO, Unrel | -0.066 | 0.024 | 73 | -2.808 | -0.600 | .096 | .070 |

* p < .05, ** p < .01, *** p < .001
*Note.* P-value adjusted for comparing a family of 15 estimates.

#### ROI 5: Left vATL

##### Repeated Measures ANOVA

*Table S27. Within Subjects Effects*

| Cases | Sphericity Correction | Sum of Squares | df | Mean Square | F | p | $\eta^2_p$ |
|---|---|---|---|---|---|---|---|
| TASK | None | 0.053 | 1.000 | 0.053 | 3.461 | .067 | 0.045 |
| TASK ✻ Group | None | 0.052 | 1.000 | 0.052 | 3.385 | .070 | 0.044 |
| Residuals | None | 1.119 | 73.000 | 0.015 | | | |
| RELATION | None | 0.026 | 2.000 | 0.013 | 6.900 | .001 | 0.086 |
| | Huynh-Feldt | 0.026 | 1.969 | 0.013 | 6.900 | .001 | 0.086 |
| RELATION ✻ Group | None | $1.331 \times 10^{-4}$ | 2.000 | $6.655 \times 10^{-5}$ | 0.036 | .965 | $4.898 \times 10^{-4}$ |
| | Huynh-Feldt | $1.331 \times 10^{-4}$ | 1.969 | $6.761 \times 10^{-5}$ | 0.036 | .963 | $4.898 \times 10^{-4}$ |
| Residuals | None | 0.272 | 146.000 | 0.002 | | | |
| | Huynh-Feldt | 0.272 | 143.727 | 0.002 | | | |
| TASK ✻ RELATION | None | 0.002 | 2.000 | $9.724 \times 10^{-4}$ | 0.460 | .632 | 0.006 |
| | Huynh-Feldt | 0.002 | 1.939 | 0.001 | 0.460 | .627 | 0.006 |
| TASK ✻ RELATION ✻ Group | None | 0.005 | 2.000 | 0.003 | 1.203 | .303 | 0.016 |
| | Huynh-Feldt | 0.005 | 1.939 | 0.003 | 1.203 | .302 | 0.016 |
| Residuals | None | 0.309 | 146.000 | 0.002 | | | |
| | Huynh-Feldt | 0.309 | 141.582 | 0.002 | | | |

*Table S27. Within Subjects Effects*

| Cases | Sphericity Correction | Sum of Squares | df | Mean Square | F | p | η²p |
|---|---|---|---|---|---|---|---|

*Note.* Sphericity corrections not available for factors with 2 levels.
*Note.* Type III Sum of Squares

*Table S28. Between Subjects Effects*

| Cases | Sum of Squares | df | Mean Square | F | p | η²p |
|---|---|---|---|---|---|---|
| Group | 0.169 | 1 | 0.169 | 4.326 | .041 | 0.056 |
| Residuals | 2.859 | 73 | 0.039 | | | |

*Note.* Type III Sum of Squares

#### Post Hoc Tests

*Table S29. Post Hoc Comparisons - RELATION*

| | | Mean Difference | SE | df | t | Cohen's d | $p_{bonf}$ | $p_{holm}$ |
|---|---|---|---|---|---|---|---|---|
| Tax | Them | -0.019 | 0.004 | 73 | -4.147 | -0.182 | < .001 | < .001 |
| | Unrel | -0.010 | 0.005 | 73 | -1.963 | -0.103 | .160 | .107 |
| Them | Unrel | 0.008 | 0.005 | 73 | 1.562 | 0.079 | .368 | .123 |

*Note.* P-value adjusted for comparing a family of 3 estimates.
*Note.* Results are averaged over the levels of: Group, TASK

*Table S30. Post Hoc Comparisons - Group ✻ TASK - Conditional on Group*

| Group | | | Mean Difference | SE | df | t | Cohen's d | $p_{bonf}$ | $p_{holm}$ |
|---|---|---|---|---|---|---|---|---|---|
| 1 | TaxTask | ThemTask | $-2.389\times10^{-4}$ | 0.016 | 73 | -0.015 | -0.002 | .988 | .988 |
| 2 | | ThemTask | -0.043 | 0.017 | 73 | -2.502 | -0.425 | .015 | .015 |

*Note.* Results are averaged over the levels of: RELATION

*Table S31. Post Hoc Comparisons - Group ✻ TASK - Conditional on TASK*

| TASK | | | Mean Difference | SE | df | t | Cohen's d | $p_{bonf}$ | $p_{holm}$ |
|---|---|---|---|---|---|---|---|---|---|
| TaxTask | GroupY | GroupO | -0.017 | 0.022 | 73 | -0.787 | -0.171 | .434 | .434 |
| ThemTask | | GroupO | -0.061 | 0.022 | 73 | -2.742 | -0.593 | .008 | .008 |

*Note.* Results are averaged over the levels of: RELATION

### ROI 6: Right vATL

#### Repeated Measures ANOVA

*Table S32. Within Subjects Effects*

| Cases | Sphericity Correction | Sum of Squares | df | Mean Square | F | p | η²p |
|---|---|---|---|---|---|---|---|
| TASK | None | $2.357\times10^{-4}$ | 1.000 | $2.357\times10^{-4}$ | 0.010 | .920 | $1.400\times10^{-4}$ |
| TASK ✻ Group | None | 0.029 | 1.000 | 0.029 | 1.260 | .265 | 0.017 |

*Table S32. Within Subjects Effects*

| Cases | Sphericity Correction | Sum of Squares | df | Mean Square | F | p | $\eta^2_p$ |
|---|---|---|---|---|---|---|---|
| Residuals | None | 1.683 | 73.000 | 0.023 | | | |
| RELATION | None | 0.035 | 2.000 | 0.017 | 7.072 | .001 | 0.088 |
| | Huynh-Feldt | 0.035 | 1.929 | 0.018 | 7.072 | .001 | 0.088 |
| RELATION ✻ Group | None | $3.109\times10^{-5}$ | 2.000 | $1.554\times10^{-5}$ | 0.006 | .994 | $8.637\times10^{-5}$ |
| | Huynh-Feldt | $3.109\times10^{-5}$ | 1.929 | $1.612\times10^{-5}$ | 0.006 | .993 | $8.637\times10^{-5}$ |
| Residuals | None | 0.360 | 146.000 | 0.002 | | | |
| | Huynh-Feldt | 0.360 | 140.796 | 0.003 | | | |
| TASK ✻ RELATION | None | 0.014 | 2.000 | 0.007 | 1.918 | .151 | 0.026 |
| | Huynh-Feldt | 0.014 | 2.000 | 0.007 | 1.918 | .151 | 0.026 |
| TASK ✻ RELATION ✻ Group | None | 0.015 | 2.000 | 0.007 | 2.079 | .129 | 0.028 |
| | Huynh-Feldt | 0.015 | 2.000 | 0.007 | 2.079 | .129 | 0.028 |
| Residuals | None | 0.524 | 146.000 | 0.004 | | | |
| | Huynh-Feldt | 0.524 | 149.627 | 0.004 | | | |

*Note.* Sphericity corrections not available for factors with 2 levels.
*Note.* Type III Sum of Squares

*Table S33. Between Subjects Effects*

| Cases | Sum of Squares | df | Mean Square | F | p | $\eta^2_p$ |
|---|---|---|---|---|---|---|
| Group | 0.172 | 1 | 0.172 | 3.291 | .074 | 0.043 |
| Residuals | 3.820 | 73 | 0.052 | | | |

*Note.* Type III Sum of Squares

#### Post Hoc Tests

*Table S34. Post Hoc Comparisons - RELATION*

| | | Mean Difference | SE | df | t | Cohen's d | $p_{bonf}$ | $p_{holm}$ |
|---|---|---|---|---|---|---|---|---|
| Tax | Them | -0.021 | 0.005 | 73 | -4.028 | -0.173 | < .001 | < .001 |
| | Unrel | -0.005 | 0.006 | 73 | -0.969 | -0.045 | 1.000 | .336 |
| Them | Unrel | 0.015 | 0.006 | 73 | 2.408 | 0.128 | .056 | .037 |

*Note.* P-value adjusted for comparing a family of 3 estimates.
*Note.* Results are averaged over the levels of: Group, TASK

*Table S35. Post Hoc Comparisons - Group*

| | | Mean Difference | SE | df | t | Cohen's d | $p_{bonf}$ | $p_{holm}$ |
|---|---|---|---|---|---|---|---|---|
| GroupY | GroupO | -0.039 | 0.022 | 73 | -1.814 | -0.325 | .074 | .074 |

*Note.* Results are averaged over the levels of: RELATION, TASK

#### ROI 7: Left PreCS

##### Repeated Measures ANOVA

*Table S36. Within Subjects Effects*

| Cases | Sphericity Correction | Sum of Squares | df | Mean Square | F | p | $\eta^2_p$ |
|---|---|---|---|---|---|---|---|
| Task | None | 0.011 | 1.000 | 0.011 | 0.299 | .586 | 0.004 |
| Task ✻ Group | None | 0.192 | 1.000 | 0.192 | 5.228 | .025 | 0.067 |
| Residuals | None | 2.687 | 73.000 | 0.037 | | | |
| Relation | None | 0.781[a] | 2.000[a] | 0.390[a] | 60.551[a] | < .001[a] | 0.453 |
| | Huynh-Feldt | 0.781 | 1.687 | 0.463 | 60.551 | < .001 | 0.453 |
| Relation ✻ Group | None | 0.002[a] | 2.000[a] | 0.001[a] | 0.174[a] | .841[a] | 0.002 |
| | Huynh-Feldt | 0.002 | 1.687 | 0.001 | 0.174 | .804 | 0.002 |
| Residuals | None | 0.941 | 146.000 | 0.006 | | | |
| | Huynh-Feldt | 0.941 | 123.115 | 0.008 | | | |
| Task ✻ Relation | None | 0.028 | 2.000 | 0.014 | 2.875 | .060 | 0.038 |
| | Huynh-Feldt | 0.028 | 2.000 | 0.014 | 2.875 | .060 | 0.038 |
| Task ✻ Relation ✻ Group | None | 0.019 | 2.000 | 0.010 | 1.978 | .142 | 0.026 |
| | Huynh-Feldt | 0.019 | 2.000 | 0.010 | 1.978 | .142 | 0.026 |
| Residuals | None | 0.718 | 146.000 | 0.005 | | | |
| | Huynh-Feldt | 0.718 | 149.617 | 0.005 | | | |

*Note.* Sphericity corrections not available for factors with 2 levels.
*Note.* Type III Sum of Squares
[a] Mauchly's test of sphericity indicates that the assumption of sphericity is violated (p < .05).

*Table S37. Between Subjects Effects*

| Cases | Sum of Squares | df | Mean Square | F | p | $\eta^2_p$ |
|---|---|---|---|---|---|---|
| Group | 0.069 | 1 | 0.069 | 0.481 | .490 | 0.007 |
| Residuals | 10.431 | 73 | 0.143 | | | |

*Note.* Type III Sum of Squares

##### Post Hoc Tests

*Table S38. Post Hoc Comparisons - Group ✻ Task - Conditional on Group*

| Group | | | Mean Difference | SE | df | t | Cohen's d | $p_{bonf}$ | $p_{holm}$ |
|---|---|---|---|---|---|---|---|---|---|
| 1 | TaxTask | ThemTask | 0.032 | 0.024 | 73 | 1.292 | 0.172 | .200 | .200 |
| 2 | | ThemTask | -0.051 | 0.027 | 73 | -1.916 | -0.280 | .059 | .059 |

*Note.* Results are averaged over the levels of: Relation

*Table S39. Post Hoc Comparisons - Group ✻ Task - Conditional on Task*

| Task | | | Mean Difference | SE | df | t | Cohen's d | $p_{bonf}$ | $p_{holm}$ |
|---|---|---|---|---|---|---|---|---|---|
| TaxTask | GroupY | GroupO | 0.017 | 0.039 | 73 | 0.424 | 0.091 | .673 | .673 |
| ThemTask | | GroupO | -0.066 | 0.041 | 73 | -1.623 | -0.361 | .109 | .109 |

*Table S39. Post Hoc Comparisons - Group ✻ Task - Conditional on Task*

| Task | Mean Difference | SE | df | t | Cohen's d | $p_{bonf}$ | $p_{holm}$ |
|---|---|---|---|---|---|---|---|

*Note.* Results are averaged over the levels of: Relation

*Table S40. Post Hoc Comparisons - Relation*

| | | Mean Difference | SE | df | t | Cohen's d | $p_{bonf}$ | $p_{holm}$ |
|---|---|---|---|---|---|---|---|---|
| Tax | Them | -0.019 | 0.007 | 73 | -2.641 | -0.103 | .030* | .010* |
| | Unrel | 0.078 | 0.009 | 73 | 8.278 | 0.423 | < .001*** | < .001*** |
| Them | Unrel | 0.097 | 0.011 | 73 | 8.814 | 0.526 | < .001*** | < .001*** |

* p < .05, *** p < .001

*Note.* P-value adjusted for comparing a family of 3 estimates.

*Note.* Results are averaged over the levels of: Group, Task

#### ROI 8: Left IFG/MFG

##### Repeated Measures ANOVA

*Table S41. Within Subjects Effects*

| Cases | Sphericity Correction | Sum of Squares | df | Mean Square | F | p | $\eta^2_p$ |
|---|---|---|---|---|---|---|---|
| Task | None | $9.643 \times 10^{-4}$ | 1.000 | $9.643 \times 10^{-4}$ | 0.038 | .846 | $5.203 \times 10^{-4}$ |
| Task ✻ Group | None | 0.152 | 1.000 | 0.152 | 6.002 | .017 | 0.076 |
| Residuals | None | 1.852 | 73.000 | 0.025 | | | |
| Relation | None | 0.215[a] | 2.000[a] | 0.107[a] | 18.272[a] | < .001[a] | 0.200 |
| | Huynh-Feldt | 0.215 | 1.737 | 0.124 | 18.272 | < .001 | 0.200 |
| Relation ✻ Group | None | 0.008[a] | 2.000[a] | 0.004[a] | 0.719[a] | .489[a] | 0.010 |
| | Huynh-Feldt | 0.008 | 1.737 | 0.005 | 0.719 | .471 | 0.010 |
| Residuals | None | 0.859 | 146.000 | 0.006 | | | |
| | Huynh-Feldt | 0.859 | 126.822 | 0.007 | | | |
| Task ✻ Relation | None | 0.075 | 2.000 | 0.037 | 7.921 | < .001 | 0.098 |
| | Huynh-Feldt | 0.075 | 1.936 | 0.039 | 7.921 | < .001 | 0.098 |
| Task ✻ Relation ✻ Group | None | 0.017 | 2.000 | 0.009 | 1.807 | .168 | 0.024 |
| | Huynh-Feldt | 0.017 | 1.936 | 0.009 | 1.807 | .169 | 0.024 |
| Residuals | None | 0.688 | 146.000 | 0.005 | | | |
| | Huynh-Feldt | 0.688 | 141.318 | 0.005 | | | |

*Note.* Sphericity corrections not available for factors with 2 levels.

*Note.* Type III Sum of Squares

[a] Mauchly's test of sphericity indicates that the assumption of sphericity is violated (p < .05).

*Table S42. Between Subjects Effects*

| Cases | Sum of Squares | df | Mean Square | F | p | $\eta^2_p$ |
|---|---|---|---|---|---|---|
| Group | 0.088 | 1 | 0.088 | 0.781 | .380 | 0.011 |
| Residuals | 8.189 | 73 | 0.112 | | | |

*Note.* Type III Sum of Squares

#### Post Hoc Tests

*Table S43. Post Hoc Comparisons - Group ✻ Task - Conditional on Group*

| Group | | | Mean Difference | SE | df | t | Cohen's d | $p_{bonf}$ | $p_{holm}$ |
|---|---|---|---|---|---|---|---|---|---|
| 1 | TaxTask | ThemTask | 0.034 | 0.020 | 73 | 1.675 | 0.209 | .098 | .098 |
| 2 | | ThemTask | -0.040 | 0.022 | 73 | -1.789 | -0.245 | .078 | .078 |

*Note.* Results are averaged over the levels of: Relation

*Table S44. Post Hoc Comparisons - Group ✻ Task - Conditional on Task*

| Task | | | Mean Difference | SE | df | t | Cohen's d | $p_{bonf}$ | $p_{holm}$ |
|---|---|---|---|---|---|---|---|---|---|
| TaxTask | GroupY | GroupO | 0.009 | 0.037 | 73 | 0.238 | 0.055 | .812 | .812 |
| ThemTask | | GroupO | -0.065 | 0.033 | 73 | -1.989 | -0.400 | .050 | .050 |

*Note.* Results are averaged over the levels of: Relation

*Table S45. Post Hoc Comparisons - Task ✻ Relation - Conditional on Task*

| Task | | | Mean Difference | SE | df | t | Cohen's d | $p_{bonf}$ | $p_{holm}$ |
|---|---|---|---|---|---|---|---|---|---|
| TaxTask | Tax | Them | 0.014 | 0.009 | 73 | 1.530 | 0.084 | .391 | .130 |
| | | Unrel | 0.058 | 0.013 | 73 | 4.421 | 0.359 | < .001*** | < .001*** |
| | Them | Unrel | 0.045 | 0.012 | 73 | 3.735 | 0.275 | .001** | < .001*** |
| ThemTask | Tax | Them | -0.047 | 0.013 | 73 | -3.585 | -0.292 | .002** | .001** |
| | | Unrel | 0.013 | 0.010 | 73 | 1.285 | 0.081 | .608 | .203 |
| | Them | Unrel | 0.061 | 0.013 | 73 | 4.549 | 0.373 | < .001*** | < .001*** |

** p < .01, *** p < .001

*Note.* P-value adjusted for comparing a family of 3 estimates.

*Note.* Results are averaged over the levels of: Group

*Table S46. Post Hoc Comparisons - Task ✻ Relation - Conditional on Relation*

| Relation | | | Mean Difference | SE | df | t | Cohen's d | $p_{bonf}$ | $p_{holm}$ |
|---|---|---|---|---|---|---|---|---|---|
| Tax | TaxTask | ThemTask | 0.032 | 0.019 | 73 | 1.710 | 0.200 | .091 | .091 |
| Them | | ThemTask | -0.029 | 0.018 | 73 | -1.603 | -0.176 | .113 | .113 |
| Unrel | | ThemTask | -0.013 | 0.016 | 73 | -0.791 | -0.078 | .432 | .432 |

*Note.* Results are averaged over the levels of: Group

### ROI 9: Left IPS

#### Repeated Measures ANOVA

*Table S47. Within Subjects Effects*

| Cases | Sphericity Correction | Sum of Squares | df | Mean Square | F | p | $\eta^2_p$ | $\omega^2$ |
|---|---|---|---|---|---|---|---|---|
| Task | None | 0.005 | 1.000 | 0.005 | 0.112 | .739 | 0.002 | 0.000 |
| Task ✻ Group | None | 0.201 | 1.000 | 0.201 | 4.908 | .030 | 0.063 | 0.008 |

*Table S47. Within Subjects Effects*

| Cases | Sphericity Correction | Sum of Squares | df | Mean Square | F | p | $\eta^2_p$ | $\omega^2$ |
|---|---|---|---|---|---|---|---|---|
| Residuals | None | 2.987 | 73.000 | 0.041 | | | | |
| Relation | None | 2.690[a] | 2.000[a] | 1.345[a] | 109.076[a] | < .001[a] | 0.599 | 0.130 |
| | Huynh-Feldt | 2.690 | 1.698 | 1.584 | 109.076 | < .001 | 0.599 | 0.130 |
| Relation ✻ Group | None | 0.003[a] | 2.000[a] | 0.001[a] | 0.106[a] | .899[a] | 0.001 | 0.000 |
| | Huynh-Feldt | 0.003 | 1.698 | 0.002 | 0.106 | .869 | 0.001 | 0.000 |
| Residuals | None | 1.800 | 146.000 | 0.012 | | | | |
| | Huynh-Feldt | 1.800 | 123.925 | 0.015 | | | | |
| Task ✻ Relation | None | 0.040 | 2.000 | 0.020 | 2.899 | .058 | 0.038 | 0.002 |
| | Huynh-Feldt | 0.040 | 1.906 | 0.021 | 2.899 | .061 | 0.038 | 0.002 |
| Task ✻ Relation ✻ Group | None | 0.005 | 2.000 | 0.003 | 0.378 | .686 | 0.005 | 0.000 |
| | Huynh-Feldt | 0.005 | 1.906 | 0.003 | 0.378 | .676 | 0.005 | 0.000 |
| Residuals | None | 0.998 | 146.000 | 0.007 | | | | |
| | Huynh-Feldt | 0.998 | 139.126 | 0.007 | | | | |

*Note.* Sphericity corrections not available for factors with 2 levels.
*Note.* Type III Sum of Squares
[a] Mauchly's test of sphericity indicates that the assumption of sphericity is violated ($p < .05$).

*Table S48. Between Subjects Effects*

| Cases | Sum of Squares | df | Mean Square | F | p | $\eta^2_p$ | $\omega^2$ |
|---|---|---|---|---|---|---|---|
| Group | 0.381 | 1 | 0.381 | 1.755 | .189 | 0.023 | 0.005 |
| Residuals | 15.856 | 73 | 0.217 | | | | |

*Note.* Type III Sum of Squares

#### Post Hoc Tests

*Table S49. Post Hoc Comparisons - Group ✻ Task - Conditional on Group*

| Group | | | Mean Difference | SE | df | t | Cohen's d | $p_{holm}$ |
|---|---|---|---|---|---|---|---|---|
| 1 | TaxTask | ThemTask | 0.049 | 0.026 | 73 | 1.893 | 0.220 | .062 |
| 2 | | ThemTask | -0.036 | 0.028 | 73 | -1.272 | -0.162 | .207 |

*Note.* Results are averaged over the levels of: Relation

*Table S50. Post Hoc Comparisons - Group ✻ Task - Conditional on Task*

| Task | | | Mean Difference | SE | df | t | Cohen's d | $p_{holm}$ |
|---|---|---|---|---|---|---|---|---|
| TaxTask | GroupY | GroupO | 0.101 | 0.049 | 73 | 2.078 | 0.454 | .041 |
| ThemTask | | GroupO | 0.016 | 0.048 | 73 | 0.336 | 0.072 | .738 |

*Note.* Results are averaged over the levels of: Relation

*Table S51. Post Hoc Comparisons - Relation*

| | | Mean Difference | SE | df | t | Cohen's d | $p_{holm}$ |
|---|---|---|---|---|---|---|---|
| Tax | Them | -0.049 | 0.010 | 73 | -4.835 | -0.219 | < .001 |
| | Unrel | 0.135 | 0.013 | 73 | 10.501 | 0.607 | < .001 |
| Them | Unrel | 0.184 | 0.015 | 73 | 12.082 | 0.826 | < .001 |

*Note.* P-value adjusted for comparing a family of 3 estimates.
*Note.* Results are averaged over the levels of: Group, Task

#### ROI 10: Left IFGorb

##### Repeated Measures ANOVA

*Table S52. Within Subjects Effects*

| Cases | Sphericity Correction | Sum of Squares | df | Mean Square | F | p | $\eta^2_p$ |
|---|---|---|---|---|---|---|---|
| Task | None | 0.022 | 1.000 | 0.022 | 0.510 | .478 | 0.007 |
| Task ✻ Group | None | 0.093 | 1.000 | 0.093 | 2.197 | .143 | 0.029 |
| Residuals | None | 3.106 | 73.000 | 0.043 | | | |
| Relation | None | 1.523[a] | 2.000[a] | 0.761[a] | 112.708[a] | < .001[a] | 0.607 |
| | Huynh-Feldt | 1.523 | 1.785 | 0.853 | 112.708 | < .001 | 0.607 |
| Relation ✻ Group | None | 0.028[a] | 2.000[a] | 0.014[a] | 2.053[a] | .132[a] | 0.027 |
| | Huynh-Feldt | 0.028 | 1.785 | 0.016 | 2.053 | .138 | 0.027 |
| Residuals | None | 0.986 | 146.000 | 0.007 | | | |
| | Huynh-Feldt | 0.986 | 130.285 | 0.008 | | | |
| Task ✻ Relation | None | 0.090[a] | 2.000[a] | 0.045[a] | 8.364[a] | < .001[a] | 0.103 |
| | Huynh-Feldt | 0.090 | 1.893 | 0.047 | 8.364 | < .001 | 0.103 |
| Task ✻ Relation ✻ Group | None | 0.014[a] | 2.000[a] | 0.007[a] | 1.301[a] | .276[a] | 0.018 |
| | Huynh-Feldt | 0.014 | 1.893 | 0.007 | 1.301 | .275 | 0.018 |
| Residuals | None | 0.783 | 146.000 | 0.005 | | | |
| | Huynh-Feldt | 0.783 | 138.196 | 0.006 | | | |

*Note.* Sphericity corrections not available for factors with 2 levels.
*Note.* Type III Sum of Squares
[a] Mauchly's test of sphericity indicates that the assumption of sphericity is violated (p < .05).

*Table S53. Between Subjects Effects*

| Cases | Sum of Squares | df | Mean Square | F | p | $\eta^2_p$ |
|---|---|---|---|---|---|---|
| Group | 0.315 | 1 | 0.315 | 2.919 | .092 | 0.038 |
| Residuals | 7.871 | 73 | 0.108 | | | |

*Note.* Type III Sum of Squares

##### Post Hoc Tests

*Table S54. Post Hoc Comparisons - Task ✻ Relation - Conditional on Task*

| Task | | | Mean Difference | SE | df | t | Cohen's d | $p_{bonf}$ | $p_{holm}$ |
|---|---|---|---|---|---|---|---|---|---|
| TaxTask | Tax | Them | 0.034 | 0.011 | 73 | 3.136 | 0.200 | .007** | .002** |

*Table S54. Post Hoc Comparisons - Task ✻ Relation - Conditional on Task*

| Task | | | Mean Difference | SE | df | t | Cohen's d | $p_{bonf}$ | $p_{holm}$ |
|---|---|---|---|---|---|---|---|---|---|
| | | Unrel | 0.146 | 0.015 | 73 | 9.516 | 0.854 | < .001*** | < .001*** |
| | Them | Unrel | 0.112 | 0.012 | 73 | 9.683 | 0.654 | < .001*** | < .001*** |
| ThemTask | Tax | Them | -0.035 | 0.013 | 73 | -2.648 | -0.202 | .030* | .010** |
| | | Unrel | 0.102 | 0.011 | 73 | 9.068 | 0.598 | < .001*** | < .001*** |
| | Them | Unrel | 0.137 | 0.014 | 73 | 9.751 | 0.800 | < .001*** | < .001*** |

* p < .05, ** p < .01, *** p < .001

*Note.* P-value adjusted for comparing a family of 3 estimates.

*Note.* Results are averaged over the levels of: Group

*Table S55. Post Hoc Comparisons - Task ✻ Relation - Conditional on Relation*

| Relation | | | Mean Difference | SE | df | t | Cohen's d | $p_{bonf}$ | $p_{holm}$ |
|---|---|---|---|---|---|---|---|---|---|
| Tax | TaxTask | ThemTask | 0.051 | 0.023 | 73 | 2.222 | 0.301 | .029* | .029* |
| Them | | ThemTask | -0.017 | 0.021 | 73 | -0.810 | -0.101 | .420 | .420 |
| Unrel | | ThemTask | 0.008 | 0.021 | 73 | 0.367 | 0.045 | .715 | .715 |

* p < .05

*Note.* Results are averaged over the levels of: Group

#### ROI 11: Left pMTG/ITG

##### Repeated Measures ANOVA

*Table S56. Within Subjects Effects*

| Cases | Sphericity Correction | Sum of Squares | df | Mean Square | F | p | $\eta^2_p$ |
|---|---|---|---|---|---|---|---|
| Task | None | 0.100 | 1.000 | 0.100 | 3.478 | .066 | 0.045 |
| Task ✻ Group | None | 0.141 | 1.000 | 0.141 | 4.920 | .030 | 0.063 |
| Residuals | None | 2.098 | 73.000 | 0.029 | | | |
| Relation | None | 0.769[a] | 2.000[a] | 0.384[a] | 79.313[a] | < .001[a] | 0.521 |
| | Huynh-Feldt | 0.769 | 1.734 | 0.443 | 79.313 | < .001 | 0.521 |
| Relation ✻ Group | None | 0.001[a] | 2.000[a] | $7.025\times10^{-4}$[a] | 0.145[a] | .865[a] | 0.002 |
| | Huynh-Feldt | 0.001 | 1.734 | $8.101\times10^{-4}$ | 0.145 | .836 | 0.002 |
| Residuals | None | 0.708 | 146.000 | 0.005 | | | |
| | Huynh-Feldt | 0.708 | 126.608 | 0.006 | | | |
| Task ✻ Relation | None | 0.175 | 2.000 | 0.087 | 24.324 | < .001 | 0.250 |
| | Huynh-Feldt | 0.175 | 1.926 | 0.091 | 24.324 | < .001 | 0.250 |
| Task ✻ Relation ✻ Group | None | $2.246\times10^{-4}$ | 2.000 | $1.123\times10^{-4}$ | 0.031 | .969 | $4.281\times10^{-4}$ |
| | Huynh-Feldt | $2.246\times10^{-4}$ | 1.926 | $1.166\times10^{-4}$ | 0.031 | .966 | $4.281\times10^{-4}$ |
| Residuals | None | 0.525 | 146.000 | 0.004 | | | |
| | Huynh-Feldt | 0.525 | 140.587 | 0.004 | | | |

*Note.* Sphericity corrections not available for factors with 2 levels.

*Note.* Type III Sum of Squares

[a] Mauchly's test of sphericity indicates that the assumption of sphericity is violated (p < .05).

*Table S57. Between Subjects Effects*

| Cases | Sum of Squares | df | Mean Square | F | p | $\eta^2_p$ |
|---|---|---|---|---|---|---|
| Group | 0.040 | 1 | 0.040 | 0.371 | .545 | 0.005 |
| Residuals | 7.845 | 73 | 0.107 | | | |

*Note.* Type III Sum of Squares

#### Post Hoc Tests

*Table S58. Post Hoc Comparisons - Group ✻ Task - Conditional on Group*

| Group | | | Mean Difference | SE | df | t | Cohen's d | $p_{bonf}$ | $p_{holm}$ |
|---|---|---|---|---|---|---|---|---|---|
| 1 | TaxTask | ThemTask | 0.006 | 0.022 | 73 | 0.262 | 0.036 | .794 | .794 |
| 2 | | ThemTask | -0.066 | 0.024 | 73 | -2.761 | -0.410 | .007 | .007 |

*Note.* Results are averaged over the levels of: Relation

*Table S59. Post Hoc Comparisons - Group ✻ Task - Conditional on Task*

| Task | | | Mean Difference | SE | df | t | Cohen's d | $p_{bonf}$ | $p_{holm}$ |
|---|---|---|---|---|---|---|---|---|---|
| TaxTask | GroupY | GroupO | 0.017 | 0.036 | 73 | 0.462 | 0.105 | .645 | .645 |
| ThemTask | | GroupO | -0.055 | 0.034 | 73 | -1.619 | -0.341 | .110 | .110 |

*Note.* Results are averaged over the levels of: Relation

*Table S60. Post Hoc Comparisons - Task ✻ Relation - Conditional on Task*

| Task | | | Mean Difference | SE | df | t | Cohen's d | $p_{bonf}$ | $p_{holm}$ |
|---|---|---|---|---|---|---|---|---|---|
| TaxTask | Tax | Them | 0.004 | 0.009 | 73 | 0.511 | 0.028 | 1.000 | .611 |
| | | Unrel | 0.080 | 0.012 | 73 | 6.570 | 0.500 | < .001 | < .001 |
| | Them | Unrel | 0.075 | 0.010 | 73 | 7.273 | 0.472 | < .001 | < .001 |
| ThemTask | Tax | Them | -0.092 | 0.011 | 73 | -8.483 | -0.579 | < .001 | < .001 |
| | | Unrel | 0.035 | 0.009 | 73 | 3.711 | 0.219 | .001 | < .001 |
| | Them | Unrel | 0.127 | 0.012 | 73 | 10.690 | 0.798 | < .001 | < .001 |

*Note.* P-value adjusted for comparing a family of 3 estimates.
*Note.* Results are averaged over the levels of: Group

*Table S61. Post Hoc Comparisons - Task ✻ Relation - Conditional on Relation*

| Relation | | | Mean Difference | SE | df | t | Cohen's d | $p_{bonf}$ | $p_{holm}$ |
|---|---|---|---|---|---|---|---|---|---|
| Tax | TaxTask | ThemTask | 0.017 | 0.019 | 73 | 0.930 | 0.108 | .355 | .355 |
| Them | | ThemTask | -0.080 | 0.018 | 73 | -4.360 | -0.498 | < .001 | < .001 |
| Unrel | | ThemTask | -0.028 | 0.017 | 73 | -1.628 | -0.173 | .108 | .108 |

*Note.* Results are averaged over the levels of: Group

#### ROI 12: Bilateral PreCun

##### Repeated Measures ANOVA

*Table S62. Within Subjects Effects*

| Cases | Sphericity Correction | Sum of Squares | df | Mean Square | F | p | $\eta^2_p$ |
|---|---|---|---|---|---|---|---|
| Task | None | 0.096 | 1.000 | 0.096 | 2.733 | .103 | 0.036 |
| Task ✻ Group | None | 0.155 | 1.000 | 0.155 | 4.393 | .040 | 0.057 |
| Residuals | None | 2.571 | 73.000 | 0.035 | | | |
| Relation | None | 0.802[a] | 2.000[a] | 0.401[a] | 72.933[a] | < .001[a] | 0.500 |
| | Huynh-Feldt | 0.802 | 1.773 | 0.452 | 72.933 | < .001 | 0.500 |
| Relation ✻ Group | None | 0.008[a] | 2.000[a] | 0.004[a] | 0.688[a] | .504[a] | 0.009 |
| | Huynh-Feldt | 0.008 | 1.773 | 0.004 | 0.688 | .487 | 0.009 |
| Residuals | None | 0.803 | 146.000 | 0.005 | | | |
| | Huynh-Feldt | 0.803 | 129.402 | 0.006 | | | |
| Task ✻ Relation | None | 0.015 | 2.000 | 0.008 | 1.992 | .140 | 0.027 |
| | Huynh-Feldt | 0.015 | 2.000 | 0.008 | 1.992 | .140 | 0.027 |
| Task ✻ Relation ✻ Group | None | 0.008 | 2.000 | 0.004 | 1.030 | .360 | 0.014 |
| | Huynh-Feldt | 0.008 | 2.000 | 0.004 | 1.030 | .360 | 0.014 |
| Residuals | None | 0.552 | 146.000 | 0.004 | | | |
| | Huynh-Feldt | 0.552 | 148.341 | 0.004 | | | |

*Note.* Sphericity corrections not available for factors with 2 levels.
*Note.* Type III Sum of Squares
[a] Mauchly's test of sphericity indicates that the assumption of sphericity is violated (p < .05).

*Table S63. Between Subjects Effects*

| Cases | Sum of Squares | df | Mean Square | F | p | $\eta^2_p$ |
|---|---|---|---|---|---|---|
| Group | 0.718 | 1 | 0.718 | 4.094 | .047 | 0.053 |
| Residuals | 12.805 | 73 | 0.175 | | | |

*Note.* Type III Sum of Squares

##### Post Hoc Tests

*Table S64. Post Hoc Comparisons - Group ✻ Task - Conditional on Group*

| Group | | | Mean Difference | SE | df | t | Cohen's d | $p_{bonf}$ | $p_{holm}$ |
|---|---|---|---|---|---|---|---|---|---|
| 1 | TaxTask | ThemTask | 0.067 | 0.024 | 73 | 2.784 | 0.341 | .007** | .007** |
| 2 | | ThemTask | -0.008 | 0.026 | 73 | -0.299 | -0.040 | .765 | .765 |

** p < .01
*Note.* Results are averaged over the levels of: Relation

*Table S65. Post Hoc Comparisons - Group ✻ Task - Conditional on Task*

| Task | | | Mean Difference | SE | df | t | Cohen's d | $p_{bonf}$ | $p_{holm}$ |
|---|---|---|---|---|---|---|---|---|---|
| TaxTask | GroupY | GroupO | 0.117 | 0.045 | 73 | 2.624 | 0.601 | .011* | .011* |
| ThemTask | | GroupO | 0.043 | 0.042 | 73 | 1.021 | 0.220 | .311 | .311 |

*Table S65. Post Hoc Comparisons - Group ✻ Task - Conditional on Task*

| Task | Mean Difference | SE | df | t | Cohen's d | $p_{bonf}$ | $p_{holm}$ |
|---|---|---|---|---|---|---|---|

* p < .05
*Note.* Results are averaged over the levels of: Relation

*Table S66. Post Hoc Comparisons - Relation*

| | | Mean Difference | SE | df | t | Cohen's d | $p_{bonf}$ | $p_{holm}$ |
|---|---|---|---|---|---|---|---|---|
| Tax | Them | -0.007 | 0.007 | 73 | -0.988 | -0.034 | .979 | .326 |
| | Unrel | 0.086 | 0.009 | 73 | 9.382 | 0.442 | < .001*** | < .001*** |
| Them | Unrel | 0.093 | 0.010 | 73 | 9.720 | 0.476 | < .001*** | < .001*** |

*** p < .001
*Note.* P-value adjusted for comparing a family of 3 estimates.
*Note.* Results are averaged over the levels of: Group, Task

#### ROI 13: Bilateral mPFC

##### Repeated Measures ANOVA

*Table S67. Within Subjects Effects*

| Cases | Sphericity Correction | Sum of Squares | df | Mean Square | F | p | $\eta^2_p$ |
|---|---|---|---|---|---|---|---|
| Task | None | 0.008 | 1.000 | 0.008 | 0.223 | .638 | 0.003 |
| Task ✻ Group | None | 0.101 | 1.000 | 0.101 | 2.992 | .088 | 0.039 |
| Residuals | None | 2.475 | 73.000 | 0.034 | | | |
| Relation | None | 0.405[a] | 2.000[a] | 0.203[a] | 38.596[a] | < .001[a] | 0.346 |
| | Huynh-Feldt | 0.405 | 1.780 | 0.228 | 38.596 | < .001 | 0.346 |
| Relation ✻ Group | None | $2.213\times10^{-4}$[a] | 2.000[a] | $1.106\times10^{-4}$[a] | 0.021[a] | .979[a] | $2.887\times10^{-4}$ |
| | Huynh-Feldt | $2.213\times10^{-4}$ | 1.780 | $1.243\times10^{-4}$ | 0.021 | .970 | $2.887\times10^{-4}$ |
| Residuals | None | 0.766 | 146.000 | 0.005 | | | |
| | Huynh-Feldt | 0.766 | 129.931 | 0.006 | | | |
| Task ✻ Relation | None | 0.144 | 2.000 | 0.072 | 16.755 | < .001 | 0.187 |
| | Huynh-Feldt | 0.144 | 1.998 | 0.072 | 16.755 | < .001 | 0.187 |
| Task ✻ Relation ✻ Group | None | 0.016 | 2.000 | 0.008 | 1.912 | .151 | 0.026 |
| | Huynh-Feldt | 0.016 | 1.998 | 0.008 | 1.912 | .152 | 0.026 |
| Residuals | None | 0.628 | 146.000 | 0.004 | | | |
| | Huynh-Feldt | 0.628 | 145.864 | 0.004 | | | |

*Note.* Sphericity corrections not available for factors with 2 levels.
*Note.* Type III Sum of Squares
[a] Mauchly's test of sphericity indicates that the assumption of sphericity is violated (p < .05).

*Table S68. Between Subjects Effects*

| Cases | Sum of Squares | df | Mean Square | F | p | $\eta^2_p$ |
|---|---|---|---|---|---|---|
| Group | 0.002 | 1 | 0.002 | 0.018 | .893 | $2.478\times10^{-4}$ |
| Residuals | 7.135 | 73 | 0.098 | | | |

*Table S68. Between Subjects Effects*

| Cases | Sum of Squares | df | Mean Square | F | p | $\eta^2_p$ |
|---|---|---|---|---|---|---|

*Note.* Type III Sum of Squares

#### Post Hoc Tests

*Table S69. Post Hoc Comparisons - Task ✻ Relation - Conditional on Task*

| Task | | | Mean Difference | SE | df | t | Cohen's d | $p_{bonf}$ | $p_{holm}$ |
|---|---|---|---|---|---|---|---|---|---|
| TaxTask | Tax | Them | 0.027 | 0.010 | 73 | 2.786 | 0.170 | .020* | .007** |
| | | Unrel | 0.077 | 0.014 | 73 | 5.684 | 0.487 | < .001*** | < .001*** |
| | Them | Unrel | 0.050 | 0.012 | 73 | 4.170 | 0.317 | < .001*** | < .001*** |
| ThemTask | Tax | Them | -0.061 | 0.010 | 73 | -5.833 | -0.385 | < .001*** | < .001*** |
| | | Unrel | 0.030 | 0.011 | 73 | 2.868 | 0.191 | .016* | .005** |
| | Them | Unrel | 0.091 | 0.011 | 73 | 8.086 | 0.576 | < .001*** | < .001*** |

* p < .05, ** p < .01, *** p < .001

*Note.* P-value adjusted for comparing a family of 3 estimates.

*Note.* Results are averaged over the levels of: Group

*Table S70. Post Hoc Comparisons - Task ✻ Relation - Conditional on Relation*

| Relation | | | Mean Difference | SE | df | t | Cohen's d | $p_{bonf}$ | $p_{holm}$ |
|---|---|---|---|---|---|---|---|---|---|
| Tax | TaxTask | ThemTask | 0.037 | 0.020 | 73 | 1.858 | 0.232 | .067 | .067 |
| Them | | ThemTask | -0.051 | 0.021 | 73 | -2.492 | -0.323 | .015* | .015* |
| Unrel | | ThemTask | -0.010 | 0.018 | 73 | -0.560 | -0.064 | .577 | .577 |

* p < .05

*Note.* Results are averaged over the levels of: Group

### ROI 14: Left IFG/MFG (defined using all conditions)

#### Repeated Measures ANOVA

*Table S71. Within Subjects Effects*

| Cases | Sphericity Correction | Sum of Squares | df | Mean Square | F | p | $\eta^2_p$ |
|---|---|---|---|---|---|---|---|
| TASK | None | 0.002 | 1.000 | 0.002 | 0.084 | .773 | 0.001 |
| TASK ✻ Group | None | 0.151 | 1.000 | 0.151 | 5.711 | .019 | 0.073 |
| Residuals | None | 1.926 | 73.000 | 0.026 | | | |
| RELATION | None | 0.133[a] | 2.000[a] | 0.066[a] | 10.420[a] | < .001[a] | 0.125 |
| | Huynh-Feldt | 0.133 | 1.674 | 0.079 | 10.420 | < .001 | 0.125 |
| RELATION ✻ Group | None | 0.008[a] | 2.000[a] | 0.004[a] | 0.590[a] | .556[a] | 0.008 |
| | Huynh-Feldt | 0.008 | 1.674 | 0.004 | 0.590 | .527 | 0.008 |
| Residuals | None | 0.930 | 146.000 | 0.006 | | | |
| | Huynh-Feldt | 0.930 | 122.205 | 0.008 | | | |
| TASK ✻ RELATION | None | 0.029 | 2.000 | 0.015 | 2.923 | .057 | 0.038 |
| | Huynh-Feldt | 0.029 | 1.922 | 0.015 | 2.923 | .059 | 0.038 |
| TASK ✻ RELATION ✻ Group | None | 0.025 | 2.000 | 0.013 | 2.559 | .081 | 0.034 |
| | Huynh-Feldt | 0.025 | 1.922 | 0.013 | 2.559 | .083 | 0.034 |

*Table S71. Within Subjects Effects*

| Cases | Sphericity Correction | Sum of Squares | df | Mean Square | F | p | $\eta^2_p$ |
|---|---|---|---|---|---|---|---|
| Residuals | None | 0.725 | 146.000 | 0.005 | | | |
| | Huynh-Feldt | 0.725 | 140.283 | 0.005 | | | |

*Note.* Sphericity corrections not available for factors with 2 levels.
*Note.* Type III Sum of Squares
[a] Mauchly's test of sphericity indicates that the assumption of sphericity is violated (p < .05).

*Table S72. Between Subjects Effects*

| Cases | Sum of Squares | df | Mean Square | F | p | $\eta^2_p$ |
|---|---|---|---|---|---|---|
| Group | 0.092 | 1 | 0.092 | 0.770 | .383 | 0.010 |
| Residuals | 8.690 | 73 | 0.119 | | | |

*Note.* Type III Sum of Squares

#### Post Hoc Tests

*Table S73. Post Hoc Comparisons - Group ✻ TASK - Conditional on Group*

| Group | | | Mean Difference | SE | df | t | Cohen's d | $p_{bonf}$ | $p_{holm}$ |
|---|---|---|---|---|---|---|---|---|---|
| 1 | TaxTask | ThemTask | 0.032 | 0.021 | 73 | 1.560 | 0.193 | .123 | .123 |
| 2 | | ThemTask | -0.041 | 0.023 | 73 | -1.812 | -0.246 | .074 | .074 |

*Note.* Results are averaged over the levels of: RELATION

*Table S74. Post Hoc Comparisons - Group ✻ TASK - Conditional on TASK*

| TASK | | | Mean Difference | SE | df | t | Cohen's d | $p_{bonf}$ | $p_{holm}$ |
|---|---|---|---|---|---|---|---|---|---|
| TaxTask | Group1 | Group2 | 0.008 | 0.037 | 73 | 0.217 | 0.048 | .829 | .829 |
| ThemTask | | Group2 | -0.065 | 0.035 | 73 | -1.871 | -0.391 | .065 | .065 |

*Note.* Results are averaged over the levels of: RELATION

*Table S75. Post Hoc Comparisons - TASK ✻ RELATION - Conditional on TASK*

| TASK | | | Mean Difference | SE | df | t | Cohen's d | $p_{bonf}$ | $p_{holm}$ |
|---|---|---|---|---|---|---|---|---|---|
| TaxTask | Tax | Them | 0.001 | 0.009 | 73 | 0.135 | 0.008 | 1.000 | .893 |
| | | Unrel | 0.040 | 0.013 | 73 | 3.049 | 0.240 | .010 | .006 |
| | Them | Unrel | 0.039 | 0.012 | 73 | 3.224 | 0.233 | .006 | .006 |
| ThemTask | Tax | Them | -0.036 | 0.013 | 73 | -2.701 | -0.213 | .026 | .017 |
| | | Unrel | 0.009 | 0.011 | 73 | 0.865 | 0.057 | 1.000 | .390 |
| | Them | Unrel | 0.045 | 0.015 | 73 | 3.107 | 0.269 | .008 | .008 |

*Note.* P-value adjusted for comparing a family of 3 estimates.
*Note.* Results are averaged over the levels of: Group

*Table S76. Post Hoc Comparisons - TASK ✻ RELATION - Conditional on RELATION*

| RELATION | | | Mean Difference | SE | df | t | Cohen's d | $p_{bonf}$ | $p_{holm}$ |
|---|---|---|---|---|---|---|---|---|---|
| Tax | TaxTask | ThemTask | 0.018 | 0.019 | 73 | 0.945 | 0.108 | .348 | .348 |
| Them | | ThemTask | -0.019 | 0.019 | 73 | -1.011 | -0.112 | .316 | .316 |
| Unrel | | ThemTask | -0.013 | 0.016 | 73 | -0.776 | -0.075 | .440 | .440 |

*Note.* Results are averaged over the levels of: Group

#### ROI 15: Left pMTG/ITG (defined using all conditions)

##### Repeated Measures ANOVA

*Table S77. Within Subjects Effects*

| Cases | Sphericity Correction | Sum of Squares | df | Mean Square | F | p | $\eta^2_p$ |
|---|---|---|---|---|---|---|---|
| TASK | None | 0.055 | 1.000 | 0.055 | 2.265 | .137 | 0.030 |
| TASK ✻ Group | None | 0.109 | 1.000 | 0.109 | 4.501 | .037 | 0.058 |
| Residuals | None | 1.772 | 73.000 | 0.024 | | | |
| RELATION | None | 0.583[a] | 2.000[a] | 0.292[a] | 54.536[a] | < .001[a] | 0.428 |
| | Huynh-Feldt | 0.583 | 1.704 | 0.342 | 54.536 | < .001 | 0.428 |
| RELATION ✻ Group | None | 0.002[a] | 2.000[a] | 0.001[a] | 0.223[a] | .800[a] | 0.003 |
| | Huynh-Feldt | 0.002 | 1.704 | 0.001 | 0.223 | .765 | 0.003 |
| Residuals | None | 0.781 | 146.000 | 0.005 | | | |
| | Huynh-Feldt | 0.781 | 124.420 | 0.006 | | | |
| TASK ✻ RELATION | None | 0.063 | 2.000 | 0.032 | 10.300 | < .001 | 0.124 |
| | Huynh-Feldt | 0.063 | 1.906 | 0.033 | 10.300 | < .001 | 0.124 |
| TASK ✻ RELATION ✻ Group | None | 0.002 | 2.000 | 0.001 | 0.382 | .683 | 0.005 |
| | Huynh-Feldt | 0.002 | 1.906 | 0.001 | 0.382 | .673 | 0.005 |
| Residuals | None | 0.450 | 146.000 | 0.003 | | | |
| | Huynh-Feldt | 0.450 | 139.149 | 0.003 | | | |

*Note.* Sphericity corrections not available for factors with 2 levels.
*Note.* Type III Sum of Squares
[a] Mauchly's test of sphericity indicates that the assumption of sphericity is violated (p < .05).

*Table S78. Between Subjects Effects*

| Cases | Sum of Squares | df | Mean Square | F | p | $\eta^2_p$ |
|---|---|---|---|---|---|---|
| Group | 0.077 | 1 | 0.077 | 0.816 | .369 | 0.011 |
| Residuals | 6.880 | 73 | 0.094 | | | |

*Note.* Type III Sum of Squares

#### Post Hoc Tests

*Table S79. Post Hoc Comparisons - Group ✻ TASK - Conditional on Group*

| Group | | | Mean Difference | SE | df | t | Cohen's d | $p_{bonf}$ | $p_{holm}$ |
|---|---|---|---|---|---|---|---|---|---|
| 1 | TaxTask | ThemTask | 0.009 | 0.020 | 73 | 0.458 | 0.061 | .648 | .648 |
| 2 | | ThemTask | -0.054 | 0.022 | 73 | -2.453 | -0.356 | .017 | .017 |

*Note.* Results are averaged over the levels of: RELATION

*Table S80. Post Hoc Comparisons - Group ✻ TASK - Conditional on TASK*

| TASK | | | Mean Difference | SE | df | t | Cohen's d | $p_{bonf}$ | $p_{holm}$ |
|---|---|---|---|---|---|---|---|---|---|
| TaxTask | Group1 | Group2 | 0.005 | 0.034 | 73 | 0.146 | 0.034 | .884 | .884 |
| ThemTask | | Group2 | -0.058 | 0.031 | 73 | -1.876 | -0.383 | .065 | .065 |

*Note.* Results are averaged over the levels of: RELATION

*Table S81. Post Hoc Comparisons - TASK ✻ RELATION - Conditional on TASK*

| TASK | | | Mean Difference | SE | df | t | Cohen's d | $p_{bonf}$ | $p_{holm}$ |
|---|---|---|---|---|---|---|---|---|---|
| TaxTask | Tax | Them | -0.007 | 0.009 | 73 | -0.814 | -0.049 | 1.000 | .418 |
| | | Unrel | 0.066 | 0.012 | 73 | 5.294 | 0.438 | < .001 | < .001 |
| | Them | Unrel | 0.073 | 0.010 | 73 | 7.032 | 0.487 | < .001 | < .001 |
| ThemTask | Tax | Them | -0.066 | 0.010 | 73 | -6.469 | -0.438 | < .001 | < .001 |
| | | Unrel | 0.037 | 0.009 | 73 | 3.968 | 0.249 | < .001 | < .001 |
| | Them | Unrel | 0.103 | 0.012 | 73 | 8.592 | 0.687 | < .001 | < .001 |

*Note.* P-value adjusted for comparing a family of 3 estimates.
*Note.* Results are averaged over the levels of: Group

*Table S82. Post Hoc Comparisons - TASK ✻ RELATION - Conditional on RELATION*

| RELATION | | | Mean Difference | SE | df | t | Cohen's d | $p_{bonf}$ | $p_{holm}$ |
|---|---|---|---|---|---|---|---|---|---|
| Tax | TaxTask | ThemTask | 0.007 | 0.018 | 73 | 0.373 | 0.045 | .710 | .710 |
| Them | | ThemTask | -0.052 | 0.016 | 73 | -3.212 | -0.344 | .002 | .002 |
| Unrel | | ThemTask | -0.022 | 0.015 | 73 | -1.422 | -0.144 | .159 | .159 |

*Note.* Results are averaged over the levels of: Group

**Table S83. Stimulus characteristics.** Ratings for taxonomic and thematic associations were obtained from the same participants in the fMRI study after they completed the main experiment (Likert scale from 1-5).

| Pair | Relationship | Probe | Target | Mean Taxonomic Rating | Mean Thematic Rating | SD Taxonomic Rating | SD Thematic Rating | Probe Manipulable | Probe Natural | Target Manipulable | Target Natural |
|---|---|---|---|---|---|---|---|---|---|---|---|
| 1 | Taxonomic | apple | pineapple | 4.34 | 2.24 | 1.19 | 1.41 | 1 | 1 | 1 | 1 |
| 2 | Taxonomic | daisy | tree | 3.73 | 2.04 | 1.31 | 1.22 | 1 | 1 | 1 | 1 |
| 3 | Taxonomic | sneaker | hat | 3.78 | 2.53 | 1.24 | 1.26 | 1 | 0 | 1 | 0 |
| 4 | Taxonomic | rose | plant | 3.68 | 2.62 | 1.42 | 1.38 | 1 | 1 | 1 | 1 |
| 5 | Taxonomic | smallboat | smalltruck | 3.24 | 1.72 | 1.31 | 1.17 | 0 | 0 | 0 | 0 |
| 6 | Taxonomic | sink | fridge | 2.42 | 1.71 | 1.41 | 0.90 | 0 | 0 | 0 | 0 |
| 7 | Taxonomic | bee | spider | 4.34 | 2.44 | 1.23 | 1.50 | 0 | 1 | 0 | 1 |
| 8 | Taxonomic | hoe | pitchfork | 4.25 | 3.06 | 1.33 | 1.47 | 1 | 0 | 1 | 0 |
| 9 | Taxonomic | truck | largeboat | 3.59 | 2.42 | 1.38 | 1.35 | 0 | 0 | 0 | 0 |
| 10 | Taxonomic | caterpillar | snail | 4.22 | 2.19 | 1.17 | 1.27 | 0 | 1 | 0 | 1 |
| 11 | Taxonomic | ivy | wheat | 4.31 | 2.46 | 1.12 | 1.48 | 1 | 1 | 1 | 1 |
| 12 | Taxonomic | dog | bear | 4.25 | 1.94 | 1.23 | 1.34 | 0 | 1 | 0 | 1 |
| 13 | Taxonomic | ear | foot | 4.17 | 2.17 | 1.20 | 1.38 | 1 | 1 | 1 | 1 |
| 14 | Taxonomic | motorbike | helicopter | 3.72 | 2.06 | 1.29 | 1.28 | 0 | 0 | 0 | 0 |
| 15 | Taxonomic | mouse | turtle | 4.14 | 1.67 | 1.17 | 1.14 | 0 | 1 | 0 | 1 |
| 16 | Taxonomic | Xmastree | flower | 3.17 | 1.74 | 1.39 | 1.19 | 1 | 1 | 1 | 1 |
| 17 | Taxonomic | shirt | sock | 4.00 | 2.42 | 1.26 | 1.32 | 1 | 0 | 1 | 0 |
| 18 | Taxonomic | bucket | box | 3.10 | 1.71 | 1.51 | 1.04 | 1 | 0 | 1 | 0 |
| 19 | Taxonomic | tropicalfish | elephant | 3.77 | 1.65 | 1.25 | 1.18 | 0 | 1 | 0 | 1 |
| 20 | Taxonomic | camel | rooster | 3.69 | 1.51 | 1.30 | 1.07 | 0 | 1 | 0 | 1 |
| 21 | Taxonomic | scooter | rollerskates | 3.24 | 2.00 | 1.29 | 1.26 | 0 | 0 | 0 | 0 |
| 22 | Taxonomic | kite | balloon | 3.61 | 2.57 | 1.51 | 1.46 | 1 | 0 | 1 | 0 |
| 23 | Taxonomic | strawberry | grapes | 4.46 | 2.65 | 1.20 | 1.46 | 1 | 1 | 1 | 1 |
| 24 | Taxonomic | mitten | pajamas | 3.61 | 2.76 | 1.42 | 1.37 | 1 | 0 | 1 | 0 |
| 25 | Taxonomic | car | bus | 4.44 | 2.75 | 1.02 | 1.44 | 0 | 0 | 0 | 0 |
| 26 | Taxonomic | hand | eye | 3.97 | 2.46 | 1.33 | 1.24 | 1 | 1 | 1 | 1 |
| 27 | Taxonomic | monkey | swan | 4.00 | 1.53 | 1.16 | 1.05 | 0 | 1 | 0 | 1 |
| 28 | Taxonomic | cow | snake | 3.93 | 1.56 | 1.24 | 0.98 | 0 | 1 | 0 | 1 |
| 29 | Taxonomic | sparrow | dolphin | 3.86 | 1.50 | 1.26 | 1.06 | 0 | 1 | 0 | 1 |
| 30 | Taxonomic | nose | mouth | 4.33 | 2.86 | 1.22 | 1.39 | 1 | 1 | 1 | 1 |
| 31 | Taxonomic | sled | unicycle | 3.04 | 1.62 | 1.36 | 0.98 | 1 | 0 | 1 | 0 |
| 32 | Taxonomic | sheep | panda | 4.15 | 1.79 | 1.21 | 1.22 | 0 | 1 | 0 | 1 |
| 33 | Taxonomic | tent | house | 3.55 | 1.97 | 1.39 | 1.32 | 0 | 0 | 0 | 0 |
| 34 | Taxonomic | cherries | pear | 4.35 | 2.62 | 1.30 | 1.51 | 1 | 1 | 1 | 1 |
| 35 | Taxonomic | squirrel | goat | 4.13 | 1.71 | 1.25 | 1.16 | 0 | 1 | 0 | 1 |
| 36 | Taxonomic | hammer | pliers | 4.30 | 2.99 | 1.43 | 1.61 | 1 | 0 | 1 | 0 |
| 37 | Taxonomic | pipe | cigarette | 3.57 | 2.97 | 1.60 | 1.53 | 1 | 0 | 1 | 0 |

| | | | | | | | | | | | |
|---|---|---|---|---|---|---|---|---|---|---|---|
| 38 | Taxonomic | carrot | tomato | 4.35 | 2.57 | 1.10 | 1.55 | 1 | 1 | 1 | 1 |
| 39 | Taxonomic | axe | drill | 4.16 | 2.38 | 1.14 | 1.40 | 1 | 0 | 1 | 0 |
| 40 | Taxonomic | bed | chair | 4.06 | 2.57 | 1.17 | 1.44 | 0 | 0 | 0 | 0 |
| 41 | Taxonomic | couch | dresser | 3.99 | 2.61 | 1.37 | 1.49 | 0 | 0 | 0 | 0 |
| 42 | Taxonomic | razor | tweezers | 3.48 | 2.65 | 1.52 | 1.39 | 1 | 0 | 1 | 0 |
| 43 | Taxonomic | telescope | microscope | 4.22 | 2.38 | 1.22 | 1.57 | 1 | 0 | 1 | 0 |
| 44 | Taxonomic | train | bike | 3.73 | 2.35 | 1.23 | 1.30 | 0 | 0 | 0 | 0 |
| 45 | Taxonomic | castle | hut | 3.69 | 1.93 | 1.35 | 1.26 | 0 | 0 | 0 | 0 |
| 46 | Taxonomic | lion | frog | 3.79 | 1.54 | 1.32 | 1.06 | 0 | 1 | 0 | 1 |
| 47 | Taxonomic | dress | shorts | 4.25 | 2.32 | 1.16 | 1.34 | 1 | 0 | 1 | 0 |
| 48 | Taxonomic | saw | screwdriver | 4.46 | 2.54 | 1.10 | 1.57 | 1 | 0 | 1 | 0 |
| 49 | Thematic | axe | wood | 1.46 | 4.56 | 1.05 | 0.92 | 1 | 0 | 1 | 1 |
| 50 | Thematic | tent | feather | 1.46 | 3.90 | 1.01 | 1.18 | 0 | 0 | 1 | 1 |
| 51 | Thematic | monkey | banana | 1.44 | 4.49 | 1.06 | 1.03 | 0 | 1 | 1 | 1 |
| 52 | Thematic | dress | girl | 1.75 | 4.00 | 1.27 | 1.10 | 1 | 0 | 0 | 1 |
| 53 | Thematic | tropicalfish | aquariumplant | 1.53 | 3.19 | 1.05 | 1.49 | 0 | 1 | 1 | 1 |
| 54 | Thematic | squirrel | hazelnuts | 1.65 | 4.53 | 1.19 | 1.02 | 0 | 1 | 1 | 1 |
| 55 | Thematic | couch | cushion | 2.28 | 4.17 | 1.37 | 1.11 | 0 | 0 | 1 | 0 |
| 56 | Thematic | camel | palmtree | 1.54 | 4.31 | 1.12 | 1.03 | 0 | 1 | 0 | 1 |
| 57 | Thematic | cow | grass | 1.53 | 3.97 | 1.06 | 1.27 | 0 | 1 | 1 | 1 |
| 58 | Thematic | saw | carpenter | 1.42 | 4.49 | 1.10 | 1.01 | 1 | 0 | 0 | 1 |
| 59 | Thematic | train | businessman | 1.32 | 3.76 | 0.92 | 1.19 | 0 | 0 | 0 | 1 |
| 60 | Thematic | car | trafficlight | 1.46 | 4.65 | 1.05 | 0.77 | 0 | 0 | 0 | 0 |
| 61 | Thematic | telescope | moon | 1.47 | 4.50 | 1.14 | 0.98 | 1 | 0 | 0 | 1 |
| 62 | Thematic | strawberry | jam | 1.91 | 4.44 | 1.34 | 0.92 | 1 | 1 | 1 | 0 |
| 63 | Thematic | nose | barbecue | 1.19 | 3.32 | 0.73 | 1.25 | 1 | 1 | 0 | 0 |
| 64 | Thematic | shirt | iron | 1.54 | 4.29 | 1.16 | 1.04 | 1 | 0 | 1 | 0 |
| 65 | Thematic | lion | cage | 1.64 | 3.85 | 1.34 | 1.35 | 0 | 1 | 0 | 0 |
| 66 | Thematic | bed | babysleeping | 1.46 | 4.64 | 1.12 | 0.94 | 0 | 0 | 0 | 1 |
| 67 | Thematic | ivy | pot | 1.41 | 4.19 | 1.02 | 1.16 | 1 | 1 | 1 | 0 |
| 68 | Thematic | sink | toothbrush | 1.76 | 4.35 | 1.24 | 1.00 | 0 | 0 | 1 | 0 |
| 69 | Thematic | castle | knight | 1.74 | 4.60 | 1.28 | 0.83 | 0 | 0 | 0 | 1 |
| 70 | Thematic | ear | phone | 1.37 | 4.32 | 0.93 | 1.05 | 1 | 1 | 1 | 0 |
| 71 | Thematic | hammer | nail | 2.58 | 4.46 | 1.42 | 1.16 | 1 | 0 | 1 | 0 |
| 72 | Thematic | cherries | basket | 1.31 | 3.90 | 0.89 | 1.18 | 1 | 1 | 1 | 0 |
| 73 | Thematic | truck | road | 1.44 | 4.56 | 0.98 | 0.98 | 0 | 0 | 0 | 0 |
| 74 | Thematic | caterpillar | leaf | 1.69 | 4.38 | 1.20 | 1.00 | 0 | 1 | 1 | 1 |
| 75 | Thematic | pipe | match | 1.76 | 4.40 | 1.24 | 0.97 | 1 | 0 | 1 | 0 |
| 76 | Thematic | rose | wateringcan | 1.57 | 4.29 | 1.25 | 1.08 | 1 | 1 | 1 | 0 |
| 77 | Thematic | carrot | rake | 1.34 | 3.56 | 0.88 | 1.31 | 1 | 1 | 1 | 0 |
| 78 | Thematic | razor | hair | 1.21 | 2.65 | 0.67 | 1.28 | 1 | 0 | 1 | 1 |
| 79 | Thematic | daisy | butterfly | 1.85 | 4.22 | 1.23 | 1.13 | 1 | 1 | 0 | 1 |

| | | | | | | | | | | | |
|---|---|---|---|---|---|---|---|---|---|---|---|
| 80 | Thematic | motorbike | bomberjacket | 1.28 | 3.11 | 0.77 | 1.36 | 0 | 0 | 1 | 0 |
| 81 | Thematic | apple | knife | 1.39 | 3.75 | 1.04 | 1.12 | 1 | 1 | 1 | 0 |
| 82 | Thematic | smallboat | oars | 2.07 | 4.62 | 1.39 | 0.97 | 0 | 0 | 1 | 0 |
| 83 | Thematic | scooter | helmet | 1.60 | 4.64 | 1.12 | 0.92 | 0 | 0 | 1 | 0 |
| 84 | Thematic | hoe | celery | 1.37 | 3.57 | 0.91 | 1.28 | 1 | 0 | 1 | 1 |
| 85 | Thematic | sled | snowman | 1.49 | 4.42 | 1.16 | 1.14 | 1 | 0 | 0 | 0 |
| 86 | Thematic | sheep | ballofyarn | 1.47 | 4.35 | 1.13 | 0.94 | 0 | 1 | 1 | 0 |
| 87 | Thematic | sneaker | football | 1.60 | 4.39 | 1.15 | 0.94 | 1 | 0 | 1 | 0 |
| 88 | Thematic | mouse | smallwheel | 1.40 | 4.24 | 0.99 | 1.17 | 0 | 1 | 1 | 0 |
| 89 | Thematic | mitten | skis | 1.92 | 4.31 | 1.34 | 1.07 | 1 | 0 | 1 | 0 |
| 90 | Thematic | sparrow | egg | 1.37 | 3.03 | 0.72 | 1.57 | 0 | 1 | 1 | 1 |
| 91 | Thematic | kite | lightning | 1.25 | 2.65 | 0.71 | 1.26 | 1 | 0 | 0 | 0 |
| 92 | Thematic | hand | ring | 1.39 | 4.50 | 1.04 | 0.98 | 1 | 1 | 1 | 0 |
| 93 | Thematic | Xmastree | gift | 1.45 | 4.51 | 1.05 | 0.92 | 1 | 1 | 1 | 0 |
| 94 | Thematic | dog | animaldish | 1.31 | 4.65 | 0.89 | 0.91 | 0 | 1 | 1 | 0 |
| 95 | Thematic | bee | fingerwithsting | 1.37 | 4.57 | 0.98 | 1.00 | 0 | 1 | 0 | 1 |
| 96 | Thematic | bucket | faucet | 1.67 | 4.11 | 1.15 | 1.09 | 1 | 0 | 1 | 0 |
| 97 | Unrelated | sled | giraffe | 1.00 | 1.01 | 0.00 | 0.12 | 1 | 0 | 0 | 1 |
| 98 | Unrelated | cherries | babybottle | 1.14 | 1.33 | 0.46 | 0.67 | 1 | 1 | 1 | 0 |
| 99 | Unrelated | castle | beetle | 1.01 | 1.21 | 0.12 | 0.50 | 0 | 0 | 0 | 1 |
| 100 | Unrelated | bucket | toaster | 1.71 | 1.26 | 1.08 | 0.65 | 1 | 0 | 1 | 0 |
| 101 | Unrelated | carrot | pacifier | 1.19 | 1.39 | 0.68 | 0.68 | 1 | 1 | 1 | 0 |
| 102 | Unrelated | hammer | washingmachine | 1.34 | 1.60 | 0.75 | 1.07 | 1 | 0 | 1 | 0 |
| 103 | Unrelated | scooter | lobster | 1.01 | 1.03 | 0.12 | 0.24 | 0 | 0 | 0 | 1 |
| 104 | Unrelated | sheep | peach | 1.16 | 1.19 | 0.58 | 0.60 | 0 | 1 | 1 | 1 |
| 105 | Unrelated | razor | mushroom | 1.00 | 1.04 | 0.00 | 0.20 | 1 | 0 | 1 | 1 |
| 106 | Unrelated | bed | sword | 1.06 | 1.11 | 0.29 | 0.43 | 0 | 0 | 1 | 0 |
| 107 | Unrelated | lion | sweater | 1.01 | 1.01 | 0.12 | 0.12 | 0 | 1 | 1 | 0 |
| 108 | Unrelated | sparrow | rocket | 1.36 | 2.19 | 0.86 | 1.22 | 0 | 1 | 0 | 0 |
| 109 | Unrelated | hand | pasture | 1.19 | 1.85 | 0.68 | 1.19 | 1 | 1 | 0 | 1 |
| 110 | Unrelated | mitten | tulip | 1.09 | 1.25 | 0.50 | 0.80 | 1 | 0 | 1 | 1 |
| 111 | Unrelated | pipe | jigsawpuzzle | 1.06 | 1.33 | 0.29 | 0.69 | 1 | 0 | 1 | 0 |
| 112 | Unrelated | kite | fox | 1.03 | 1.11 | 0.24 | 0.52 | 1 | 0 | 0 | 1 |
| 113 | Unrelated | apple | volcano | 1.04 | 1.06 | 0.21 | 0.29 | 1 | 1 | 0 | 1 |
| 114 | Unrelated | dog | orange | 1.15 | 1.78 | 0.50 | 1.38 | 0 | 1 | 1 | 1 |
| 115 | Unrelated | tent | onion | 1.00 | 1.08 | 0.00 | 0.33 | 0 | 0 | 1 | 1 |
| 116 | Unrelated | squirrel | tire | 1.03 | 1.12 | 0.24 | 0.50 | 0 | 1 | 1 | 0 |
| 117 | Unrelated | mouse | mountain | 1.06 | 1.14 | 0.24 | 0.42 | 0 | 1 | 0 | 1 |
| 118 | Unrelated | dress | crane | 1.00 | 1.03 | 0.00 | 0.17 | 1 | 0 | 1 | 0 |
| 119 | Unrelated | axe | wallet | 1.04 | 1.21 | 0.20 | 0.69 | 1 | 0 | 1 | 0 |
| 120 | Unrelated | train | shark | 1.00 | 1.14 | 0.00 | 0.59 | 0 | 0 | 0 | 1 |
| 121 | Unrelated | sneaker | walnut | 1.01 | 1.26 | 0.12 | 0.58 | 1 | 0 | 0 | 1 |

| | | | | | | | | | | | |
|---|---|---|---|---|---|---|---|---|---|---|---|
| 122 | Unrelated | bee | book | 1.01 | 1.36 | 0.12 | 0.84 | 0 | 1 | 1 | 0 |
| 123 | Unrelated | Xmastree | lemon | 1.66 | 1.31 | 1.08 | 0.70 | 1 | 1 | 1 | 1 |
| 124 | Unrelated | telescope | bone | 1.03 | 1.04 | 0.17 | 0.35 | 1 | 0 | 1 | 1 |
| 125 | Unrelated | smallboat | lightbulb | 1.04 | 1.12 | 0.26 | 0.47 | 0 | 0 | 1 | 0 |
| 126 | Unrelated | saw | horse | 1.00 | 1.10 | 0.00 | 0.42 | 1 | 0 | 0 | 1 |
| 127 | Unrelated | couch | dice | 1.17 | 1.62 | 0.45 | 0.97 | 0 | 0 | 1 | 0 |
| 128 | Unrelated | hoe | piano | 1.06 | 1.00 | 0.23 | 0.00 | 1 | 0 | 1 | 0 |
| 129 | Unrelated | daisy | lock | 1.00 | 1.08 | 0.00 | 0.40 | 1 | 1 | 1 | 0 |
| 130 | Unrelated | tropicalfish | camper | 1.01 | 1.36 | 0.12 | 0.81 | 0 | 1 | 0 | 0 |
| 131 | Unrelated | ear | garlic | 1.01 | 1.06 | 0.12 | 0.23 | 1 | 1 | 1 | 1 |
| 132 | Unrelated | car | stairs | 1.04 | 1.10 | 0.20 | 0.30 | 0 | 0 | 0 | 0 |
| 133 | Unrelated | monkey | star | 1.00 | 1.11 | 0.00 | 0.46 | 0 | 1 | 0 | 1 |
| 134 | Unrelated | sink | donkey | 1.01 | 1.07 | 0.12 | 0.48 | 0 | 0 | 0 | 1 |
| 135 | Unrelated | shirt | peanut | 1.04 | 1.07 | 0.27 | 0.31 | 1 | 0 | 1 | 1 |
| 136 | Unrelated | rose | tv | 1.04 | 1.11 | 0.20 | 0.36 | 1 | 1 | 1 | 0 |
| 137 | Unrelated | strawberry | fence | 1.13 | 1.93 | 0.44 | 1.18 | 1 | 1 | 0 | 0 |
| 138 | Unrelated | camel | treebranch | 1.20 | 1.82 | 0.58 | 1.03 | 0 | 1 | 1 | 1 |
| 139 | Unrelated | cow | glasses | 1.04 | 1.08 | 0.20 | 0.44 | 0 | 1 | 1 | 0 |
| 140 | Unrelated | motorbike | clothespin | 1.09 | 1.14 | 0.51 | 0.61 | 0 | 0 | 1 | 0 |
| 141 | Unrelated | truck | shell | 1.00 | 1.10 | 0.00 | 0.34 | 0 | 0 | 1 | 1 |
| 142 | Unrelated | caterpillar | artichoke | 1.38 | 2.42 | 0.88 | 1.32 | 0 | 1 | 1 | 1 |
| 143 | Unrelated | nose | church | 1.01 | 1.22 | 0.12 | 0.75 | 1 | 1 | 0 | 0 |
| 144 | Unrelated | ivy | comb | 1.01 | 1.04 | 0.12 | 0.26 | 1 | 1 | 1 | 0 |

##### Table S84. Category percentages per condition.

| | Probe Manipulable | Probe Natural | Target Manipulable | Target Natural |
|---|---|---|---|---|
| **Taxonomic** | 52.08 | 47.92 | 52.08 | 47.92 |
| **Thematic** | 52.08 | 47.92 | 68.75 | 39.58 |
| **Unrelated** | 52.08 | 47.92 | 64.58 | 50.00 |

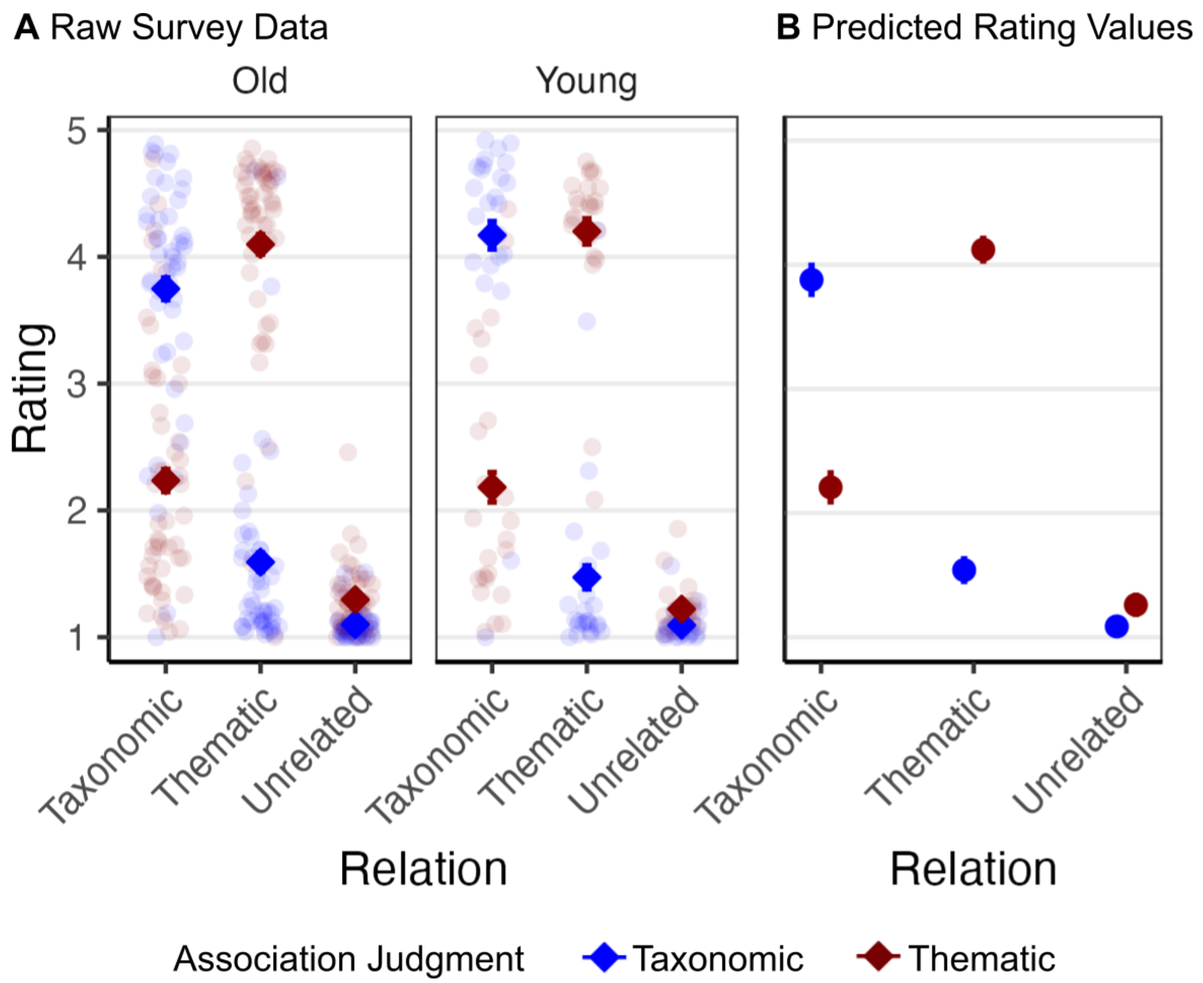

**Figure S10. Semantic relatedness ratings.** (A) Groupwise mean and individual ratings for all items acquired after the fMRI experiment. (B) Predicted rating values for the interaction of Relation and Judgment coming from the LMM: *Association_Rating ~ Relation + Association_judgment + Group + Relation:Association_judgment + (1 + Relation | Participant) + (1 | TargetItem).*

**Table S85. Results from the LMM for survey ratings.**

| | *Association Rating* | | | |
|---|---|---|---|---|
| *Predictors* | *Estimates* | *std. Error* | *Statistic* | *p* |
| *Relation [Taxonomic]* | *1.87* | *0.08* | *24.18* | ***2.36e-127*** |
| *Relation [Thematic]* | *1.66* | *0.07* | *24.29* | ***2.03e-128*** |
| *Association judgment [Thematic]* | *0.36* | *0.01* | *25.66* | ***9.37e-143*** |
| *Group [Older adults]* | *0.06* | *0.04* | *1.61* | *1.06e-01* |
| *Relation [Taxonomic] × Association judgment [Thematic]* | *-1.85* | *0.03* | *-53.53* | ***0.00e+00*** |
| *Relation [Thematic] × Association judgment [Thematic]* | *2.41* | *0.03* | *69.86* | ***0.00e+00*** |
| *ICC* | *0.16* | | | |
| *N Participant* | *72* | | | |
| *N TargetItem* | *144* | | | |
| *Marginal $R^2$ / Conditional $R^2$* | *0.552 / 0.623* | | | |

**Table S86. Results from paired comparisons for survey ratings.**

| *Contrast* | *Estimate* | *SE* | *z ratio* | *p value* |
|---|---|---|---|---|
| Unrelated Taxonomic - Taxonomic Taxonomic | -2.79 | 0.08 | -35.24 | <0.001 |
| Unrelated Taxonomic - Thematic Taxonomic | -0.45 | 0.07 | -6.45 | <0.001 |
| Unrelated Taxonomic - Unrelated Thematic | -0.17 | 0.02 | -7.12 | <0.001 |
| Unrelated Taxonomic - Taxonomic Thematic | -1.12 | 0.08 | -14.14 | <0.001 |
| Unrelated Taxonomic - Thematic Thematic | -3.03 | 0.07 | -43.12 | <0.001 |
| Taxonomic Taxonomic - Thematic Taxonomic | 2.34 | 0.08 | 29.02 | <0.001 |
| Taxonomic Taxonomic - Unrelated Thematic | 2.62 | 0.08 | 33.06 | <0.001 |
| Taxonomic Taxonomic - Taxonomic Thematic | 1.67 | 0.02 | 68.63 | <0.001 |
| Taxonomic Taxonomic - Thematic Thematic | -0.24 | 0.08 | -3.02 | 0.0382 |
| Thematic Taxonomic - Unrelated Thematic | 0.28 | 0.07 | 3.98 | 0.0010 |
| Thematic Taxonomic - Taxonomic Thematic | -0.67 | 0.08 | -8.27 | <0.001 |
| Thematic Taxonomic - Thematic Thematic | -2.58 | 0.02 | -106.04 | <0.001 |
| Unrelated Thematic - Taxonomic Thematic | -0.95 | 0.08 | -11.96 | <0.001 |
| Unrelated Thematic - Thematic Thematic | -2.86 | 0.07 | -40.68 | <0.001 |
| Taxonomic Thematic - Thematic Thematic | -1.91 | 0.08 | -23.78 | <0.001 |

*Note:* P-values are Bonferroni-corrected.
